# Continuous sensorimotor transformation enhances robustness of neural dynamics to perturbation in macaque motor cortex

**DOI:** 10.1101/2024.11.10.622879

**Authors:** Cong Zheng, Qifan Wang, He Cui

**Affiliations:** Institute of Neuroscience, Center for Excellence in Brain Science and Intelligence Technology, Chinese Academy of Sciences, Shanghai 200031, China; Chinese Institute for Brain Research, Beijing 102206, China

## Abstract

Neural activity in the motor cortex dynamically evolves to plan and generate movement. How motor cortex adapts to dynamic environments or perturbations remains to be fully explored. In this study, we investigated whether dynamic nature of targets in a reach task requires distinct preparatory dynamics in motor cortex and result in varying levels of robustness against disruptions. Two monkeys were trained to perform delayed center-out reaches either to a static target (static condition) or a rotating target that needed to be intercepted (moving condition). Despite nearly identical hand kinematics in both two conditions, responses to the perturbation of intracortical microstimulation (ICMS) differed. In the static condition, ICMS led to prolonged reaction times, particularly when delivered shortly before movement onset and at anterior sites in PMd, aligning with previous findings. Unexpectedly, ICMS reduced reaction times in the moving condition. Furthermore, neural firing rates differed between the static and the moving conditions, with population activity in the latter exhibiting more rapid changes post-perturbation. Spatio-temporal sensorimotor transformation dominated throughout the preparation in the moving condition, while the static condition showed less stable motor intention representation, particularly during the late delay period. An input-driven model replicated the differences in RT-prolonging effect by assuming distinct input control strategy for the static and the moving condition. These findings suggest that input from a moving target to motor cortex can counteract ICMS effects, enabling the motor network to generate appropriate commands more quickly. Lastly, we propose that ICMS may facilitate go cue recognition, providing a potential explanation for the shortened reaction times in the moving condition.

## Introduction

As Albert Einstein once said, ‘*Life is like riding a bicycle. To keep your balance, you must keep moving*’^1^, this metaphor illustrates the principle that motion, rather than remaining still, enables stability and resilience in the face of external disturbances. It is true not only because actions, or movements are the only valid ways we connect to the world^2^, but also in that the neural population functions as a dynamical system that continuously evolves over time, performing computations to generate movement^3–6^. In such process, driven by intrinsic motor cortical dynamics and inputs from other brain regions, preparatory activity sets the neural state into an optimal initial condition, seeding the near-autonomous ‘rotational dynamics’ of cortical states during movement^7–10^. Preparatory activity is acknowledged as a key link in the causal chain that generates voluntary movement, yet several paradoxes remain unsolved. Disrupting the preparatory states with intracortical microstimulation (ICMS) leads to an increase in reaction times (RTs), supporting the hypothesis that movement preparation is a time-consuming optimization process and that movement is delayed if errors are still present^11^. However, subsequent studies suggest that the preparatory activity could be bypassed or can occur unexpectedly rapidly if needed, raising questions of the nature of the delay-period activity^12,13^.

Multiple studies have revealed the conserved preparatory activity across contexts^13,14^, echoing the experimental findings that ICMS induces similarly stereotyped RT responses regardless of context^15^. While the commonality of preparatory activity is recognized, the role of differences in neural dynamics in adapting to distinct contexts, particularly dynamic environments, remains unclear. Interception is a context of reaching requiring substantial interaction with a dynamically changing environment. Successful interception requires ongoing predictions and updates of sensorimotor states^16–22^. Recently, we showed that target motion significantly affects neural activity in the macaque motor cortex during interception preparation^23^. The continuous sensorimotor transformation involves multiple brain regions, especially the posterior parietal cortex^24–26^, which continuously sends external inputs to motor cortex^23,27–31^. Although this dynamic context appears to differ from context for reaches to static targets, direct evidence elucidating whether preparatory dynamics differ between these contexts and how such differences contribute to computations is still lacking.

Recent studies highlight the importance of incorporating external inputs to fully understand intrinsic neural dynamics^32–34^. Utilizing an innovative perturbation design, researchers dissected the roles of local dynamics and inputs in cortical pattern generation, demonstrating that motor generation is driven by the inputs from motor thalamus^33^. Modeling works further indicate that optimal feedback inputs are critical in optimizing motor cortical neural states to enable rapid motor preparation^35^, and that optimal control inputs can autonomously seed preparatory states for movement generation^36^. Given the sensorimotor information that feeds to the motor cortex during interception, it is intriguing to explore whether and how these inputs shape neural dynamics.

The present study addressed the important question of whether distinct preparatory dynamics in the motor cortex generates different degrees of robustness to perturbations of these dynamics. We trained two monkeys to perform delayed center-out reaches toward either a static target (static condition) or intercepting reaches toward a rotating target (moving condition). Monkeys showed similar reach trajectories in the static and the moving conditions. However, ICMS during the late delay selectively prolonged RTs in the static but not in the moving condition. Neural states post-ICMS diverged more from intact states in the static condition, with the temporal shift between non-stimulated and stimulated states predicting RT changes. For moving targets, target or reach direction representations adapted with motion, remaining stable throughout the delay, while the static target representations were less stable, displaying two distinct preparatory phases. Using an input-driven neural network model, we replicated the differential ICMS effects on RTs, showing that optimal feedback from other regions can quickly correct perturbation-induced errors in the moving condition. These findings reveal the differential ICMS effects on distinct neural dynamics and highlight the crucial role of continuous sensorimotor transformation in resilience to unexpected perturbation.

## Results

### Reach kinematics of the static and moving conditions

We trained two monkeys (G and L) to perform delayed center-out reaches either to a static target (static condition) or a rotating target that needed to be intercepted (moving condition) ^16^. Interception requires predicting future reach directions based on a model of target motion and one’s kinematics (**Fig. 1a**). In a typical session, target velocities were either 0°/s (static) or clockwise 120°/s (−120°/s, moving), with delay periods of 200 ms (short, s) or 900 ms (long, l) (**Fig. 1b**). The initial locations for the moving targets were designed to encourage reach towards four evenly distributed directions. These designed trials were interleaved with trials featuring random initial target locations, comprising 40-60% of the total trials to maintain randomness and prevent the prediction of reach direction (**Fig. 1c**). We required the reach endpoints to fall within 2.8 cm of the targets, a criterion that both monkeys met post-training. Endpoint distributions were direction-specific, consistently lagging behind or landing ahead of the target in both target velocity conditions, with minimal variances, indicating habituated reach strategies (**Fig. 1d**).

**Fig. 1.**
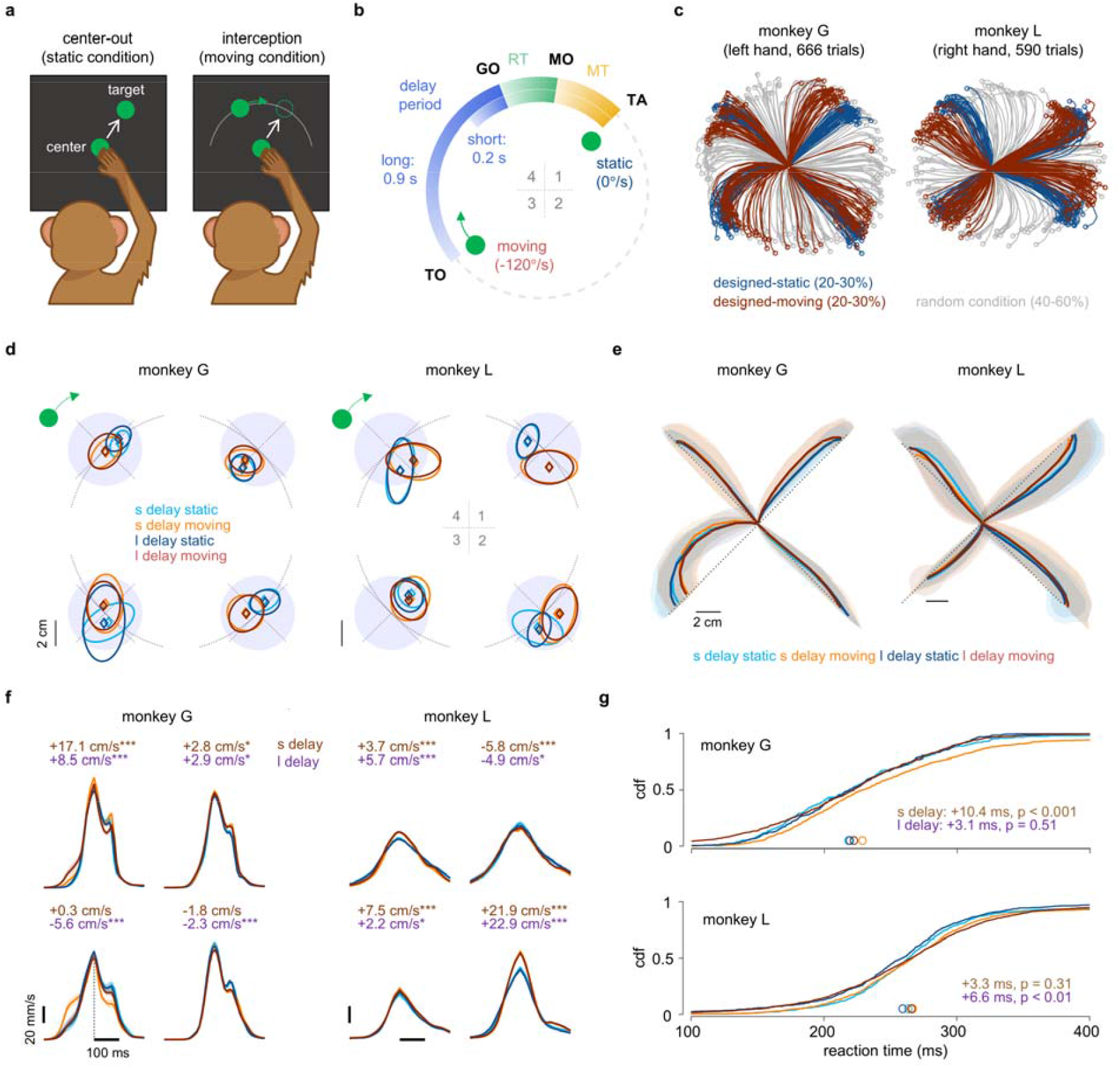
Reach kinematics of the static and moving conditions. **a**. Task description. Monkeys performed two types of reach tasks: reaching for a static target or intercepting a moving target. **b**.Task structure. Target onset (TO) is followed by a delay of either 200 ms (short, s) or 900 ms (long, l). Dimmed center green dot indicates go cue. Movement onsets (MO) and target acquirement (TA) delineate reaction times (RTs) and movement times (MTs). **c**.Reach trajectories from example sessions. Designed static and moving conditions share similar initial target location for statistical comparison, while the random condition features random initial target locations to avoid predictability. **d**.Reach endpoints relative to target locations (mean ± 95%CI). Subplots correspond to four reach directions. Dotted black lines indicate tangential and radial directions related to target location. Purple shadings denote the error tolerance range. **e**.Reach trajectories aligned to the corresponding directions (mean ± 68%CI). **f**.Hand speed profiles aligned to the peak speed (mean ± 95%CI). Differences in peak velocity (v_moving_ – v_static_) are noted. The number of trials and annotations are consistent with panel **d**, with significance indicated by asterisks based on Wilcoxon rank sum test: *p < 0.05, ***p < 0.001. **g**.Empirical cumulative distribution function (CDF) for RTs. Colored circles indicate the median RT. The difference in RT (RT_moving_ – RT_static_) is noted, with annotations matching those in panel **f**.

Reach trajectories were comparable between the static and moving conditions. When aligned to the same directions, similar curving patterns emerged in two conditions in four reach directions (**Fig, 1e**). Analyses of launching angles revealed minor differences between the static and moving conditions, suggesting that the curvature of these trajectories may be influenced by gravitational or biomechanical constraints, as this pattern was also observed in the static condition (**Supplementary Fig. 1b**). Hand peak speeds were generally higher in the moving condition, though the difference varied by reach directions (**Fig. 1f**). Reaction times (RTs) were typically longer in the moving condition (**Fig. 1g**). A longer delay period resulted in decreased RTs for the moving condition in monkey G and for the static condition in monkey L, indicating that extended preparation time facilitates better motor planning, consistent with the previous study ^37–39^. Some trials showed reaction times below 150 ms, which is expected since the moving targets can elicit fast, reactive intercepting movements^13,40,41^. Our analyses indicate that 35.3% (short delay) and 52.2% (long delay) of trials with RTs below 150 ms were from the designed conditions for monkey G, while these proportions were 42.1% (short delay) and 52.9% (long delay) for monkey L. This suggests that shorter RTs reflect biological variability rather than anticipation of the go cue (GO). Overall, with adequate preparation time, monkeys exhibited well-prepared reaches even when the target was moving, suggesting they relied on predictions of future reach directions during interception rather than targeting the instantaneous target position; otherwise, the reach trajectory would have shown significant curved ^22^.

### Perturbation delays initiation of reaches directed at the static but not the moving targets

Experimental perturbations, such as intracortical microstimulation (ICMS), can provide insights into neural dynamics^11,15,42,43^. Following the methodology established by Churchland and Shenoy (2007)^11^, we applied sub-threshold ICMS (100 ms at 300 Hz) to either primary motor cortex (M1) or dorsal premotor cortex (PMd) using a single electrode, while simultaneously recording nearby neural activity with a 64-channel linear probe (**Fig. 2a**). ICMS amplitudes were set 5-10 μA below threshold, which was determined by gradually increasing the stimulation until a muscle twitch or movement occurred, then reducing the amplitude until no movement was detected.

**Fig. 2.**
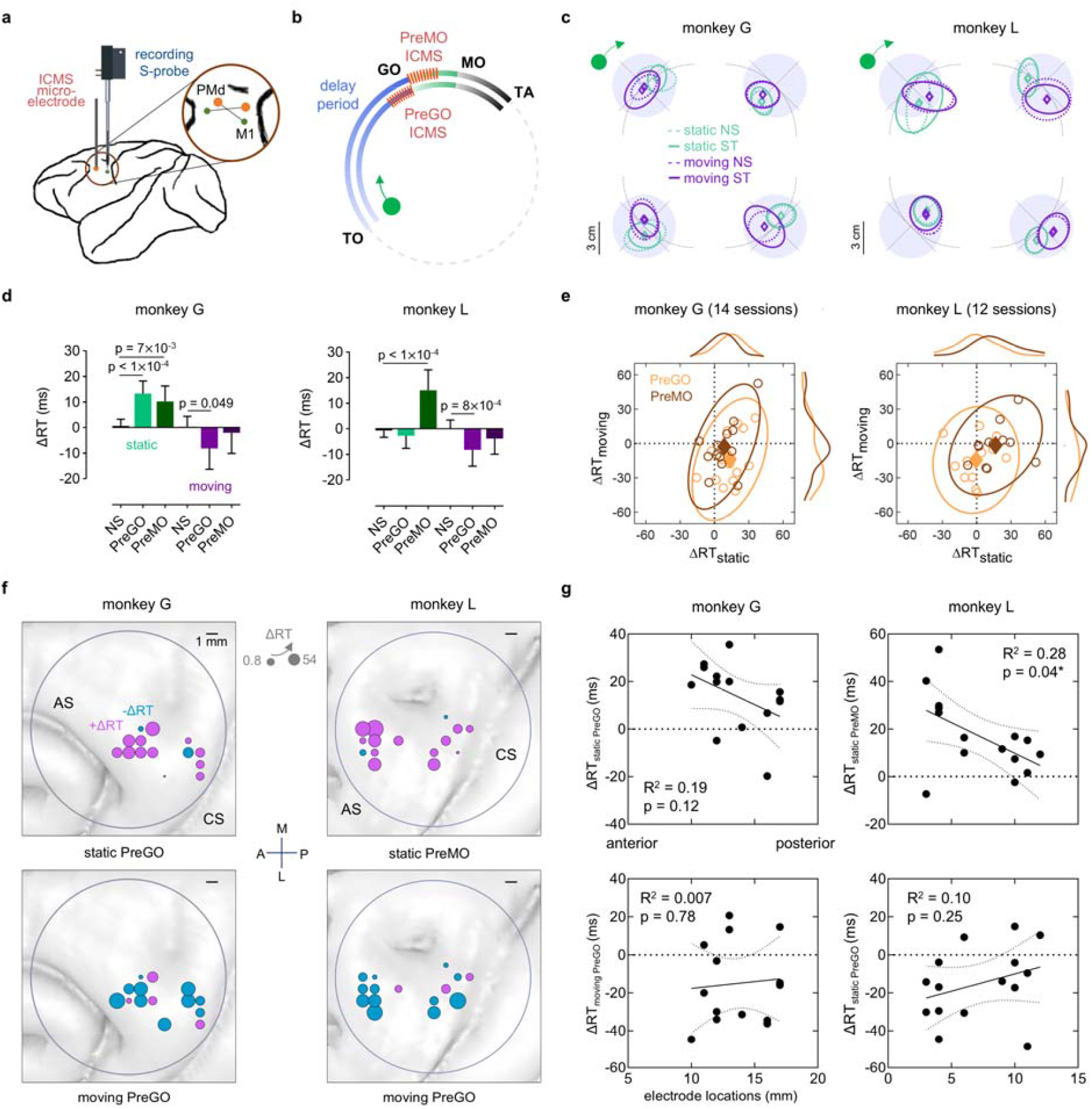
ICMS increases reaction times in the static condition but not in the moving condition. **a**.Microstimulation and recording setup. Inset illustrate the electrode pairs used for stimulation (orange) and recording (green) in either PMd or M1. Also see **Supplementary Fig. 2d**. **b**.Timing of microstimulation in the long-delay condition. ICMS is delivered at either PreGO (100-0 ms before Go cue) or PreMO (0-150 ms after go cue). The timing for the short-delay condition remains consistent with this. **c**.Reach endpoints relative to target locations (mean + 95%CI) after ICMS. Annotations are consistent with those in Fig. **1d**. **d**.Changes in reaction times (median + 95%CI) calculated as ΔRT = RT_ST_ – RT_matched NS_ averaged across reach directions in the long-delay condition. Statistical significance is assessed using Wilcoxon rank sum test. **e**.Individual session ΔRTs. Circles indicate median ΔRT of individual sessions, accompanied by marginal histograms. Filled diamonds and ellipses indicate means and 95% confidence range of PreGO and PreMO ICMS conditions. **f**.Location-dependent ICMS effect on reaction times. Dot size represents the magnitude of ΔRT, with the minimum and maximum sizes indicated in the inset. Colors denote the sign of ΔRT (cyan for negative, purple for positive). **g**.ΔRT as a function of anterior-posterior location. Electrode locations range from 0 (anterior, PMd) to 20 (posterior, M1). Dotted curves indicate the 95% CIs for a linear fit. Goodness of fit (*R*^*2*^) and p-value of non-zero slope are provided.

Electromyography (EMG) analyses indicated that ICMS rarely elicited muscle activity (**Supplementary Fig. 2a**). Since critical preparatory neural dynamics primarily occur during the late delay period, ICMS was timed around go cue (PreGO, from GO-100 ms to GO) or just before movement onset (PreMO, from GO to GO+150 ms) (Fig. 2b). Trials with stimulation (stimulated, ST) and without stimulation (non-stimulated, NS) were randomly interleaved. In the following analyses, we primarily focused on the effects of ICMS in the long-delay condition, although ICMS was also applied in the short-delay condition (**Supplementary Figure 7, 8**).

The impact of ICMS on reach accuracy or movement kinematics, such as hand speed, was minimal (**Fig. 2c, Supplementary Fig. 7, 8**). However, ICMS produced contrasting effects on RTs: it prolonged RTs in the static condition while shortening RTs in the moving condition (**Fig. 2d**). We calculated changes in RTs (ΔRT = RT_ST_ – RT_matched NS_) and found that PreGO ICMS prolonged RTs in the static condition by 13.3 ms (95% CI: [7.5, 18.2]) for monkey G. PreMO ICMS increased RTs by 10.3 ms (95% CI: [3.6, 16.3]) for monkey G and 15.0 ms (95% CI: [8.0, 23.1]) for monkey L. This aligns with previous findings in delayed manual center-out task^11,42^. For monkey L, significant RT prolongation was observed only in the PreMO ICMS condition. Given monkey L’s generally longer RTs (**Fig. 1g**), PreMO ICMS timing may be closer to the critical preparatory epoch just before movement onset. In contrast, PreGO ICMS shortened RTs in the moving condition by 8.2 ms (95% CI: [−16.3, 1.3]) for monkey G and 8.2 ms (95% CI: [−14.6, −1.5]) for monkey L. The earlier timing of PreGO ICMS likely allowed for a greater reduction in RTs compared to PreMO ICMS. We also examined the averaged ΔRT across individual sessions, and the distributions of ΔRT confirmed that most sessions experienced increased RTs in the static condition and decreased RTs in the moving condition after ICMS (**Fig. 2e**).

The ICMS effect on RTs was found to be location-dependent (**Fig. 2f, g**). Specifically, stimulating of more anterior sites, such as PMd, resulted in a more pronounced RT-prolonging effect in the static condition, consistent with a previous study suggesting a close correlation between PMd activity and motor preparation^11^. Although there was a trend indicating that the RT-shortening effect in the moving condition was more evident in anterior areas, the relationship between antero-posterior location and RT differences was not statistically significant. This suggests that while the RT-shortening effect in the moving condition may be associated with preparatory activity, its underlying mechanisms could differ from those of RT-prolonging effect observed in the static condition. In addition, no significant correlations were found between ΔRT and ICMS amplitudes (**Supplementary Fig. 2c**), indicating that the location-dependent effect is less likely due to higher ICMS thresholds in PMd (**Supplementary Fig. 2b**). To further validate these findings, we conducted a series of control analyses, including bootstrapping method to compute ΔRT, and analyses to account for potential impacts from trial order and the randomness of conditions (**Supplementary Fig. 3-5**)

**Fig. 3.**
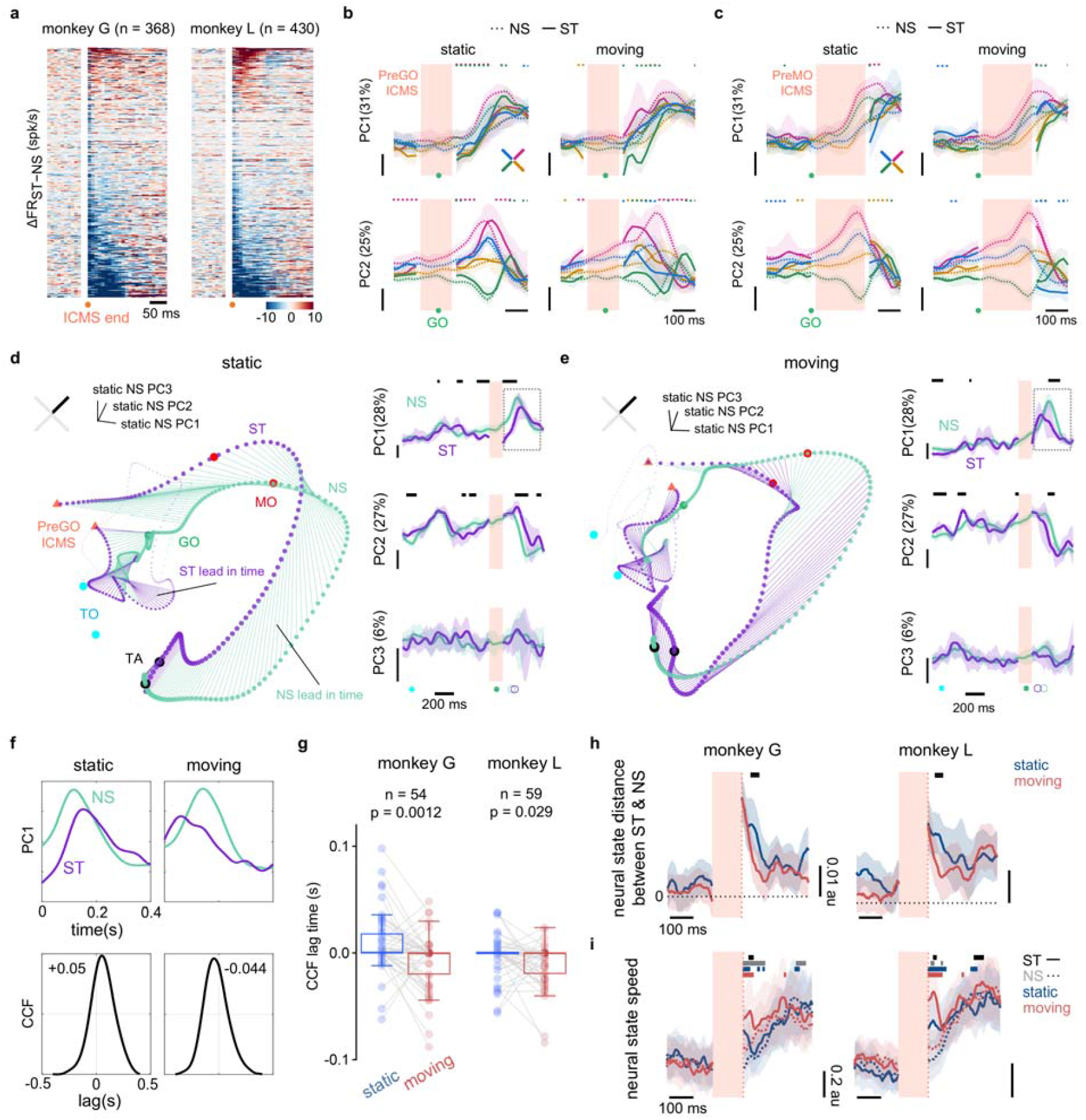
Faster recovery from perturbed states in the moving condition. **a**.Neuronal responses post-ICMS. Changes in firing rates (ΔFR = FR _ST_– FR _matched NS_) aligned to end of ICMS, with ICMS-period activity removed. Units are sorted by ΔFR within a 100 ms post-ICMS window. **b.c**. First two principal components (PCs) (mean ± 95%CI) of neural state space built from condition-averaged NS data in the static condition of an example session (56 neurons). Orange shades indicate the range of PreGO ICMS (**b**) or PreMO ICMS (**c**) period. ST data during ICMS is omitted. Colored dots denote time points with significantly differences between NS and ST trials (Wilcoxon rank sum test, p < 0.05). **d. e**. Neural trajectories (right) and the first three PCs (mean ± 95%CI) (left) of a representative static condition (**d**) or moving condition (**e**) in the same state space as in panel **b**. Left: orange triangles mark ICMS start and end, with ICMS-period activity omitted. Colored lines connecting NS and ST conditions at each time point indicate which condition is relatively ahead along the trajectory. Right: black lines indicate time intervals of significant differences between two conditions (Wilcoxon rank sum test, p < 0.05). **f**. Example cross-correlation function (CCF) for PC1. Top: PC1 of time epoch shown in panel **d** and **e** (dashed box). Bottom: CCF between NS and ST conditions. The lag time, where the CCF peaks, is noted in seconds. **g**. Pooled CCF lag times across reach directions and sessions. Statistical difference is assessed using Wilcoxon matched-pairs signed rank test. **h**. Neural state distance (mean ± 95%CI) between NS and ST conditions. Data are aligned to end of ICMS and averaged across reach directions and sessions (monkey G: n = 52; monkey L: n = 60). Orange shading marks the ICMS duration. Black lines indicate time intervals with significant differences (Wilcoxon rank sum test, p < 0.05). **i**. Neural state speed (mean ± 95%CI). Data are aligned to end of ICMS and averaged (monkey G: n = 28; monkey L: n = 48). Colored lines above indicate time intervals with significant differences (Wilcoxon rank sum test, p < 0.05) between the static and moving condition at NS (black) and ST (gray), as well as difference between NS and ST in the static (blue) and the moving condition (red). Other annotations follow panel h.

### Faster recovery from perturbed states in the moving condition

To investigate the mechanisms underlying ICMS-induced RT changes, we analyzed the neural activity recorded concurrently with ICMS. We identified a total 368 and 430 single units for monkey G and L, respectively. Due to potential ICMS artifacts, we restricted our analyses to periods outside the ICMS window, setting neural data within this window to infinite or zero and excluding it from displayed results. Post-ICMS single-unit responses predominantly exhibited either inhibitory or excitatory patterns (**Fig. 3a**). Previous studies indicate that higher ICMS currents typically lead to longer inhibition periods, with durations increasing exponentially beyond a 40 μA threshold^44^. In our sessions, mean ICMS amplitudes were 67.5 ± 22.1 μA for monkey G and 34.7 ± 11.4 μA for monkey L, potentially explaining the greater proportion of inhibitory responses observed. We also examined the impact of proximity to stimulating sites on ICMS responses, finding a modest but significant positive correlation in monkey L’s data (**Supplementary Fig. 1h**), suggesting that units located farther from the stimulation site exhibit higher rates of excitation, consistent with previous findings^45^.

We next applied principal component analysis (PCA) to analyze neural population activity. The low-dimensional subspace was constructed using the condition-averaged NS trials in the static condition, which served as a base for projecting other conditions. The first principal component (PC1) was condition-invariant and reflected movement time, while the PC2 varied according to reach direction (**Fig. 3b, c**), consistent with previous studies ^46^. Following ICMS, differences between NS and ST conditions emerged in both the timing and condition-dependent components. In 3D subspace (explaining over 60% of the variance), neural trajectories for NS and ST initially evolved parallel but diverged after ICMS, with ST trajectories lagging in the static condition and leading in the moving condition (**Fig. 3d, e, Supplementary Fig. 9**). Given the established association between the advancement of neural states at the go cue and RTs^47,48^, we performed cross-correlation function (CCF) on PC1 to estimate the lag time between NS and ST conditions (**Fig. 3f**). Results indicated that the average lag times were positive in the static condition but negative in the moving condition, corresponding with the observed RT changes post-ICMS (**Fig. 3g**).

To further examine difference between NS and ST trials, we computed the neural state distance between all-dimensional neural trajectories using unbiased estimation approach^49^. Results indicated a greater neural distance following ICMS in the static condition, suggesting that ICMS induced more pronounced deviations in neural states from intact states in the static condition (**Fig. 3h**). To explore mechanisms underlying these differences, we analyzed the evolving speed of neural states (**Fig. 3i**). Post-ICMS neural state speeds were higher in the moving condition than in the static condition across both ST and NS trials, although neural speed in the static condition also increased by the end of ICMS. This finding suggests that the smaller neural distances in the moving condition may result from quicker recovery from ICMS-induced deviations. Overall, the prolonged RTs observed in the static condition post-ICMS align with a lag in neural states, likely due to deviations that require additional time to recover. In contrast, neural states in the moving condition are less disrupted and even exhibit accelerated dynamics, likely because the faster evolving speed of neural states reduces recovery time.

Previous study proposed that ICMS-induced changes in RTs can be interpreted within a framework where the brain delays movement initiation until preparatory errors are minimized, with larger errors resulting in longer RTs^11^. Deviations in neural states following ICMS may represent these preparatory errors, and recovery from such states would correspond to error resolution. The faster these errors are corrected, the quicker motor preparation completes, enabling timely movement execution. Our findings support this framework and raise an important question: do the differences in ICMS responses stem from intrinsic variations in preparatory neural dynamics between the static and moving conditions, despite their similar reach kinematics?

### The static and moving conditions differ at the single-unit level

To explore the potential different neural activity between the static and moving conditions, we firstly conducted analyses on single neural activity. The majority of sorted single units displayed activations during either delay period or movement (**Fig. 4a**); however, the proportion of neurons with pronounce pre-movement or preparatory responses was relatively lower in monkey L, likely due to sampling bias. While single-neuron activity exhibited reach-direction tunings, this tuning was more complex and sustained in the moving condition (**Fig. 4b**). For instance, example unit 1 maintained consistent directional tuning throughout the delay period in the static condition, whereas its tuning varied in the moving condition. Unit 2 and 3 tuned to reach directions only around TO and MO in the static condition, but exhibited intricate directional tuning throughout the entire preparatory epoch in the moving condition. Unit 4 did not show directional tuning until MO in the static condition, but began encoding reach-direction information midway through the delay period in the moving condition. To quantify the above findings, we calculated the number of time bins with significant differences in firing rates across reach directions, summed across all units, and observed that the static condition had larger proportion of directional tuning around TO and during movement, whereas the moving condition showed steadily increased and more directional tuning during late delay (**Fig. 4c**), suggesting distinct tuning dynamics in two conditions.

**Fig. 4.**
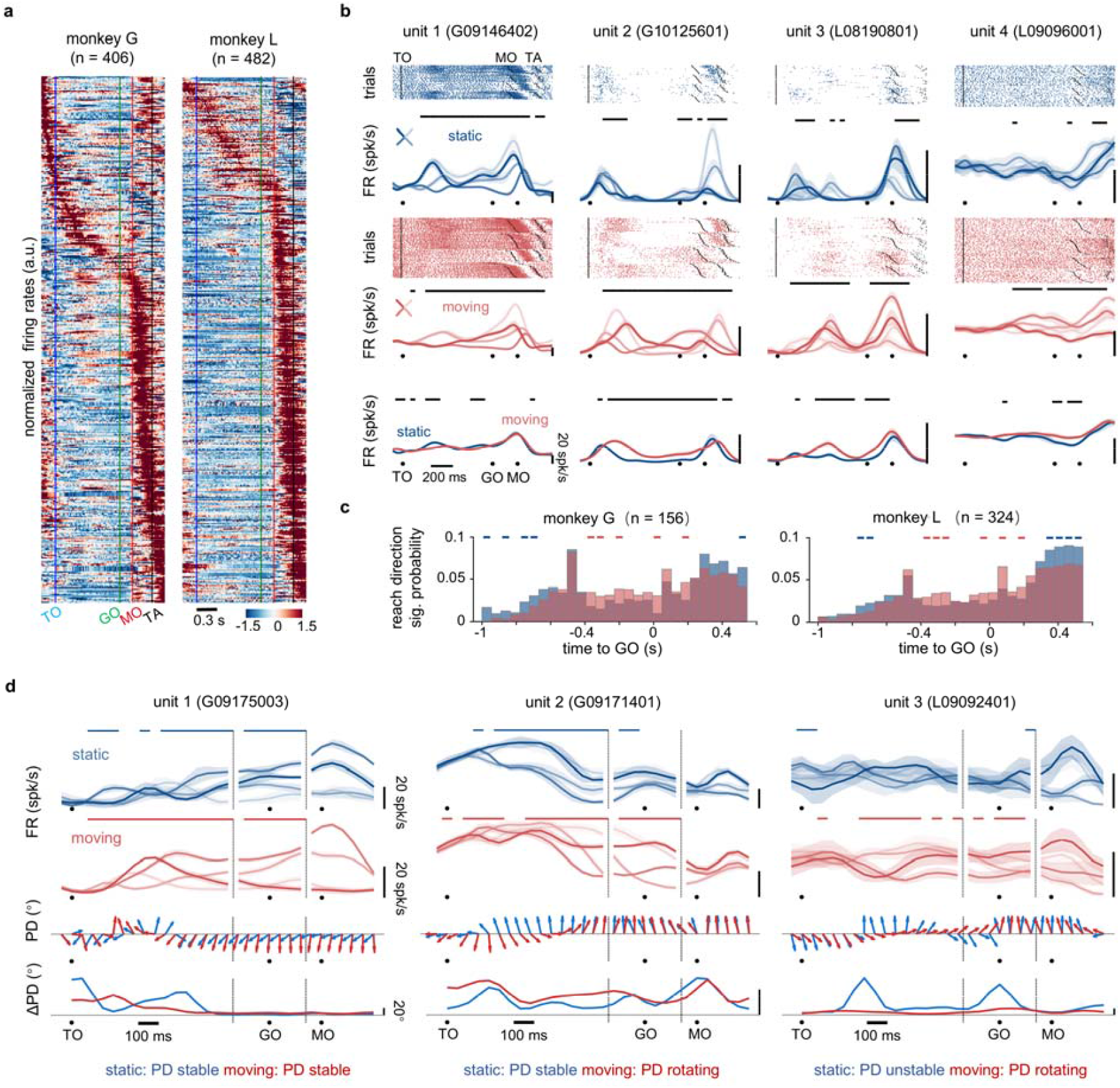
The static and moving conditions differ at the single-unit level. **a**. Heatmap of normalized firing rates for single units. Neural activity in the long-delay condition is aligned to Go cue, with units sorted by the time of peak firing rates. **b**. Example neural spike rasters and peri-event time histograms (PETHs; mean ± SEM). Black lines above indicate time intervals with significances (p < 0.05) based on Kruskal-Wallis test (top and middle) and Wilcoxon rank sum test (bottom). **c**. Normalized histogram showing the distribution of time bins where firing rates differs significantly (Kruskal-Wallis test, p < 0.05) across reach directions, summed across units. Colored lines: Fisher’s exact test p < 0.05. **d**. Concatenated PETHs (upper), preferred direction (PD, middle) and changes in preferred direction (Δ*PD* = PD_t_ − PD_t-1_, bottom) for example neurons. Each unit represents one type of PD change pattern.

We further examined the preferred directions (PDs) of individual neurons and found varying degrees of PD shifts, consistent with prior studies^23^. These PD shits indicate distinct computational processes that neurons undergo during motor preparation. Although the PD shift patterns were obscure for most neurons, three patterns are identified: (1) neurons with PD unchanged in both conditions (12/53), (2) neurons with PD stable in the static condition but continuously rotating in the moving condition (17/53), and (3) neurons with PD differing between early and late delay period in the static condition, buts rotating in the moving condition (24/53) (**Fig. 4d, Supplementary Fig. 10**). The first group are neurons with stable representation, served as a steady readout for motor execution. The second group of neurons likely prefers visual information, as their PD shifts are closely tied to target velocity. In contrast, the third subgroup shows unstable PD in the static condition, explaining decreased directional tuning at late delay period (**Fig. 4c**). Notably, we did not identify a certain group of neurons that exhibited PDs unstable in the moving condition while maintaining stable in the static condition. This suggests that the static condition possesses more heterogeneous neural activity, with a group of neurons maintaining sensorimotor information and another shifting to different neural computations, whereas neurons in the moving condition exhibited dynamic but stable representation of visual and motor information.

### Continuous sensorimotor transformation during preparation in the moving condition

Given the intriguing single-neuron activity, we further sought to explore the neural population dynamics. Using PCA to visualize low-dimensional neural population activity, we observed that the primary difference in neural trajectories between the static and moving conditions occurred during the late delay period (**Fig. 5a**). Although both conditions evolved roughly in parallel, the neural trajectories in the static condition twisted significantly around GO, whereas those in the moving condition were relatively short and straight. To quantify the geometry of these trajectories, we computed the detour index, defined as the ratio of the actual trajectory length to the shortest Euclidean distance between two neural states. We found that detour index was higher in the static condition, indicating a more elongated path of neural trajectories compared to the moving condition (**Fig. 5b**). This finding aligns with our single-unit observations (**Fig. 4d, e**), suggesting that neurons engage in different computations during the early or late delay period in the static condition, while their activities remain more homogenous in the moving condition.

**Fig. 5.**
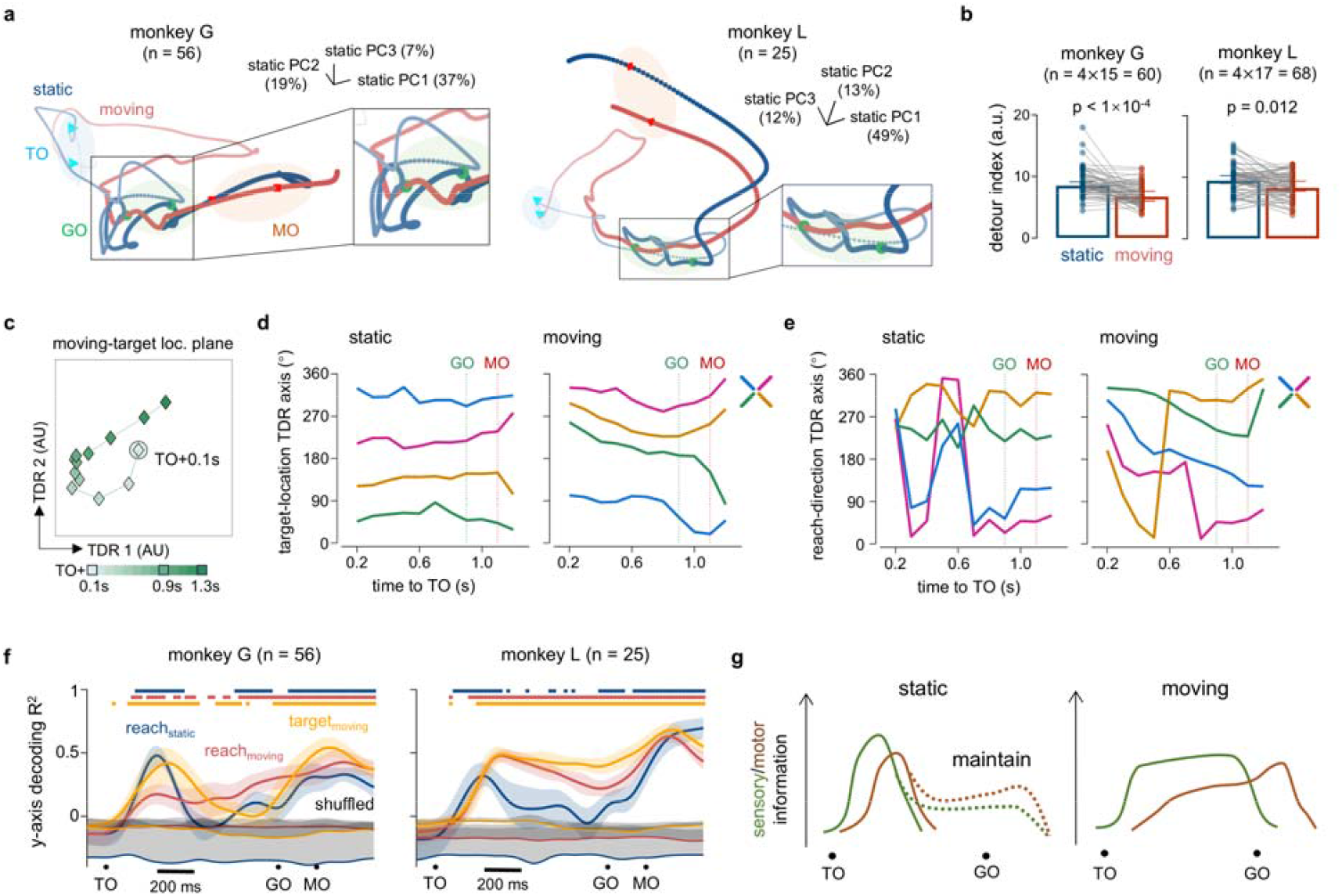
Continuous sensorimotor transformation during preparation in the moving condition. **a**. Neural trajectories for one example reach direction in the subspace constructed from the static condition. Colored filled circles mark events. Insets provide enlarged views of neural trajectories around the GO. **b**. Detour index for each reach direction across sessions. A higher detour index suggests longer neural trajectories during late delay period. Differences between conditions are assessed were assessed using Wilcoxon matched-pairs signed rank test. **c**. Neural states in the moving condition over time in target-location TDR subspace, constructed from neural activity of the moving condition at TO+0.1 s (large filled circles). One representative reach direction (left-down) is shown. Neural states evolved clockwise after TO. Data are averaged across seven sessions of monkey G with 181 neurons. **d**. Temporal changes in neural states relative to TO + 0.1 s within the static (left) and moving (right) target-location TDR subspace. Neural states in the moving condition evolved clockwise, while those in the static condition remained unchanged after TO. Data are the same as in panel **c**. **e**. Time course of absolute change in neural state position in the static (left) and moving (right) target-location TDR subspace. Neural states in both conditions converged to similar relative positions by MO. Data are the same as in panel **c**. **f**. Goodness of fit (*R*^*2*^, mean ± 95%CI) for linear decoding of reach directions (y-axis components) in the static (reach_static_) and moving (reach_moving_) condition, as well as instantaneous target location in the moving condition (target_moving_), based on neural activity in example datasets. Colored lines above denote intervals with significant differences from the shuffled baseline (Wilcoxon rank sum test, p < 0.05). **g**. Schematic of sensorimotor transformation in motor cortex. In the static condition, the transformation is completed shortly after target onset, with the motor plan maintained locally or elsewhere (dashed lines). In the moving condition, sensory information is continuously updated, and motor intention gradually forms until GO, when movement is initiated.

Next, we performed targeted dimensionality reduction (TDR) on the condition-averaged neural activity that pooled across sessions, aiming to identify a low-dimensional subspace that captures variance associated with task variables like target location or reach direction^50–52^. We constructed the target location subspace separately for the static and moving conditions using respective neural activity from 100 ms to 200 ms after TO. In the moving condition with a target velocity of −120°/s, the neural states corresponding to different reach directions rotated clockwise around the baseline neural state (TO+100 ms) until GO, while neural states in the static condition remained largely unchanged (**Fig. 5c, d**). This suggests the presence of a subgroup of neurons responsive to target motion. In the reach direction plane, constructed with neural activity 100 ms before MO, the organization of baseline neural states (MO-100 ms) aligned precisely with the actual reach directions. Notably, the neural states in the moving condition also rotated clockwise in the original coordinate system, gradually evolved towards the states near MO (**Fig. 5e, Supplementary Fig. 13**). Conversely, neural states in the static condition displayed organization resemble four reach directions around 300 ms after TO, followed by a large bound and mixed distribution, which later aligned with the moving condition around MO. This finding confirms the third type of PD pattern **(Fig. 4e**), indicating that a subgroup of neurons shifts their PDs at late delay period and then reverts upon MO.

Given the sensory and motor representation differences revealed by TDR, we further examined the information conveyed by neural population activity through linear decoding of target location and reach direction (**Fig. 5f**). In the static condition, decoding accuracy of reach direction (equivalent to target location) in y-axis peaked shortly after TO, then dropped to near chance level before rising again after GO, likely reflecting the rapid formation of the motor plan. The moving condition showed a similar rise-and-drop pattern in decoding accuracy for instantaneous target locations.

However, the accuracy for final reach direction increased steadily from TO to MO, indicating a progressive motor plan formation during the delay period. Monkey L’s results differed slightly, as both target location and reach direction decoding accuracy increased after TO. This may be attributed to the high correlation between reach directions and initial target location. Decoding results for x-axis and a combined cosine similarity metric consistently supported the difference between the static and the moving conditions (**Supplementary Fig. 14**). Overall, we hypothesize that there are two distinct preparatory epochs in the static condition: one involving sensorimotor transformation shortly after TO, and another at late delay period when the motor plan is maintained. In contrast, the moving condition experiences continuous sensorimotor transformation or motor planning throughout the entire delay period (**Fig. 5g**).

Cortical neural dynamics can be shaped and strengthened by external inputs. Recent studies have shown that the motor cortex receives essential pattern-generating inputs from the thalamus during movement generation, providing a basis for reliable motor outputs^33^. Modeling suggests that optimal feedback inputs from the thalamus could help resolve preparatory motor errors, thus resisting perturbation^35^. Additionally, preparatory activity in premotor cortex remains resilient even under large-scale unilateral silencing, likely due to corrective information conveyed by the opposite hemisphere^53^. Considering that continuous sensorimotor transformations involve a steady influx of inputs to the motor cortex^54–56^, we aim to develop a model to test whether differences in preparation-related external inputs could lead to differing neural dynamics and ICMS responses.

### Input-driven model simulates ICMS effect on RTs

To test our hypothesis, we utilized an input-driven neural network model developed by Kao et al., that employs optimal feedback control to realize fast motor preparation ^35^. During a reaching task, upstream areas received target information and form a motor intention, the desired optimal states *x*^*^ (fixed points of the population dynamics corresponding to the future reach directions). Based on *x*^*^ and current states of motor cortex, a control input *u* emerges from other recurrently connected brain regions would drive the circuit into the appropriate state, and control the arm to conduct a desired reach (**Fig. 6a**). This network consists of N_*E*_ = 100 excitatory (E) and N_I_ = 100 inhibitory neurons (I), with a fixed optimized connectivity matrix, *W* that maintains E/I balance^57,58^:

**Fig. 6.**
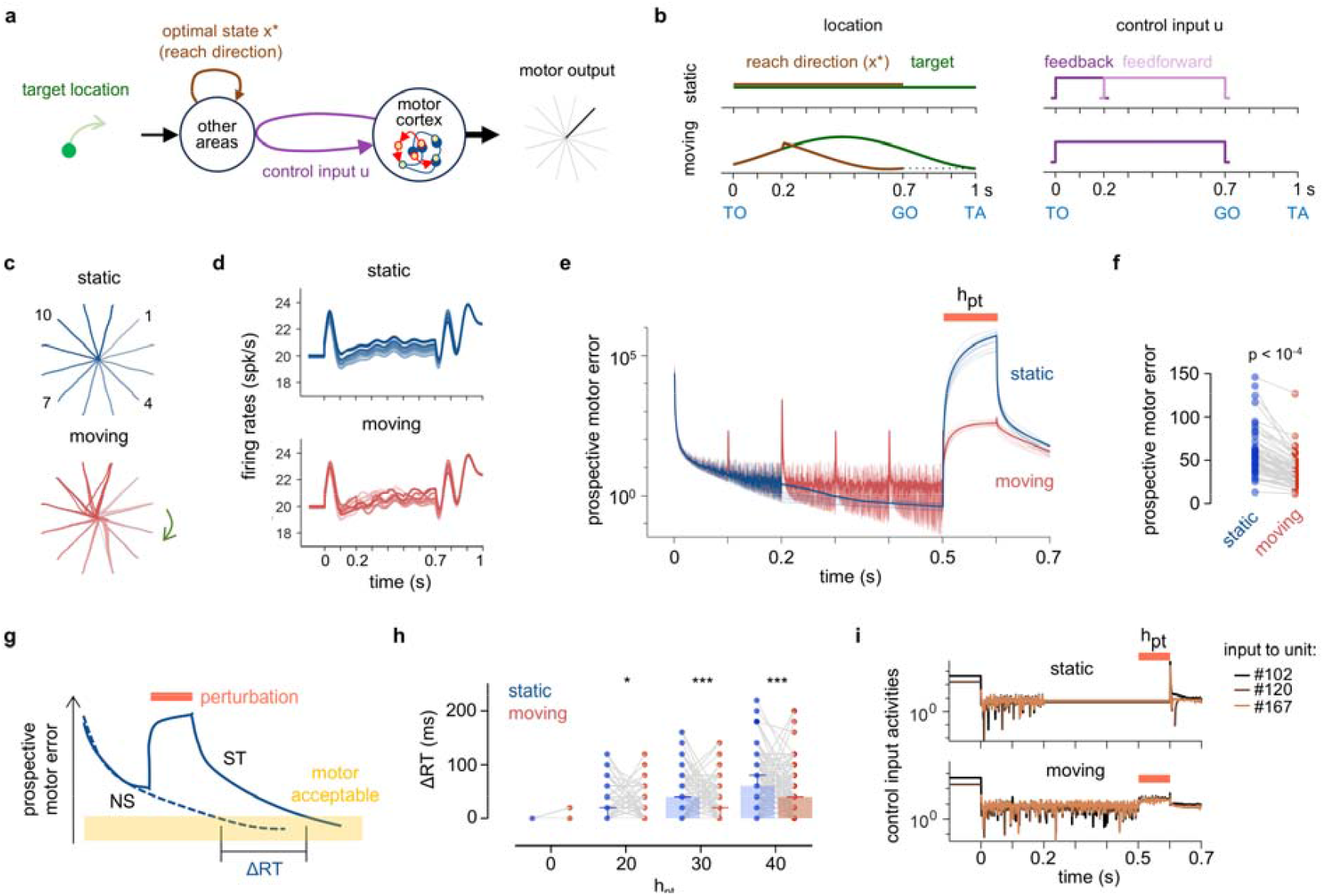
Input-driven network model reproduces ICMS effect on RTs. **a**.Diagram of input-driven network model. During goal-directed movement, target information is translated to motor intentions (optimal state x*) by upstream areas. Control input u from recurrently-connected areas drives motor cortex activity x (with balanced excitatory-inhibitory connectivity) towards an optimal subspace during preparation, with initial states converted to motor outputs (hand positions). **b**.Illustration of optimal states and inputs. In the static condition, the optimal state is fixed, while in the moving condition, it is adjusted to lead the target. Control input u in the static condition is feedback-driven initially, switching to feedforward, while remaining feedback-driven in the moving condition. **c**.Reach trajectories generated by the model. Dashed black lines indicate desired movement paths. Ten trials with varying initial values are shown for each reach direction. **d**.Example PETH for a model unit (#197). Firing rates (mean ± 95%CI) across reach directions are averaged over 10 repetitions with varied initial states (color-coded as in panel **c**). Data before time 0 are extended to clarify initial values. **e**.Prospective motor error in perturbed condition. Perturbation input *h*_*pt*_ (orange bar) of amplitude 50 a.u. is applied 200 ms before MO. Thick lines show mean prospective motor error across 12 directions with 10 initial values. Thin lines show10 randomly selected examples. **f**.Prospective motor error at MO under perturbation (20 repetitions). Perturbation details match those in panel **e**. Statistical significance was assessed using Wilcoxon matched-pairs signed rank test. **g**.Schematic for computing ΔRT. Movement onset occurs when hand trajectory *R*^*2*^ > 0.9, indicating prospective motor errors have declined to permit accurate movement. ΔRT = RT_perturbed_–RT_intact_, with RT_intact_ = 0. **h**.ΔRT in the static and moving conditions. Higher *h*_*pt*_ amplitudes correspond to longer ΔRTs (20 repetitions). Statistical significance tested via Wilcoxon matched-pairs signed rank test (*p < 0.05, *** p < 0.001). **i**.Example of control input u activity during a perturbed trial. Each line represents the u to a specific unit. Data before time 0 are extended for clearer initial values.

**Fig. 7.**
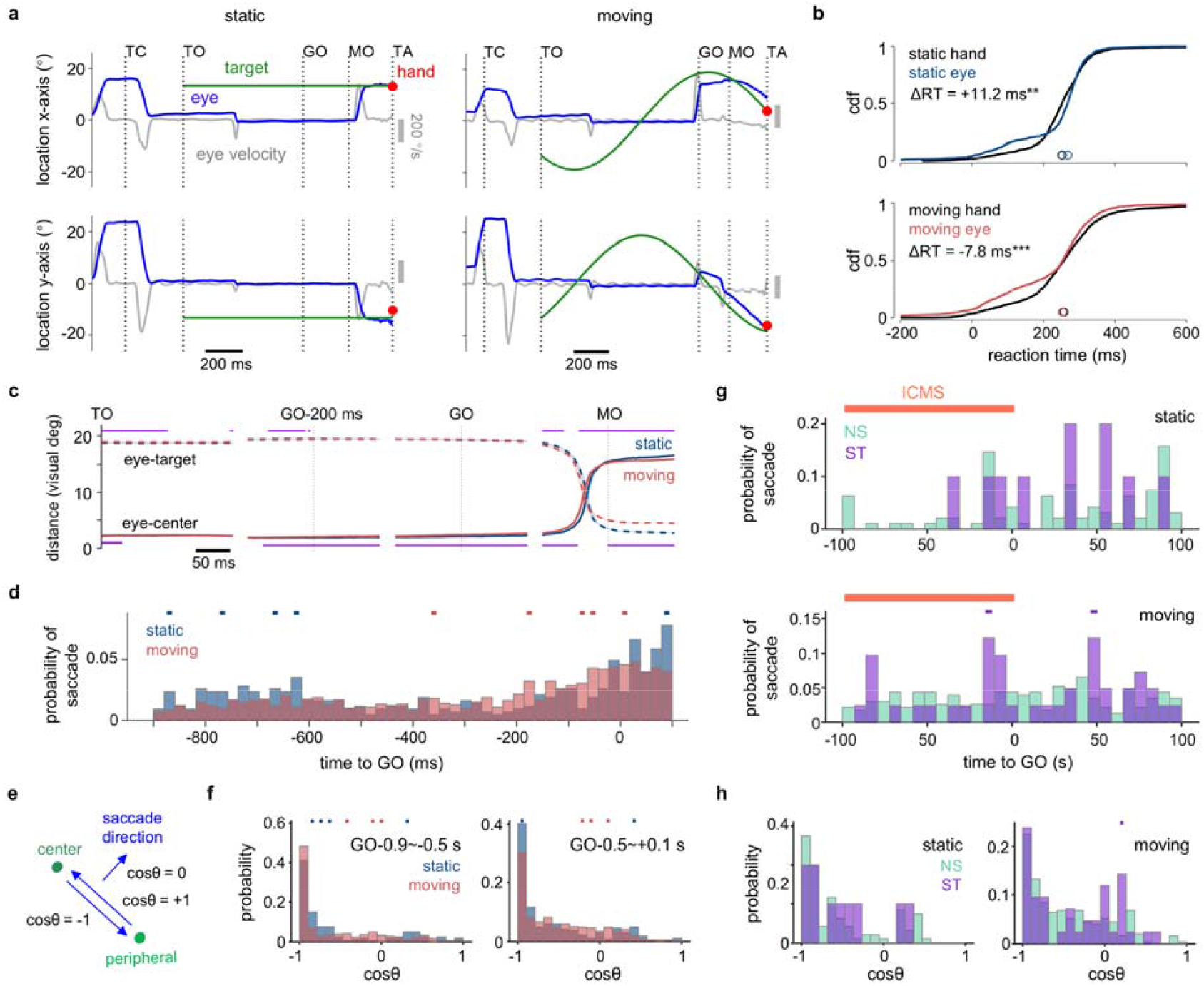
Potential explanation for the ICMS-induced acceleration effect in the moving condition. **a**. Eye and target positions, along with eye speed in a representative trial of the static (left) and moving (right) condition. Gray vertical bar shows the eye speed scale. The red dot indicates hand touch positions. **b**. Empirical cumulative distribution functions (CDFs) of reaction times for eye movements and hand reach movements, with the RT difference (ΔRT = RT_eye_ – RT_hand_) indicated. Trials: static n = 1927; moving n = 2486. **c**. Distance between the eye and center GO (mean ± 95%CI). Purple lines above (eye-target) and below (eye-center) indicate significant differences between conditions (Wilcoxon rank-sum test, p < 0.05). Trial numbers are consistent with panel **b**. **d**. Probability of saccade compared between the static and moving conditions. Colored dots indicate higher probabilities determined by Fisher’s exact test, p < 0.05 (same annotation applies to panel **g, f**, and **h)**. Trials: static n = 755; moving n = 2588. **e**. Schematic illustrating how the cosθ metric is used to determine saccade direction. **f**. Distributions of cosθ for saccade of two representative epochs in panel **d**. **g**. Probability of saccade compared between NS and ST trials in the static (top) and moving condition (bottom), with a target velocity of ±120°/s. The orange line marks ICMS timing (GO-100ms to GO). Trials: static NS n = 391; static ST n = 76; moving NS n = 971; moving ST n = 138. **h**. Distributions of cosθ for saccade in panel **g**. ST trials in the moving condition showed higher proportion of saccade directed away from target.

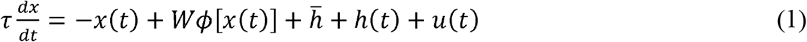

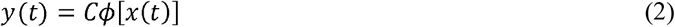

Where 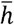 represents the baseline activity of a set of spontaneous firing rates *x*_sp_ ∼*N*(20, 2); *h* (*t*) is a condition-independent spatially uniform α-shaped input bump applied during movement; *u*(*t*)is the external feedback control input, which is only active during preparation period and is deactivated upon movement execution. The function ϕ (*x*) = max (*x*, 0) is a rectified-linear activation function that converts internal activations into momentary hand position *y*(*t*). To complete a delayed reach task of twelve radially located targets, with delay period of 700 ms and movement time of 300 ms, we optimized the desired optimal states *x*^*^ (fixed points of the population dynamics corresponding to the future reach directions) and the readout matrix *C* ∈ℝ^2 ×*N*^ to by minimizing a loss function (**Supplementary Fig. 17a, Method**).

We adapted the model for our task with two key assumptions: (1) *Rotating desired states in the moving condition*: desired optimal states x* update sequentially between adjacent radially-organized optimal states to simulate continuous sensorimotor transformation in the moving condition (**Supplementary Fig. 17b, c**). The desired state’s corresponding reach direction is ahead of the instantaneous target location to ensure accurate interception (**Supplementary Fig. 16**). In the static condition,*x* ^*^ remains constant throughout preparation and execution (**Fig. 6b**). (2) *Switching control strategies in the static condition*: in the original study^35^, “optimal feedback control” strategy computed *u*(*t*) as *u*_opt_=*u* ^*^ +*K* δ *x*(t)using Linear Quadratic Regulator (LQR) algorithms, where δ *x*(t)is the momentary deviation; *u* ^*^ is the movement-specific stationary external input; *K* is the optimal gain matrix. By contrast, “naive feedforward control” strategy, which used a stationary input *u*(*t*)= *u*^*^, was dismissed for its slower stabilization of the network (**Supplementary Fig. 17d**). However, this strategy effectively maintains a desired state, consistent with the stationary neural states observed in the late delay period of the static condition. Thus, we applied the “feedforward control” in the static condition starting 200 ms after TO, transitioning from “feedback control” once the desired state is achieved. In the moving condition, to fulfill both the continuous transformation and accurate interception, “feedback control” remains active throughout preparation. Additionally, we required a delayed perturbation response in the static condition, allowing switch from “feedforward” to “feedback” strategy only after the perturbation ends.

Our model generated hand reach trajectories with high accuracy (**Fig. 6c**), with neuronal units showing stable directional tuning in the static condition and more complex firing patterns in the moving condition (**Fig. 6d, Supplementary Fig. 17g, h**). To simulate the ICMS effect, we applied a perturbation input, *h*_*pt*_, to half of the network units for 100 ms, 200 ms before MO. We assessed perturbation-induced errors using prospective motor error metrics, which estimate potential movement error if initiated from the current neural state^35^. Lower motor errors correspond to more accurate reach trajectories. As expected, perturbations resulted in higher prospective motor errors in the static condition. In contrast, the feedback-controlled network in the moving condition rapidly corrected errors (**Fig. 6e, f, Supplementary Fig. 14i**). By setting an accuracy threshold (*R*^*2*^ > 0.9), we measured the additional times, RTs, needed for errors to fall below this threshold (**Fig. 6g**). Consequently, perturbations led to longer RTs in the static condition, with increased *h*_*pt*_ amplitude correlating with longer RTs (**Fig. 6h**). During perturbation, inputs in the moving condition responded immediately, consistent with feedback control. In the static condition, inputs remained constant due to stationary input, adjusting only after perturbation ceased and feedback control resumed (**Fig. 6i**). Overall, this model supports the hypothesis that sensorimotor transformation-related external input to motor cortex could enhance resilience to perturbations by efficiently reducing errors.

### Potential explanation for the ICMS-induced acceleration effect in the moving condition

An open question in our study is the PreGO ICMS-induced RT-shortening effect in the moving condition. Unlike the RT-prolonging effect in the static condition (**Fig. 2f, g**), this RT reduction does not align with the expected antero-posterior location relationship, suggesting a different underlying mechanism. Since this reduction in RT was achieved without sacrificing reach accuracy, we suspect it may reflect a voluntary decision rather than a passive response. To explore this, we analyzed eye movements during ICMS experiments (**Supplementary Table 3**) and found that in the delayed interception task, the monkeys faced a trade-off between promptly responding to GO and accurately tracking the target. PreGO ICMS applied just before GO may act as an implicit cue that facilitates recognition of visual GO, thereby shortening RTs for interception.

Before the GO, the monkey’s gaze tended to fixate near the center. After GO, monkey would make a saccade followed by smooth pursuit of the target (**Fig. 7a**). While eye movements lag behind the instantaneous location of the moving target^16^, they still provide delayed but valuable information about the target^59–61^. Notably, in the static condition, saccade onset was generally slower, often not occurring until close to MO. In contrast, in the moving condition, saccade frequently began almost immediately after GO. As a result, RTs for eye movements were longer than those for hand movements in the static condition, but shorter in the moving condition (**Fig. 7b**). The distance between eye and center also increased earlier in the moving condition (**Fig. 7c**). The timing of saccade onset also varied between conditions. In the static condition, saccade mainly occurred after TO and after GO, whereas in the moving condition, they often occurred during the late delay period (**Fig. 7d**). This distribution mirrored the epochs showing significant directional tunings in neural activity (**Fig. 4c**), raising the question of whether these saccades were target-related. To determine saccade direction, we calculated the metric ‘cosθ’, the cosine of angle between the eye movement direction vector at the peak and the relative vector pointing to the center (**Fig. 7e**). We found that most cosθ were around −1, both in the early and late delay period, suggesting that these saccades were primarily from center to peripheral (**Fig. 7f**). These findings imply that high direction-tuning in neural activity may be correlated with target-related saccades, and sensorimotor transformations remain highly active just before movement onset in the moving condition, which may potentially interfere with GO recognition. By contrast, gaze remained largely centered in late delay period of the static condition, which can facilitate prompt responses to GO. This may partially explain the shorter RTs observed before ICMS in the static condition and the longer RTs in the moving condition (**Fig. 1g**).

Intriguingly, we found that ICMS applied before GO (GO-100 ms to GO) significantly increased the probability of saccade occurring around GO and GO+50 ms in the moving condition, but not in the static condition (**Fig. 7g**). This increased saccade also appeared with other target velocities, showing a similar peak around GO+50 ms (**Supplementary Figure 18a**). In the moving condition, cosθ has a higher portion above zero in ST trials, suggesting ICMS induced more saccade directed away from the peripheral. By contrast, in the static condition, there was no such difference between NS and ST trials (**Fig. 7h**). Further examination of other ICMS timings revealed no increase in saccade probability (**Supplementary Figure 18b, c**). Previous research shows that ICMS can evoke sensations in non-sensory cortical regions^62–64^ and even instruct specific actions when applied to premotor cortex^65^. Integrating multiple sensory cues enhances perceptual sensitivity^66^, and combining ICMS with weak visual signals improved target estimation^67^. Given that ICMS influenced the timing of saccades around the GO, we propose that ICMS might act as an auxiliary cue to facilitate movement initiation, when preparatory neural activity is undisturbed and prompt recognition of GO is needed.

## Discussion

Despite that cortical control of movement has undergone conceptual shifts from representational to dynamical systems view over the decades^3,5,68^, how neural dynamics adapting to dynamic environment or perturbations is not yet thoroughly understood. In this study, we explored the intriguing effect of ICMS on a novel reaching task, proposing that continuous sensorimotor transformation may enhance the robustness of motor cortex against ICMS perturbations. Monkeys exhibited similar reach trajectories towards both the static (center-out task) and the moving targets (interception task) (**Fig. 1**). However, ICMS applied during the late delay period selectively prolonged reaction times for reaching the static targets, but not the moving targets (**Fig. 2**). Post-ICMS neural states deviated more significantly from intact states in the static condition, and the temporal shift between non-stimulated and stimulated states predicted changes in RTs (**Fig. 3**). Target or reach direction representations for the moving targets adapted continuously with target motion and remained stable throughout the delay period. In contrast, representations for the static targets were less stable in the late delay period, exhibiting two distinct preparation epochs (**Fig. 4, 5**). To simulate these effects, we adapted an input-driven neural network model and successfully replicated the differential impact of ICMS on RTs, demonstrating that optimal feedback inputs from other brain regions can efficiently mitigate perturbations-induced errors by in the moving condition (**Fig. 6**). Ultimately, we propose that, in the absence of disturbances, PreGO ICMS may act as a go signal that accelerates movement onset during delayed-instructed interception (**Fig. 7**).

The RT-prolonging ICMS effect observed in center-out task supported the hypothesis that the brain can detect errors and delay movement until they can be corrected^11,69^. Our study offers direct evidence for this hypothesis by showing that ICMS in the static condition caused neural states to deviate, requiring time to recover. Surprisingly, however, ICMS did not disrupt reach accuracy or increase RTs in interception task. To address potential biases from randomness and trial repetitions imbalances, we conducted control experiments confirming the robustness of this phenomenon across various task designs (**Supplementary Figure 3-5**). Interestingly, a similar ICMS-resilient effect was noted in a previous study: ICMS to supplementary motor area (SMA) during interceptive reaches to outward-moving targets resulted in a smaller RT increase^15^. However, this study did not fully explain this ICMS effect. Our findings suggest a plausible mechanism aligned with the optimal-subspace hypothesis, indicating that disrupting activity when it is still variable has minimal impact^11^. Our results showed that the moving target induced continuous sensorimotor transformation in motor cortex, manifesting as varied neural dynamics throughout the delay period. However, contrary to the mechanism suggested by the hypothesis that perturbation merely shifts the system between non-optimal states^11^, we observed less deviation and faster recovery of neural states in the moving condition, suggesting quicker error resolution. Thus, our study provides an alternative explanation for ICMS resilience: continuously evolving preparatory neural states operate as part of an optimal feedback controller, akin to a bicycle maintaining stability while in motion, naturally resisting disturbance.

Our findings reveal notable differences in preparatory neural dynamics between the static and moving conditions, despite similar reach kinematics. Similar to prior findings^13,14,70^, we observed partial but not complete overlap in preparatory space across contexts (**Supplementary Figure 15**), reflecting that commonality and divergence coexist: shared traits reveal connections, while unique features highlight distinctions. ICMS effect could be similar across contexts, suggesting that perturbation may affect these shared neural dynamics^15^, while our results show that distinct perturbation responses may emerge from differences in dynamics. Recent study demonstrated that behaviors change only when perturbation impact neural states within the task dynamics space, highlighting the importance of identifying the intrinsic manifolds for causal inferences^42,71^. Recent progress unveils a more integrated sensory-motor network than previously recognized^72–74^, reinforcing the view that cortical dynamics are shaped by external inputs, and considering these factors is crucial for accurate interpretation of intrinsic neural behaviors^75^. Pattern-generating inputs from thalamus to motor cortex are essential for movement generation^33,35^. Optimal input-driven neural dynamics generate multiple preparation phases that rely on inputs preceding movement in reach sequences^76^. Our recent study found that motor cortical neurons are tuned to both reach direction and target motion, suggesting sensory signals contribute to shaping neural dynamics during interception^23^. We also showed in a model that different input control strategies can lead to distinct preparatory dynamics and responses to perturbation. Therefore, further investigation using techniques that directly measuring external input could illuminate some prior paradoxes and deepen our understanding of neural dynamics^12,14,32,70^.

Recent progress in modeling have reshapes our comprehension of motor preparation and the role of external inputs in neural dynamics^35,36^. By simulating the motor cortex as a self-contained, low-dimensional dynamical system, recent study effectively explained the differing responses to optogenetics and ICMS^42^. In this study, we utilized a well-established input-driven neural network model to validate our hypothesis that the differential ICMS effect across conditions stem from differences in neural dynamics driven by external inputs. While our model’s performance aligns with the findings of Kao et al^35^, which was our expectation, a novel finding from our model is that the ‘optimal feedback control’ strategy is a pre-requisite for accurate interception, and for replicating key features of its preparatory activity. In addition, since the “naïve feedforward strategy” was not intended as the true mechanism for motor preparation—serving only as a control for the “feedback strategy” in Kao et al.’s work-it is significant that we identify a connection between this strategy and the stationary late-delay neural dynamics in the static condition. Thus, our model’s findings refine and expand the framework proposed by Kao et al. to better explain various phenomena related to motor preparation. Nevertheless, our model has several limitations. First, we simplified the model by leaving aside circuit architecture, and made simple assumption about perturbation *h*_*pt*_, which may fail to capture the nonlinearity of real neural networks and lead to discrepancies between our model’s predictions and empirical data, such as the absence of a significant correlation between *h*_*pt*_ amplitude and RTs. This issue could be resolved by examining the results at a nonlinear circuit level. Second, our model relies on multiple assumptions based on our empirical observations, in contrary to task-based models like recurrent neural networks (RNNs) ^8,77–79^. Given that preparatory activities can arise from model with physiological constraints rather than being assumed from the outset^36^, and RNNs with spatio-temporal structures can better simulate shift between computations^78,79^, future research could benefit from these designs to reduce artificial assumptions. Nevertheless, our simulation offers a plausible explanation for the distinct ICMS effects, and lays the groundwork for future validation through physiological evidence and biologically interpretable models.

One limitation of our study is the relatively small size of the recorded neuronal population, which may affect the interpretation of subtle differences in neural dynamics, such as tortuosity difference in neural trajectories. Differences in tortuosity could arise in certain reach directions from biased population PD or neurons exhibiting condition bias. However, the detour index quantifying tortuosity is averaged across reach directions and sessions, and the proportion of neurons showing condition preference is negligible, making it unlikely that tortuosity differences were influenced by number of neurons. Additionally, we conducted bootstrapping analyses on the neuronal data, confirming that our findings are not overly sensitive to the specific number of neurons recorded (**Supplementary Figure 12**). Given previous discussions about the nature of the ‘hold’ states in the center-out task^12^, we propose that the twisting neural trajectories may reflect a computation shift from sensorimotor transformation to other preparatory activity, such as time estimation^46,47^. Both the unstable PD and prolonged eye fixation on the center suggest that responding promptly to go cue is a primary demand for the static condition at late delay period^80–84^. Therefore, it is conceivable that, with limited preparatory time, these ‘hold’ states might be bypassed^12^. Future work should aim to find direct evidence for the “time estimation” process in delayed-instruction reach tasks to enhance our understanding of motor preparation^85,86^.

A remaining question is why PreGO ICMS reduced RTs in the moving condition. Although this phenomenon is not the primary focus of our work, we offer a plausible explanation: ICMS disrupts motor preparation equally in the static and moving conditions, but it also provides information about go cue timing. Given the robustness against perturbations conferred by varying neural dynamics and the stricter timing requirements in the moving condition, monkeys may utilize the informative ICMS “cue” to reduce their RTs compared to the static condition. We observed that including a delay period resulted in longer RTs for interception, contrasting with previous studies that did not incorporate a delay^13,15,16,40,41^. The precise time-to-contact required for interception^87–89^, along with the constrained reach time, may create a conflict between estimating target motion and go cue, potentially delaying movement onset (**Fig. 7**). As our understanding of the action maps across the interconnected fronto-parietal circuit continues to grow^90,91^, the remote effects of ICMS are receiving increasing attention^92–94^. Our data suggest that ICMS may serve as an implicit cue to facilitate go cue recognition. However, alternative explanations remain viable. For example, “StartReact” effect suggests that a prepared movement could be accelerated by an unexpected stimulus, typically acoustic^95^. Additionally, GABAergic inhibition in PMd can induce premature reaching^96^, although this is rare. No single mechanism fully accounts for the differential ICMS effects observed, underscoring the need for future research using more specific perturbation tools such as cooling^97^ and optogenetics^98^ to further unravel these effects.

In conclusion, our study demonstrates that continuous preparatory neural dynamics, shaped by recurrent external input, enhance the resilience of motor control networks during manual interception. This emphasizes that motor cortex dynamics arise from a sensorimotor transformation, integrating peripheral sensory inputs with contextual information from other cortical and subcortical areas. This interplay highlights how continuous growth, both in life and neural dynamics, enables individuals and systems to better adapt to and withstand external interference.

## Methods

### Experimental model

This study involved two adult male rhesus monkeys (*Macaca mulatta*): monkey G (8.4 kg, 13 years old) and monkey L (10.0 kg, 6 years old). Each monkey was implanted with head fixation posts and an acrylic recording chamber (19 mm inner diameter), which was secured with dental acrylic resin and titanium screws. The chambers were positioned over the upper limb areas of the primary motor cortex (M1) and the dorsal premotor cortex (PMd) in the right hemisphere for monkey G and the left hemisphere for monkey L. All procedures were in accordance with NIH guidelines and were approved by the Institutional Animal Care and Use Committee (IACUC) of the Institute of Neuroscience, CAS.

### Task design and apparatus

In the manual delayed interception task ^16^, a head-fixed monkey was seated in a custom primate chair facing a vertical touchscreen. The arm contralateral to the chamber was free to move, while the ipsilateral arm was restrained by a shutter board (**Supplementary Fig. 1a**). The task began with the monkey touching the central target for 600 ms. Subsequently, a peripheral target appeared (target on, TO), which either remained static (i.e., center-out task) or moved in a fixed-radius circular path at specified velocities. Following a delay period, the central target disappeared (Go cue, GO), and monkey was required to reach the peripheral target within 800 ms. If the touchpoint is within the error tolerance and the target is held for more than 600 ms, the touchpoint turns red, indicating a successful trial, and the monkey received one drop of liquid food as a reward; otherwise, the touch point turned blue (failed trial).

Target velocities were customized for each monkey, with variations including static (0°/s) and moving targets at clockwise (−) or counterclockwise (+) velocities (e.g., ±120°/s, ±180°/s, ±240°/s). The delay period ranged from 0 to 1000 ms, and initial target locations varied from 0 to 360°. These parameters were randomly assigned for each trial to ensure variability.

Task control and data acquisition were managed via MonkeyLogic ^99^ [https://monkeylogic.nimh.nih.gov/index.html], which logged touch interactions and target locations. Reach movements were tracked using a Vicon motion capture system (Oxford, UK) equipped with MX3+ cameras that captured the trajectory of an infrared reflective marker attached to the monkey’s free-moving hand at 100 Hz. Additionally, eye movements were recorded with an Eyelink infrared camera at 1000 Hz, and electromyograms (EMGs) were collected in some sessions using Delsys wireless sensors on the triceps, forearm lateral muscle, or forearm interior muscle at a sampling rate of 1259 Hz.

### ICMS task design

For intracortical microstimulation (ICMS) recording experiments, each session contains static and one moving target velocity condition. Two delay periods were selected: 200 ms (short) and 900 ms (long). To ensure a uniform distribution of touch points across the four screen quadrants, the initial target locations were carefully selected to align with the individual reaction times and delay periods. ICMS is delivered at two timings: within 100 ms (0-100 ms) before GO (PreGO) and within 150 ms (0-150 ms) after GO (PreMO). There are three types of trials, designed static condition, designed moving condition, and random condition. Designed-condition trials have four initial target locations, with an equal number of condition-matched (the same target velocity, initial target location, and delay period) stimulated (ST) and non-stimulated (NS) trials. Trials in the random condition are NS trials with moving targets and random initial target location. Designed ST trials were interleaved with designed NS trials and trials from the random condition. We limited the number of ST trials to be fewer than 200 per session for each recording site to protect the brain tissue. After all the designed NS and ST trials were completed, there would be 100-200 random condition trials before monkey ceasing the task naturally due to satiation. We deleted the non-stimulated trials after the last designed trial to prevent the slow drift in RTs. For monkey G, we recorded 7 sessions with target velocities of 0°/s and −120°/s, and 7 sessions at 0°/s and −240°/s. For monkey L, we recorded 12 sessions at 0°/s and −120°/s, and 3 sessions at 0°/s and 180°/s (**Supplementary Table 1, 2**).

**Table 2.**
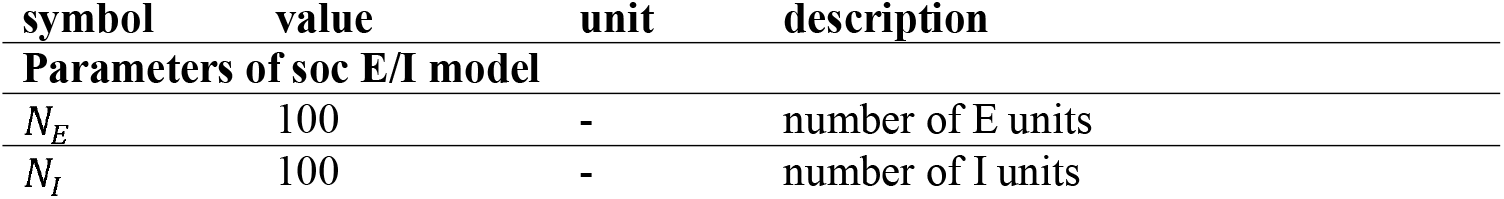
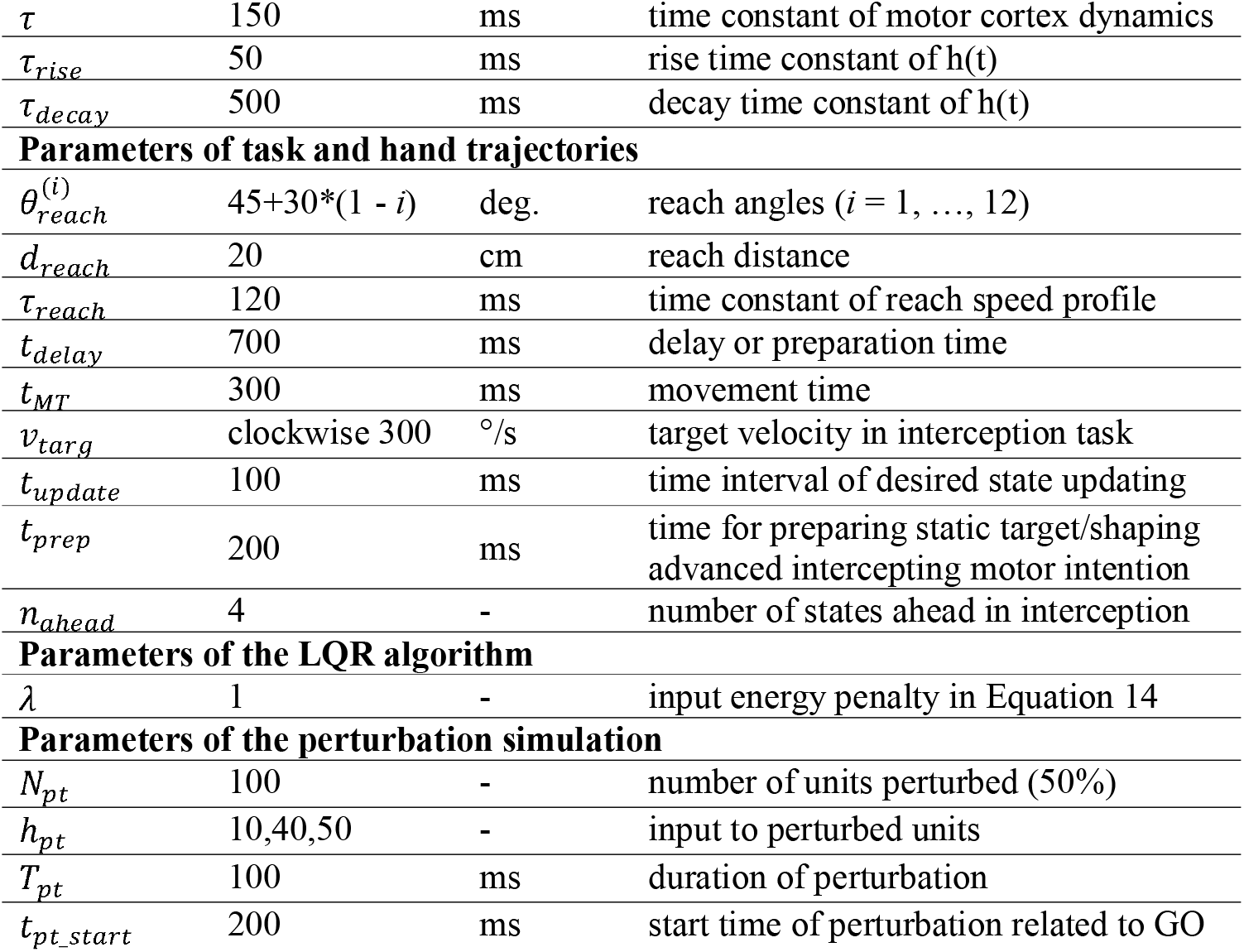
values of all the parameters of soc model.

| symbol | value | unit | description |
| --- | --- | --- | --- |
| **Parameters of soc E/I model** |  |  |  |
| $N_E$ | 100 | - | number of E units |
| $N_I$ | 100 | - | number of I units |
| $\tau$ | 150 | ms | time constant of motor cortex dynamics |
| $\tau_{rise}$ | 50 | ms | rise time constant of $h(t)$ |
| $\tau_{decay}$ | 500 | ms | decay time constant of $h(t)$ |
| **Parameters of task and hand trajectories** |  |  |  |
| $\theta_{reach}^{(i)}$ | $45+30*(1-i)$ | deg. | reach angles ($i = 1, \dots, 12$) |
| $d_{reach}$ | 20 | cm | reach distance |
| $\tau_{reach}$ | 120 | ms | time constant of reach speed profile |
| $t_{delay}$ | 700 | ms | delay or preparation time |
| $t_{MT}$ | 300 | ms | movement time |
| $v_{targ}$ | clockwise 300 | °/s | target velocity in interception task |
| $t_{update}$ | 100 | ms | time interval of desired state updating |
| $t_{prep}$ | 200 | ms | time for preparing static target/shaping advanced intercepting motor intention |
| $n_{ahead}$ | 4 | - | number of states ahead in interception |
| **Parameters of the LQR algorithm** |  |  |  |
| $\lambda$ | 1 | - | input energy penalty in Equation 14 |
| **Parameters of the perturbation simulation** |  |  |  |
| $N_{pt}$ | 100 | - | number of units perturbed (50%) |
| $h_{pt}$ | 10,40,50 | - | input to perturbed units |
| $T_{pt}$ | 100 | ms | duration of perturbation |
| $t_{pt\_start}$ | 200 | ms | start time of perturbation related to GO |

### Electrophysiological recording and ICMS parameters

In each ICMS session, we delivered stimulation using single tungsten microelectrodes (#UEWLEFSECN1E, 0.6-1.2 MΩ @1 KHz, FHC) and recorded neuronal activity with 64-channel S-probe with interelectrode spacing of 75 μm (#PLX-SP-64-15ST(100/75)-(320-140)(640-10)-570-(2)CON/32m-V, 0.3-0.5 MΩ @1 KHz, Plexon).

These electrodes were advanced into the cortex through a stainless-steel guide tube using a microdrive (Thomas Motorized Electrode Manipulator, Thomas RECORDING), which was mounted on the recording chamber. Typically, the microelectrode and S-probe were spaced approximately 4-5 mm apart, positioned either at M1 or PMd.

Neural signals were recorded either from the microelectrode before ICMS using an AlphaLab SnR Stimulation and Recording System (Alpha Omega), sampled at 44 kHz to confirm the presence of arm movement-related neural activity, or from the S-probe during ICMS using Cerebus acquisition system (Blackrock Microsystems), sampled at 30 kHz. To determine the depth of the recording site, the depth where the first neural activity was observed was set as the 0 position, with subsequent depths relative to this. Single-unit spike sorting was performed semi-automatically using Spike2 (Spike2 7.15, CED) and Wave_clus^100^.

ICMS were delivered using an AlphaLab SnR Stimulation and Recording System (Alpha Omega). Each ICMS consisted of a 100-ms train of biphasic pulses (300 Hz, 200 μs per phase, with cathode leading). To determine the amplitude threshold of each site, the monkey was instructed to keep his hand on the center of the screen to receive a reward. Stimulation amplitude was increased from 30 μA in 10 μA increments until a muscle twitch or forelimb displacement was detected, either visually or palpably, or until reaching to a maximum of 120 μA. Subsequently, the amplitude was reduced in 5 μA decrements until no muscle twitch or movement was detected. In this study, ICMS amplitudes remained below (5-10 μA lower) the threshold (M1: 10-120 μA, median 55 μA; PMd: 20-120 μA, median 62 μA).

### Behavioral analysis

Hand velocities were derived from recorded 3D reach trajectory sampled at 100 Hz. Movement onset (MO) time was determined by identifying the time of peak hand speed and tracing back until the speed dropped below 5% of the peak. Using this MO time as a reference, we computed the reaction times (RTs, from Go to MO), movement time (MT, from movement onset to screen touch), and RT+MT.

Endpoint errors were the distances between the reach endpoint and the target. Reach endpoints were monitored by a touchscreen via MonkeyLogic, and 95% covariance ellipses were computed relative to the final target location.

Reach trajectories were projected into 2D within the vertical space. Confidence intervals for the reach trajectory were calculated using Teetool^101^, which models the 2D trajectories as a Gaussian process, producing an area that encompasses the 1σ covariance around the mean path. To facilitate trajectory comparison, reach trajectories were rotated to align the head to the origin, and the tail to 45°, 135°, 225°, and 315°, respectively.

The curvature of the trajectories was quantified using the launching angle relative to the endpoint, defined as the angle between the launching direction and the final reach direction upon touching the target. The launching direction was determined as the reach direction 200 ms after movement onset to mitigate potential biases from trajectory jitter at MO.

To account for trial number imbalances and to prevent bias in examining the ICMS effect, we computed the difference (Δ) between stimulated (ST) trials and condition-matched non-stimulated (NS) trials. Specifically, each ST trial was paired with condition-equivalent NS trials within the same session, sharing identical delay duration, target velocities, and initial target position (±10°). The median of kinematic parameters (e.g., RTs) from the matched NS trials was subtracted from each ST trial and the corresponding NS trial within the paired group. Statistical significance was evaluated using the Wilcoxon rank-sum test on both the raw data and difference values (see **Fig. 2**, and **Supplementary Fig. 7, 8**).

### Neural data processing and PETH

For neural data involving ICMS, we restricted our analyses to times outside the ICMS period due to the large ICMS artifacts that confound neural sorting results. For each trial, we identified the start and end of ICMS, and set the neural data within this range to either infinite or zero. When averaging neural data without aligning to the end of ICMS, we identified the earliest start and latest end times of ICMS across trials. This period was defined as the ICMS window for the averaged data. Consequently, data within this window were excluded from the analysis (see **Fig. 3**).

To assess post-ICMS responses, we computed the difference between stimulated (ST) and equivalent non-stimulated (NS) trials. We randomly sampled from the ICMS onset time distribution of a specific condition to simulate the ICMS onset for the condition-equivalent NS trials. Subsequently, we aligned both ST and NS trials to the ICMS onset time and subtracted the average firing rates of equivalent NS trials from those of ST trials. Units were sorted based on mean firing rates within a 100-ms window following the cessation of ICMS train (see **Fig. 3a**).

Normalized firing rates heatmaps were used to assess overall neural activity. Neural activity in 20-ms bin without overlap or smoothing in long delay condition was aligned to GO (GO-1200ms to GO+700 ms), averaged across all other conditions, and normalized to a mean of 0 and a standard deviation of 1. Units were sorted based on the timing of maximum firing rates (see **Fig. 4a**).

Peri-event time histograms (PETHs) and corresponding spike rasters were used to estimate mean firing rates (FRs) over time. Spike trains were aligned to Go, binned in 10-ms bins, and smoothed with a 50-ms standard deviation Gaussian filter. Trials were averaged according to conditions, such as target velocities and reach directions. Differences in FRs between conditions (e.g., static vs. moving) were evaluated using the Wilcoxon rank-sum test at each timepoint (see **Fig. 4b**).

### Histogram of time bins of firing rates discriminating between reach directions

To estimate when the firing rates show discrimination of reach directions, we conducted Kruskal-Wallis test (p < 0.05) for each 50-ms time bin of condition-averaged PETH (GO-1000 ms to GO+600 ms) for each unit. Then, the number of significant time bins was summed across units with a bin width 55 ms. The difference between the static and moving conditions was assessed using Fisher’s exact test (p < 0.05). Only units with mean firing rates greater than 5 spk/sec around GO were selected for computation (see **Fig. 4c**).

### Bootstrap Principal Component Analysis (PCA) with Confidence Intervals

Principal component analysis (PCA) is a dimensionality reduction method that identifies the primary directions of variance in a multidimensional dataset. To estimate confidence intervals for PCA components, we applied a bootstrap resampling procedure to the dataset. We first computed a reference PCA on the mean time-series data for the static NS condition, extracting the principal components (PCs) that serve as the reference structure. Specifically, neural data were initially binned at 2-ms without overlap and smoothed using a 25-ms Gaussian kernel. Subsequently, trial-averaged firing rates were organized into a neural matrix *X* of size *n* ×*ct*, where *n* is the number of recorded neurons, *c* is the number of conditions, and *t* is the number of time points. PCA was then performed on the neural matrix of static NS condition, *X*^*ref*^. The new representations of *X*^*ref*^ in the low-dimensional subspace were computed as 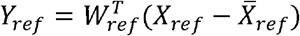, where *Y*^*ref*^ of size *k*×*ct* (with *k<n*) was the PC scores of *X*^*ref*^, and *W*^*ref*^ of size *k*×*n* was the reference projection matrix or loadings that transform the feature space into PC space. Next, we performed *n* = 1000 bootstrap resamples on the dataset. Each bootstrap sample is generated by randomly sampling (with replacement) the trials within each condition. For each bootstrap resample, PCA is recalculated on the resampled data from the static NS condition. This process yields a new set of PCs and scores. To ensure alignment with the original reference PCA components, we use Procrustes alignment to align the bootstrap-derived first three PCs, *Y*_*ctrl*_ with the reference first three PCs, *Y*_*ref*._ This step also transforms the projection matrix, *W*_*ctrl*_ into *W*_*aligned*_. Then, the other conditions *X*_*new*_ (e.g., static ST, moving NS, and moving ST) are projected into the PCA space defined by the aligned components. 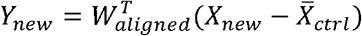.The scores *Y*_*new*_ represent *X*_*new*_ in the principal subspace. After completing all bootstrap iterations, the mean PCA scores are computed across all resamples. Confidence intervals for each PCA score are calculated using percentiles from the bootstrap distribution, specifically the 95% confidence interval (CI). This is accomplished by computing the 2.5th and 97.5th percentiles for each time point and dimension of the PCA space (see **Fig. 3d, e** right).

### Neural trajectories

To visualize the neural trajectories, the trial-averaged conditions were plotted in the subspace constructed using the first 3 PC scores of PCA on *X*_*ctrl*_. For neural trajectory analysis in ICMS data, the firing rates during ICMS were set to zero for each ST trial due to the loss of spikes caused by ICMS artifacts. Next, the firing rates were aligned to GO (−1000ms +500ms), and averaged across reach directions. The PETH in the static NS condition was used to construct the 3D state space, into which the static ST and moving conditions were projected. A representative condition (long delay, reach to quadrant 1, PreGO ICMS from dataset G0914) was selected for demonstration, with the mean MO time labeled on the trajectories. Additionally, the first three PCs are also depicted separately (see **Fig. 3d, e**).

For neural trajectory analysis for paired static and moving condition, the firing rates were aligned to GO and averaged according to the respective conditions. An example reach direction condition (quadrant 1) was selected, with the epoch chosen for the long delay condition from GO-1000 ms (i.e., TO-100 ms) to GO+500 ms (see **Fig. 5a** and **Supplementary Fig. 9**).

### Cross-Correlation Function (CCF) of PCs

To examine the post-ICMS reaction time changes embodied in neural population activity, we computed the cross-correlation function (CCF) between the population components of neural activity in non-stimulating (NS) and stimulating (ST) condition to determine the time lag that maximizes their correlation. First, population responses of static NS, static ST, moving NS, and moving ST conditions were acquired by projecting trial-averaged (of each reach direction) neural activity (2 ms bins with 25-ms Gaussian kernel smoothing, GO-1000 ms to GO+600 ms) into the state space built from the static NS neural activity using PCA. Next, as the largest response component reflects movement timing^46^, the last 200 ms of the first principal components (PCs) of ST and matched-NS trials were selected to calculate the normalized cross-correlation coefficient using MATLAB’s xcorr function. *xcorr(y, x, ‘coeff’)* computes the normalized cross-correlation coefficient *ccf*(τ) for each time lag τ, as follows:

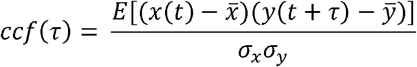

where E is the expectation, 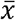 and 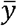 are the means of the PCs of NS and ST conditions, respectively, σ_*x*_ and σ_*y*_ and are their standard deviations. The time lag τ corresponding to the maximum cross-correlation was determined. The index of the maximum value of *ccf*(τ), denoted as *I*, was used to calculate the optimal lag time *T*_*lag*_, which quantifies the temporal shift at which the signals are maximally correlated as follows:

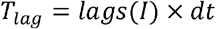

where dt is the time step 2 ms. In our case, positive *T*_*lag*_ indicates a lag of neural states of ST condition compared to NS condition, while negative values indicate a lead of ST condition. By identifying the optimal lag time for each reach condition in each session, we were able to quantify the temporal relationship between the neural states of NS and ST conditions, allowing for the estimation of RT changes post-ICMS (see **Fig. 3f, g**).

### Unbiased neural state distance

To measure the distance or difference between two multivariate distributions, such as mean firing rates in an unbiased way, we adopted the cross-validated estimator of neural distance, developed by Willett et al. (2020) [https://github.com/fwillett/cvVectorStats]. This technique estimates the difference in sample means 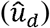 of the multivariate vector of trial i for distribution 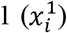, and distribution 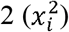 with minimal biases in a cross-validation approach:

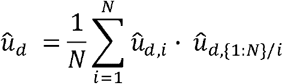

where 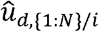 is the sample estimate of the difference in means using all trials except trial i. Neural state distance is the distance between the full-dimensional PCA neural states of stimulating (ST) and matched non-stimulating (NS) condition. Here, we first built the neural state space from trial-averaged static NS condition, and projected the neural activity of each trial into the state space to obtain single-trial neural states (2-ms bins with 25-ms Gaussian kernel smooth), thus generating, for both NS and ST conditions, for each reach direction and each session, an *t* ×*m*× × neural state matrix, where *t* is the number of time bin (MO-1000 ms +600 ms), *m* is the number of trials, and *n* is the number of neurons in each session. For each timepoint, unbiased Euclidean distances were computed for two distributions: *x*_*NS=*_ *m*_*NS*_ × *n* and *x*_*ST*_= *m*_*NS*_ × *n*. Then, single-trial neural distances were aligned to end of ICMS and averaged across reach directions and sessions. Data from ICMS end-600 ms to +300 ms were shown. For monkey G, 7 sessions of −120°/s target velocity with PreGO ICMS and 6 sessions of −240°/s target velocity with PreMO ICMS (totaling 13 4 = 52 samples) were averaged, while for monkey L, 12 sessions of −120°/s target velocity with PreMO ICMS and 3 sessions of +180°/s target velocity with PreGO ICMS (totaling 15 4 = 60 samples) were averaged (see **Fig. 3h**).

### Neural state speed

We computed the neural state speed using the neural states derived from PCA. Specifically, similar to the processes described in **Unbiased neural state distance**, single-trial neural states of each condition are derived. Neural states aligned to MO were organized into a neural state matrix of size *t* ×*m*× *k* for each condition, where *t* is the number of time bin (MO-1000 ms +600 ms), *m* is the number of trials, *k* is the number of PCs that captures >90% variance (k < n, which is the number of neurons per session). Next, the rate of change and Euclidean norm of the neural states were computed to derive overall neural speed matrix of size (t−1) × *m*. Then, data were aligned to the end of ICMS and averaged across reach directions and sessions, which resulted in a comparison between the static and moving conditions in NS or ST condition (see **Fig. 3i**).

### Preferred direction (PD) of neurons

Preferred direction (PD) is computed by summing the product of normalized mean firing rates and movement angles. Specifically, firing rates during each time period (TO-100 ms to TO+700 ms, GO-200 ms to GO+100 ms, and MO-100 ms to MO+200 ms) were binned in 100 ms size with a 50 ms overlap, without smoothing. Mean firing rates for each of the four movement directions were normalized to the 95^th^ percentile of the firing rates across all trials. PD is the angle of the resultant vector obtained by multiplying the mean firing rates with the mean reach direction angles. PD changes are calculated as the difference between adjacent PDs within a condition to estimate the variation of PD (see **Fig. 4d**).

### Detour index

To quantify the aggregation extent of neural trajectories, we defined the detour index, which is the ratio of the actual distance traveled by the neural trajectory to the shortest possible distance. A higher detour index indicates a longer and more redundant traveling path of neural states. Specifically, similar to the bootstrap PCA, we applied a bootstrap resampling procedure to the single-trial firing rates (2-ms bins with 25-ms Gaussian kernel smoothing, GO-1000 ms to GO+300 ms for monkey G and GO-1000 ms to GO+350 ms for monkey L) to obtain the condition-averaged neural states that were projected onto the state subspace built from the static condition. Next, neural states with K dimensions that capture over 90% variance were used to compute the ratio of shortest Euclidean distance between the start (GO-500 ms) and end (GO+100 ms) time points to the sum of the pairwise Euclidean distances between consecutive time points along the neural trajectory. After completing the bootstrap resampling, the mean detour index across bootstraps is obtained for each reach direction of each session. Finally, Finally, a total of 60 paired samples (4 reach directions × 15 sessions) for monkey G, and 68 paired samples (4 × 17) for monkey L were used for statistical comparison (see **Fig. 5b**).

### Targeted dimensionality reduction (TDR)

In the manuscript, we used a simplified version of targeted dimensionality reduction (TDR) firstly introduced by Mante et al. (2013)^50^ and later simplified by Sun et al. (2021)^51^ to identify low dimensional subspaces capturing variance related to the behavioral variables of interest. Specifically, to construct the TDR space, multivariable linear regression is used to determine how various behavioral variables affected the responses of each unit. The neural data, with 100-ms bin size and no overlap or smoothing, were first averaged across all trials for each condition. The activity of each neuron was then centered by subtracting its mean response from the firing rate at each time point in each condition. The centered responses of neuron i as a linear combination of several behavioral variables:

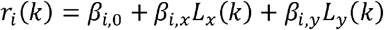

where *r*_*i*_(*k*) t is the centered trial-averaged response of unit *i* on condition *k*, averaged over a certain time window. *L*_*x*_(*k*) and *L*_*y*_(*k*) are the trial-averaged horizontal and vertical instantaneous target location or hand reach location on condition k. The regression coefficient *β*_*i*,0_ captures variance independent of the listed behavioral and task variables. To estimate the regression coefficients, we constructed the behavior variable matrix *M* with shape conditions × regressors (M =[x|1], X ∈ ℝ ^*c*× 2^), and neural activity matrix *R* with conditions × neurons. The regression coefficients could be estimated as:

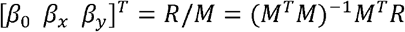

The TDR axes are computed by finding the pseudoinverse of the regression coefficients: *β*=*pinv*(*β*). To ensure that the TDR axes are independent, they are orthogonalized using the Gram-Schmidt orthogonalization process. We then projected neural data into the regression subspace by multiplying the TDR axes with the neural data matrix *R*.

The target location subspace for the static or moving condition was built by regressing neural activity at TO+0.1 s against respective instantaneous target location. The hand reach location subspace for the static or moving condition was built by regressing neural activity at MO-0.1 s against respective final hand touch location. This time (MO-0.1 s) is selected for it is before movement onset, which would not shrink the neural activity in delay period due to orthogonality of preparatory and movement activities. The neural activity of other time point (100-ms bin) that aligned to TO were projected into the above 2D TDR subspace. To demonstrate the evolving path for neural states in target location TDR plane, the base (TO+0.1 s) of each of the four reach directions condition were set as the origin, and the location of other time point relative to it were computed. For hand reach location subspace, the location of each time point is computed relative to the actual origin of the plane (see **Fig. 5c-e, Supplementary Fig. 13**).

### Neural decoding

To assess the information encoded in neural data, a ridge regression model is trained to predict reach direction or target location from neural population activity. Specifically, PETH (20-ms bin with 25-ms Gaussian kernel smoothing, GO-1000 ms to GO+500 ms) were organized into a predictor matrix of dimensions *K*×*T* × *N* (trials ×times× neurons). The corresponding instantaneous target locations or final reach directions (in x- and y-axis components) were organized into a response vector of dimensions *K*×*T* × 2. The predictor matrix and response vector were divided into training and testing sets using 10-fold cross-validation. We applied ridge regression with L2 regularization (*fitrlinear* function in Matlab). A grid of λ values, ranging from 10^−5^ to 10^5^, was explored to identify the optimal regularization strength. The best lambda was selected based on the minimum mean squared error (MSE) obtained through cross-validation within the training set. For each fold and time point, separate models were trained and tested for x- and y-axis components. To evaluate the performance of the model, we computed three metrics: coefficient of determination *R*^*2*^ for x- or y-axis components, and cosine similarities for angular direction computed from x and y components. The cosine similarity was calculated as the dot product between predicted and true angular direction, normalized by their magnitudes. To assess the significance of the model’s predictions, shuffled labels of x and y were also used to train and evaluate the model. Wilcoxon rank sum test was performed to compare the performance of the true and shuffled models for each metric (see **Fig. 5f**).

### Input-driven neural network model

We adapted the stability-optimized circuits (SOC) neural network model with optimal feedback control ^35^ to simulate the motor cortical activity in our task.

### Network dynamics

We model motor cortex as a network consisting of 100 excitatory (E) neurons and 100 inhibitory (I) neurons. We constructed the synaptic architecture, *W*, following the methodology outlined by Hennequin et al. (2014)^57^, and utilizing MATLAB codes introduced by Stroud et al. (2018)^58^. Briefly, with an initial spectral abscissa of 10, a connection density of 0.1, an inhibition/excitation ratio of 3, and a gradient-descent learning rate of 10, we iteratively updated the inhibitory synapses of a random network to minimize a measure of robust network stability. We implemented early stopping, terminating the stabilization procedure as soon as the spectral abscissa of the connectivity matrix dropped below 0.15.

We modeled neuronal activity according to Equations (1-4), which we integrate using the *ode45*function (with default parameters) in MATLAB.

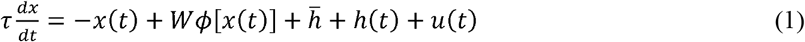

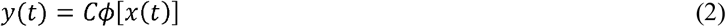

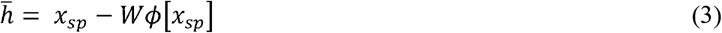

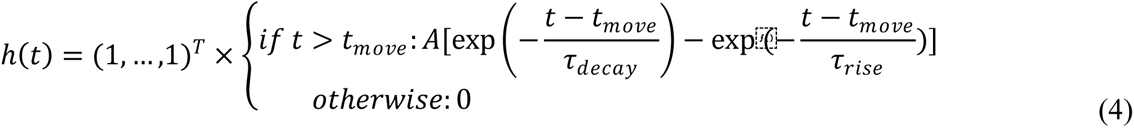

where *x*(*t*)=(*x*_*E*_(*t*)^*T*^ *x*_*I*_(*t*)^*T*^)^*T*^ is the internal state, τ is the single-neuron time constant, *W* is the synaptic connectivity matrix, and ϕ (*x*)= *max* (*x*, 0) is a rectified-linear nonlinearity that converts internal states into firing rates. 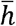 provides the baseline activity of a set of spontaneous firing rates *x*_sp_ ∼ *N*(20,2t; *h*(*t*) is a condition-independent spatially uniform-shaped input bump that is given during movement; *u*(*t*) is the preparatory control input that drive the circuit into the appropriate state for each movement. *y*(*t*) represents the hand position driven by the motor network (*u*=0), where *C*∈ ℝ^2×N^ is the readout matrix.

### Initial state and readout matrix optimization

We optimized a set of twelve initial conditions 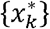, as well as the readout matrix, to calibrate the network for the generation of desired hand trajectories 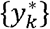, where *k* = 1, …,12. The trajectories were straight reaches of length *d*= 20 cm from the origin (0,0) into different directions, characterized by a bell-shaped speed profile *v*(*t*) given by Equation (5):

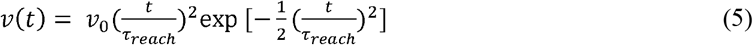

where *v*0 is adjusted to ensure that the hand reaches the target. To minimize the loss function (Equation 6), we used the MATLAB function *fminunc* with default parameters:

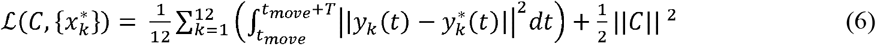

The loss function comprises two terms: the first term minimizes the squared difference between actual and desired hand trajectories, while the second term penalizes the squared Frobenius norm of the readout matrix *C*. *y* depends on *C* and 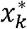 implicitly through the dynamics of the network (Equation 1–4).

For the initial conditions, 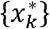, we assembled them using the top eigenvectors of the observability Gramian (a symmetric positive-definite matrix Q) of the linearized system, as described by Equation 7-8:

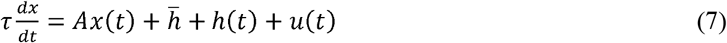

with state transition matrix *A* ≜*W*−*I* and Q is the solution to the Lyapunov equation

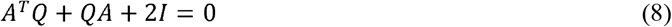

These initial conditions were linearly transformed using the top eigenvectors as the transformation matrix, achieved by performing 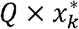. Additionally, the assembled 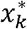 are normalized to satisfy a specific norm constraint.

Furthermore, we parameterized the readout matrix *C* to ensure that its null-space contains both the spontaneous activity vector *x*_*sp*_ and the movement-specific initial conditions 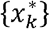. This ensures that (i) there is no muscle output during spontaneous activity and (ii) the network does not unduly generate muscle output before movement. Specifically, we enforced *Cϕ*[*x*_*sp*_]=*C x*_*sp=0*_ and 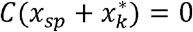.

### Control cost function and classical LQR solution

Assuming linear dynamics throughout the preparatory and movement epochs as Equation 7, we defined our total control cost functional as:

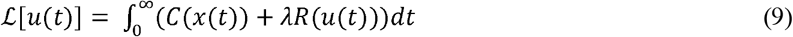

where *C* (*x*(*t*)) is the “prospective motor error”, i.e., the total error in movement that would result if movement was initiated at time *t. R* x(*u*(t)) represents an energetic cost penalizing large control signals beyond a baseline required to maintain *x*(*t*) in the optimal subspace, and sets the relative importance of this energetic cost. Regarding the impact of on the model’s performance, referred to Extended Data Fig. 7f. The cost functional can be specified as:

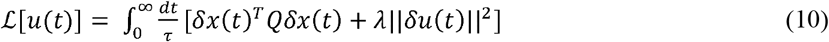

where Δ*x*(*t*)≜ *x*(t) −*x*^*^, and Δ*u*(*t*)≜*u*(t) −*u*^*^. *Q*is the observability Gramian of pair (*A,C*) and it is the solution to the Lyapunov equation:

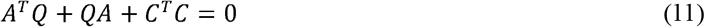

According to the linear quadratic regulator (LQR), the optimal control input:

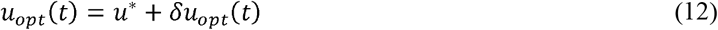

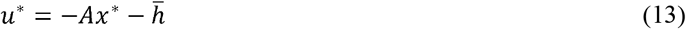

can be rewritten as

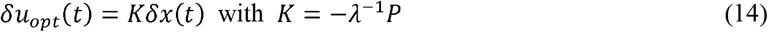

where *P* is a symmetric, positive-definite matrix, obtained as the solution to the Riccati equation:

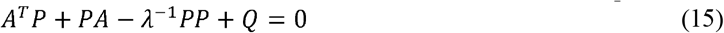

It has been verified that Equation 10 provides a good approximation to the actual *Cx*(*t*) resulting from nonlinear dynamics during the movement epoch ^35^. Thus, for simplicity, we adopted the classical LQR solution as our optimal feedback input.

The total control cost is given by 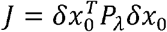, where δ*x*_0_ is the deviation of the network state from *x*^*^ at the onset of movement preparation (i.e., the state of the network at preparation onset is *x*^*+^ δ*x*_0._ The corresponding total energy cost ∫ *Rdt* is given by 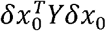, where Y is the solution to

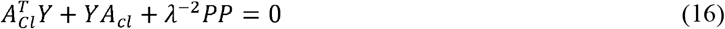

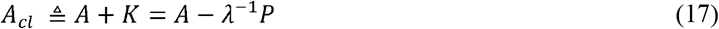

The associated integrated prospective motor cost is then given by

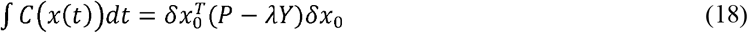

### Modeling the ICMS effect

To simulate the ICMS effect on the network, we introduced a perturbation input *h*_*pt*_ to a randomly selected subset of the neurons (*N*_*pt*_ =100) for a duration of 100 ms. The perturbation onset occurred 200 ms before movement onset to allow sufficient time for error correction. This timing ensures that errors are not entirely corrected before movement onset. The amplitudes of *h*_*pt*_ (*amp*_*pt*_ =0, 20, 40, or 50) were chosen to adequately reflect the differences in response to perturbation between the static and moving conditions.

To model the ICMS effect on reaction time, we adopted the proposal from the optimal subspace hypothesis, which posits that movement would not be initialized unless errors are minimized^11^. Accordingly, we assessed the time at which the network could generate a movement trajectory of high goodness of fit. We extended the preparatory period until 1 sec after TO, and at every 20 ms interval, we computed *R*^2^ of movement trajectory if movements were initiated at that time. We set a motor potent threshold (*R*^2 >^0.9) to determine an appropriate movement generation, and the first time at which *R*^2^ exceeds this threshold defines the movement onset *t*_*mo*_. Correspondingly, *RT*= *t*_*mo*_ − *t*_*delay*._

### Eye movements recording and analysis

During earlier ICMS experiments with monkey G (**Supplementary Table 3**), we recorded eye movements. Eye position signals were recorded at 1000 Hz using the EyeLink system (SR Research; EyeLink1000), known for its noninvasive infrared eye-tracking capabilities. An infrared “hot” mirror positioned at a 45° angle in front of the monkey facilitated eye tracking while allowing direct vision of the stimulus. Calibration of the eye tracker involved projecting a sphere (“target”) onto the display at one of the nine positions forming a 3 × 3 grid at the center of the screen. The monkey was required to fixate on the target sphere and maintain gaze for a random interval of 100–400 ms. The target sphere provided the calibration offset, and the voltage changes between targets provided the scaling factor for converting eye-tracker voltage to screen distance^102,103^.

To determine the eye movement reaction times (RTs), we first applied a low-pass filter (<50 Hz) to smooth out noise in the eye velocity signal. The first and second highest peaks in the filtered velocity signal were then identified as candidates. The peaks that meet the following criteria will be given priority as eye movement onset: (1) only peaks occurring after GO-200ms are considered; (2) if one peak occurs before Go and one after, the later peak is selected; (3) if both peaks occur before or after the GO, the peak closest to GO is selected. This ensures that eye movement onset is as late as possible after this Go, despite some potential bias. The relative time of the chosen peak to GO was recorded as the RT. Trials in the static and moving (target velocity of −120°/s) conditions, with delay-period length > 800 ms across 35 sessions, were selected for comparison (see **Fig. 7b, c**).

To compute the saccade probability during each trial, we first identified peaks in the filtered eye velocity signal. The five highest peaks were selected if they met threshold criteria (300-1000°/s) and their values and times relative to GO were stored. Trials in the static and moving (±120°/s target velocity) conditions, with delay period > 800 ms across 35 sessions, were analyzed. Saccades occurring within specific epochs were used to compute normalized probability for comparing conditions: from GO-900 ms to GO+100 ms between the static and moving conditions, and from GO-100 ms to GO+100 ms between NS and ST trials. For each histogram bin, a contingency table was constructed comparing counts within the current bin to the remaining bins, and a one-tailed Fisher’s exact test was applied for both right and left tails with an alpha level of 0.05 (see **Fig. 7d, g**).

To calculate the metric cosθ, we determined the direction of each saccade by computing two vectors: the eye movement direction 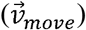, defined from the preceding time point to the peak of each eye velocity saccade, and the center vector 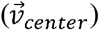, extending from the eye position at the peak to the origin. The cosine similarity (cosθ) between 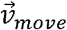 and 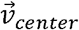 was then calculated to assess alignment of the eye movement with the center. Values close to +1 suggest that the saccade is directed towards the center, whereas values around −1 suggest it is directed towards the peripheral (see **Fig. 7f, h**).

## Data availability

The data supporting the findings of this study, along with the corresponding codes, are available on GitHub (https://github.com/zhengcong26/Interception_ICMS). Source data are provided with this paper.

## Code availability

All code related to this study is publicly available on GitHub (https://github.com/zhengcong26/Interception_ICMS).

## Acknowledgements

We are grateful to Yongxiang Xiao for invaluable discussions; to Ruichen Zheng, Tianwei Wang and Siwei Xie for surgical expertise and experiment advice; to Chao Guan and Yizhi Lu for expert veterinary care; to Tianlin Luo for preliminary works in network modeling; to Abdulraheem Nashef, Daniel J. O’Shea, and Amy Osborn for insightful advices; to Yun Chen, Chenyang Li, Joseph Malpeli, and Yiheng Zhang for helpful comments on the manuscript. This work was supported by National Key R&D Program of China (Grant 2020YFB1313402, Grant 2017YFA0701102); National Science Foundation of China (Grant 31871047, Grant 31671075); Shanghai Municipal Science and Technology Major Project (Grant 2021SHZDZX) to He Cui, and Shanghai Postdoc Fellowship Funding (Y95CN51621) to Cong Zheng.

## Contributions

C.Z., and H.C. conceived and designed the experiments. C.Z. and Q.W. performed the experiments. C.Z. analyzed data. C.Z. wrote the paper with inputs from all authors.

