## Supplementary Information for "Continuous sensorimotor transformation enhances robustness of neural dynamics to perturbation in macaque motor cortex"

Supplementary Methods

Supplementary Table 1 Session information for monkey G and L-1

Supplementary Table 2 Session information for monkey G and L-2

Supplementary Table 3 Session information (trial number) for eye movement analysis (monkey G)

Supplementary Figure 1. Experiment setup and launching angles

Supplementary Figure 2. EMGs, current amplitudes effect, reach trajectories post-ICMS, and electrode locations.

Supplementary Figure 3. Controlled analyses for post-ICMS reaction time (RT) changes in monkey G.

Supplementary Figure 4. Controlled analyses for post-ICMS reaction time (RT) changes in monkey L.

Supplementary Figure 5. Post-ICMS reaction times (RTs) changes in sessions with random conditions.

Supplementary Figure 6. Stimulation-aligned changes in the odds of initiating versus not initiating a reach.

Supplementary Figure 7. Reach kinematics after ICMS of monkey G.

Supplementary Figure 8. Reach kinematics after ICMS of monkey L.

Supplementary Figure 9. Neural trajectories in alternative reach directions corresponding to Fig. 3d, e

Supplementary Figure 10. individual neural activity examples corresponding to Fig. 4e.

Supplementary Figure 11. Preferred direction (PD) analyses

Supplementary Figure 12. The impact of the number of neurons on the tortuosity of neural trajectories

Supplementary Figure 13. Neural activity in target-location and reach-direction TDR plane of monkey G and L

Supplementary Figure 14. Supplementary neural decoding results and demixed PCA analyses.

Supplementary Figure 15. Preparatory and movement–subspace occupancy.

Supplementary Figure 16. Optimal states of the moving condition in model.

Supplementary Figure 17. Features of the input-driven model.

Supplementary Figure 18. Control results for the probability of saccade.

**Supplementary Methods**

**Bootstrapping approach to calculate differences in reaction times (RTs)**

For each bootstrap iteration, we randomly sampled RTs from the NS condition, paired them with the ST trials, and computed the difference in RTs between the ST and the paired NS trials. for NS condition, we also randomly sampled from the NS group itself, paired these sampled trials with the remaining NS, and computed the reaction time differences within the NS condition. We performed n = 10000 bootstrap resamples for each condition (reach directions). The median of the RT differences across all bootstrap iterations was then calculated (see **Supplementary Figure 3, 4**).

**Bootstrapping approach to examine the impact of number of neurons**

To examine the whether a small number of neurons influences the tortuosity of neural trajectory observed in **Figure 5a, b**, we applied a bootstrap approach to recompute the detour index. For each bootstrap iteration (neuron Bootstraps), a random selection of 10 neurons was drawn from the available neural units. The sample size was chosen because it represents the minimum number of neurons recorded in any session within our dataset. The selection was performed using the *randperm* function to ensure that each neuron was chosen without replacement. For each selected subset of neurons, trials were resampled (trial Bootstraps). Neural trajectories and detour indices were then computed for each neuron and trial bootstrap. We performed 10,000 (100 × 100) bootstrap resamples on each dataset, averaging across these bootstrap samples (see **Supplementary Figure 12**).

**Impact of microstimulation on reach initiation probability**

To examine whether the RT effects across conditions can be explained by a stimulation-aligned change in the odds of initiating to not initiating a reach, we employed the methodology proposed by Zimnik et al. (2019)^1^. Specifically, we calculated the hazard rate of reach initiation, or the probability of initiating a reach at time *t* given that initiation had not occurred before *t*, $P_{0}(t)$, as follows:

$$P_{0}\left( t \right)=P\left( T_{m}\in\left[ t,t+150 \right] \right| T_{m}\geq t)$$

where $T_{m}$ is the recorded time of movement onset. $P_{0}(t)$ is computed by taking all trials where movement was initiated after time t and computing the proportion initiated within the interval $\left[ t,t+150 \right]$. To obtain a reliable estimate, $P_{0}(t)$ was estimated only for values of t where there were at least 80 trials where initiation had not yet occurred by time t. The probability of initiation following stimulation is defined as follows:

$$P_{stim}\left( t,t_{s} \right)=P\left( T_{m}\in\left[ t,t+150 \right] \right| T_{m}\geq t,T_{s}\in\left[ t_{s},t_{s}+150 \right])$$

where $T_{s}$ is the time of stimulation. $P_{0}\left( t \right)$ and $P_{stim}\left( t,t_{s} \right)$ were calculated every 10 ms. $P_{stim}\left( t,t_{s} \right)$ was estimated only for values of $t$ and $t_{s}$ that yielded at least 20 observations. The change in the probability of moving is defined as follows:

$$\Delta P_{stim}\left( t,t_{s} \right)= P_{stim}\left( t,t_{s} \right)-P_{0}\left( t \right)$$

See **Supplementary Figure 6**.

**Demixed PCA**

To demix and analyze the neural representation of individual variables, we employed demixed principal components analysis (dPCA), a dimensionality reduction technique established by Kobak et al. 2016 ^2^ [https://github.com/machenslab/dPCA]. dPCA decomposes population activity into individual neural dimensions, each capturing variance related to specific task variables, such as reach directions. First, we organized the binned firing rate data into a trial-averaged, four-dimensional tensor, $x_{ndst}$, where $n$ is the number of neurons, $d$ is the number of reach direction conditions, $s$ is the number of target speed conditions, and $t$ is the number of time-points. Then, we decomposed $x_{ndst}$ into a set of averages over various combinations of parameters (marginalization). The original trial-averaged data tensor can be expressed as the sum of all the marginalizations: $\tilde{x}_{tds}=\bar{x}_{tds}+{\bar{x}_{ds}+\bar{x}}_{ts}+\bar{x}_{td}+\bar{x}_{s}+\bar{x}_{d}+\bar{x}_{t}+\bar{x}$. We grouped the marginalizations [$\bar{x}_{d}$, $\bar{x}_{td}$] for reach direction and selected the dPCA component with the largest explained variance for demonstration (see **Supplementary Figure 14**).

**Identifying preparatory and movement dimensions**

To assess differences in preparatory activity between the static and moving conditions, we computed the variance (“subspace occupancy”) in preparation and execution dimensions of each condition. This was achieved using the dimensionality reduction approach from Lara et al., (2018)^3^. Specifically, neural activity was first smoothed using a 20-ms bin size and a 20-ms standard deviation Gaussian kernel, soft-normalized (normalization factor = firing rate range + 5), and then mean-centered at each time by subtracting mean activity across all conditions of each neuron at each time point from each condition’s response. To estimate confidence intervals, we performed a bootstrap resampling of trials to generate surrogate datasets. Then, we defined preparatory matrices $P \in R^{N\times CT}$ using data from TO+60 ms to TO+900 ms and the movement matrices $M \in R^{N\times CT}$using data from MO-40 ms to MO+200 ms, where *N* is the number of neurons, *C* is the number of reach directions, and *T* is the number of time points. The method seeks a set of preparatory dimensions or subspace bases, $W_{prep}$, to capture the maximum variance of *P*, and an orthogonal set of movement dimensions, $W_{move}$ , to capture the maximum variance of *M*. For this purpose, the following objective function is optimized:

$$\left[ W_{prep},W_{move} \right]={argmax}_{\left[ W_{prep},W_{move} \right]}\frac{1}{2}(\frac{Tr\left( W_{prep}^{T}C_{prep}W_{prep} \right)}{\sum_{i=1}^{d_{prep}} \sigma_{prep}\left( i \right)}+\frac{Tr\left( W_{move}^{T}C_{move}W_{move} \right)}{\sum_{i=1}^{d_{move}} \sigma_{move}\left( i \right)})$$

*Subject to* $W_{prep}^{T}W_{move}=0, W_{prep}^{T}W_{prep}=I,$ $W_{move}^{T}W_{move}=I$

Where $C_{prep}=cov\left( P \right)$, $C_{move}=cov\left( M \right)$, $\sigma_{prep}\left( i \right)$ and $\sigma_{move}\left( i \right)$ is the $i^{th}$ singular value of $C_{prep}$ and $C_{move}$, respectively. $Tr\left( W_{prep}^{T}C_{prep}W_{prep} \right)$ and $Tr\left( W_{move}^{T}C_{move}W_{move} \right)$ is the preparatory-epoch and movement-epoch data variance captured by preparatory and movement subspace, respectively. The dimensionality of $W_{prep}$ was set to 30, capturing over 70% of preparatory variance, and dimensionality of $W_{move}$ to 10, capturing over 80% of movement variance.

For a given time *t* and condition $\theta$, the projection of the population response onto the $k^{th}$ preparatory dimension is a weighted sum of across neurons: $x_{n}^{prep}\left( t,\theta\right)=$ $\sum_{n=1}^{N} W_{n,k}^{prep}r_{n}(t,\theta)$ where $W_{n,k}^{prep}$ is the element in the $n^{th}$ row and $k^{th}$ column of $W_{prep}$ and $r_{n}\left( t,\theta\right)$ is the response of the $n^{th}$ neuron. Measurement of subspace occupancy is equivalent to the variance explained metric. For the preparatory subspace, occupancy was computed as: ${occupancy}^{prep}\left( t \right)=\sum_{k=1}^{30} {var}_{\theta}(x_{k}^{prep}(t,\theta))$ where ${var}_{\theta}$ is taking the variance across reach direction condition. Movement subspace occupancy was defined analogously. We normalized the variance values using min-max scaling to range between 0 and 1. To assess differences between the static and moving conditions, subspace bases were constructed using data from the static condition, the moving condition, or both (see **Supplementary Figure 15**).

**Table 1. Session information for monkey G and L-2**

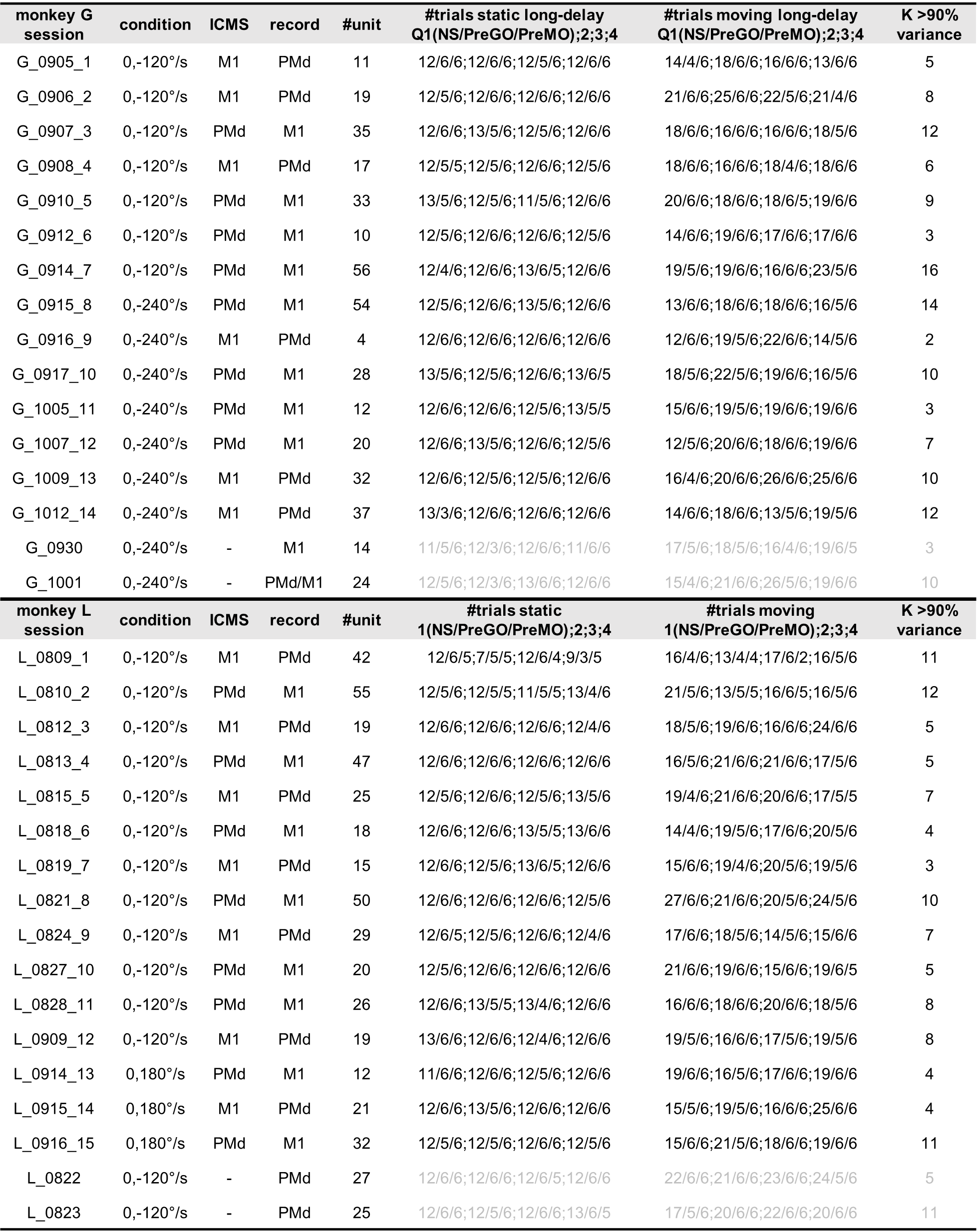

**Table 2. Session information for monkey G and L-2**

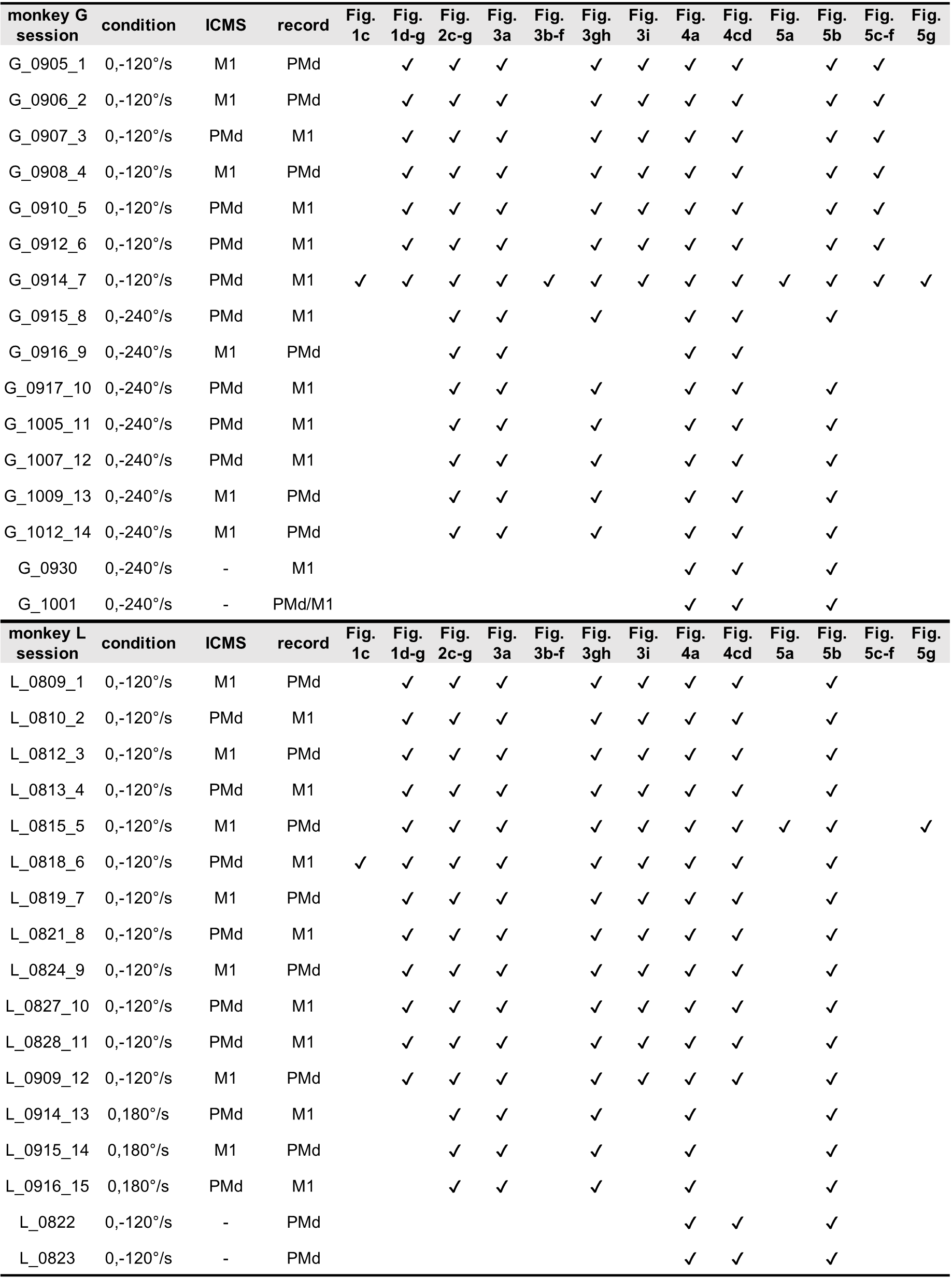

**Table 3. Session information (trial number) for eye movement analysis (monkey G)**

| Session | static NS | static ST | moving NS* | moving ST* | moving ST*  PreGO | moving ST*  PreMO |
| --- | --- | --- | --- | --- | --- | --- |
| 0915 | 14 | 16 | 139 | 64 | 0 | 32 |
| 0916 | 14 | 16 | 180 | 63 | 4 | 32 |
| 0917 | 19 | 16 | 170 | 64 | 5 | 32 |
| 0918 | 16 | 16 | 171 | 64 | 5 | 32 |
| 0920 | 8 | 16 | 139 | 64 | 3 | 32 |
| 0922 | 12 | 16 | 160 | 64 | 3 | 32 |
| 0923 | 13 | 16 | 153 | 64 | 2 | 32 |
| 0924 | 7 | 16 | 119 | 59 | 11 | 31 |
| 0926 | 18 | 16 | 165 | 64 | 10 | 32 |
| 0927 | 9 | 16 | 120 | 64 | 11 | 32 |
| 0928 | 16 | 16 | 132 | 64 | 10 | 32 |
| 0929 | 21 | 16 | 151 | 64 | 4 | 32 |
| 1008 | 15 | 10 | 81 | 46 | 3 | 22 |
| 1009 | 13 | 16 | 132 | 62 | 9 | 31 |
| 1011 | 12 | 16 | 136 | 64 | 3 | 32 |
| 1012 | 7 | 9 | 56 | 31 | 10 | 21 |
| 1013 | 6 | 14 | 103 | 60 | 28 | 29 |
| 1014 | 9 | 14 | 76 | 50 | 5 | 23 |
| 1019 | 13 | 13 | 92 | 53 | 4 | 28 |
| 1025 | 5 | 6 | 47 | 25 | 4 | 9 |
| 1026 | 11 | 11 | 95 | 48 | 6 | 15 |
| 1027 | 17 | 14 | 103 | 51 | 3 | 15 |
| 1101 | 30 | 16 | 158 | 63 | 8 | 16 |
| 1102 | 42 | 16 | 219 | 64 | 8 | 16 |
| 1114 | 31 | 16 | 138 | 64 | 7 | 16 |
| 1115 | 11 | 15 | 95 | 60 | 4 | 14 |
| 1117 | 47 | 16 | 161 | 64 | 12 | 16 |
| 1119 | 38 | 16 | 196 | 63 | 8 | 16 |
| 1120 | 14 | 14 | 80 | 52 | 7 | 14 |
| 1121 | 40 | 16 | 166 | 64 | 16 | 16 |
| 1122 | 26 | 15 | 111 | 63 | 4 | 15 |
| 1123 | 39 | 16 | 185 | 63 | 14 | 15 |
| 1124 | 32 | 16 | 178 | 64 | 4 | 16 |
| 1125 | 30 | 15 | 175 | 63 | 8 | 16 |
| 1126 | 20 | 16 | 106 | 64 | 10 | 16 |

* sp$\neq$0°/s, delay >800 ms

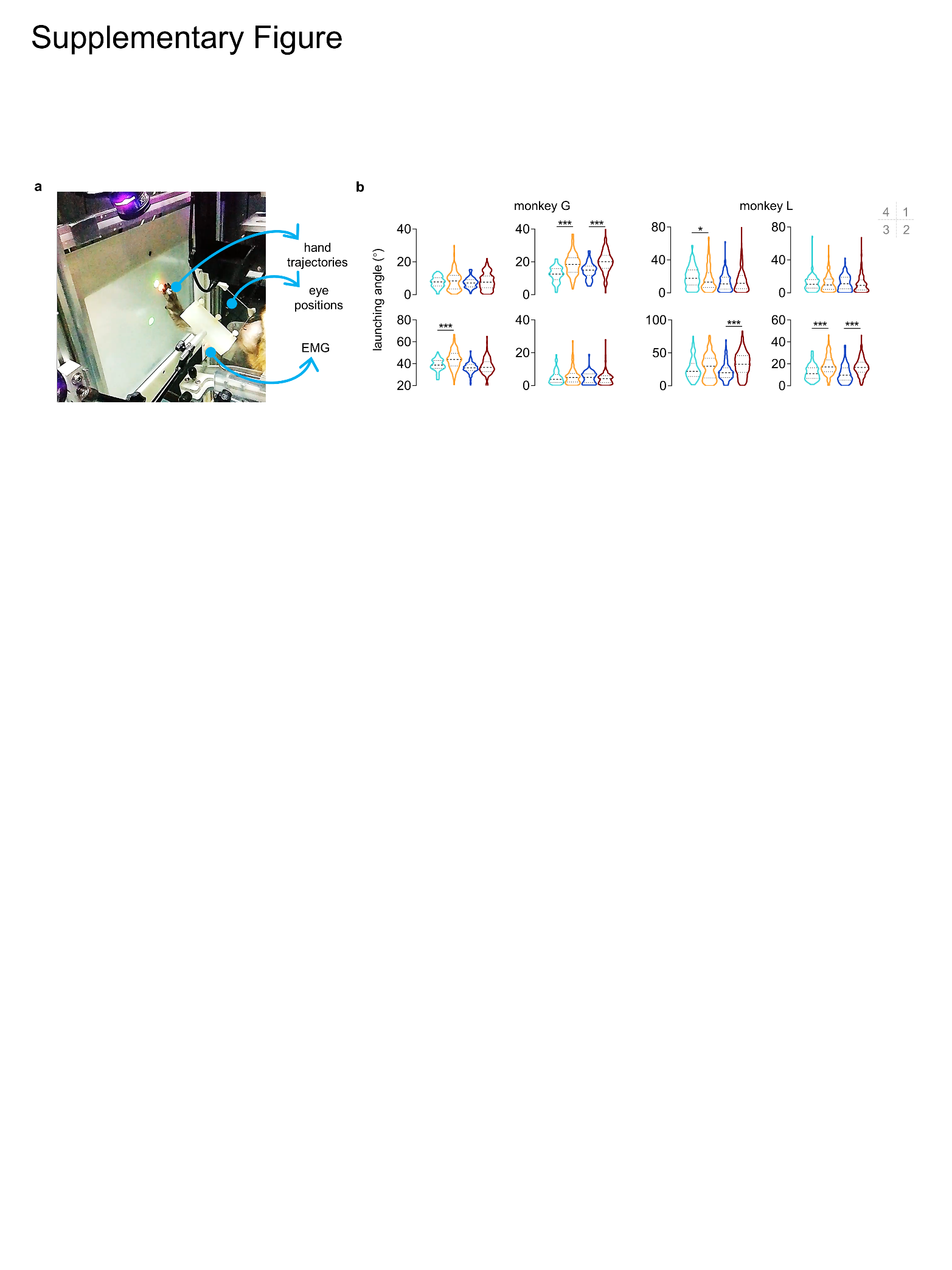

**Supplementary Figure 1. Experiment setup and launching angles**

**a.** Experimental setups. Monkey faced to a vertical touch screen while their eye movements were monitored using Eyelink system equipped with a reflecting mirror. Hand movement trajectories were recorded with a reflecting ball attached to the middle finger using Vicon motion capture, and muscle activities was captured by a wireless sensor from Delsys affixed to the forearm.

**b.** Launching angles relative to endpoints. Data for monkey G (73$\pm$5, 156$\pm$18, 68$\pm$3, and 169$\pm$18 trials for 1-4 reach direction) and monkey L (131$\pm$10, 305$\pm$33, 131$\pm$4, and 293$\pm$29 trials). Subplots correspond to these four reach directions. Data are displayed as medians (dashed lines) and quartiles (dotted lines). Asterisks indicate significance levels from Wilcoxon rank sum test with *p < 0.05, ***p < 0.001.

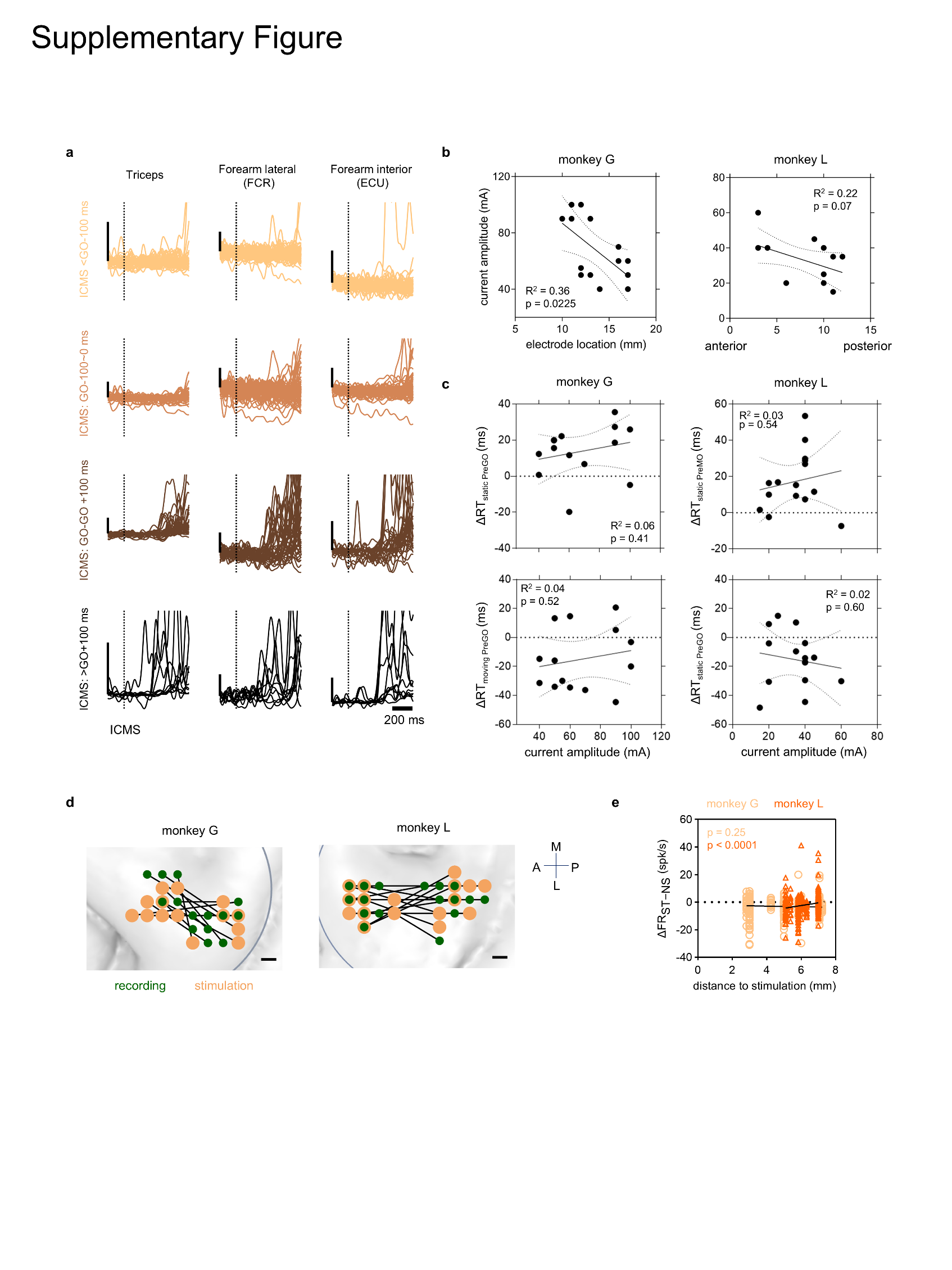

**Supplementary Figure 2. EMGs, current amplitudes effect, reach trajectories post-ICMS, and electrode locations.**

**a.** Single-trial surface electromyography (sEMG) data were recorded using a wireless sensor (Delsys Trigno Lab). The EMG data underwent band-pass filtering (6th order Butterworth, 35-40 Hz), rectification (absolute value), and normalization to the range [0, 1]. Mean muscle activity for one session is displayed, aligned to different ICMS timings. Trials: n = 87, 84, 36, 10.

**b.** ICMS current amplitudes as a function of electrode locations. Dotted lines indicate the 95% CIs for a linear fit. Goodness of fit (*R^2^*) and p-value for the non-zero slope are annotated.

**c.** The effect of ICMS as a function of current amplitudes. Annotations are consistent with panel **c**.

**d.** Electrode locations for ICMS and S-probes used for simultaneous recording. Each pair corresponds to a session. The nearest distance between sites is 1 mm. The mean distances between stimulating and recording electrodes were 4.7 mm (min 2.8 mm - max 7.2 mm) for monkey G and 6.2 mm (min 3.6 mm - max 7.0 mm) for monkey L.

**e.** Changes in firing rates (ΔFR = $\text{FR}_{\text{ST}}$ – $\text{FR}_{\text{matched NS}}$) as a function of distance of recorded units to stimulation sites. N = 368 for monkey G and N = 430 for monkey L. p value of non-zero slope are annotated.

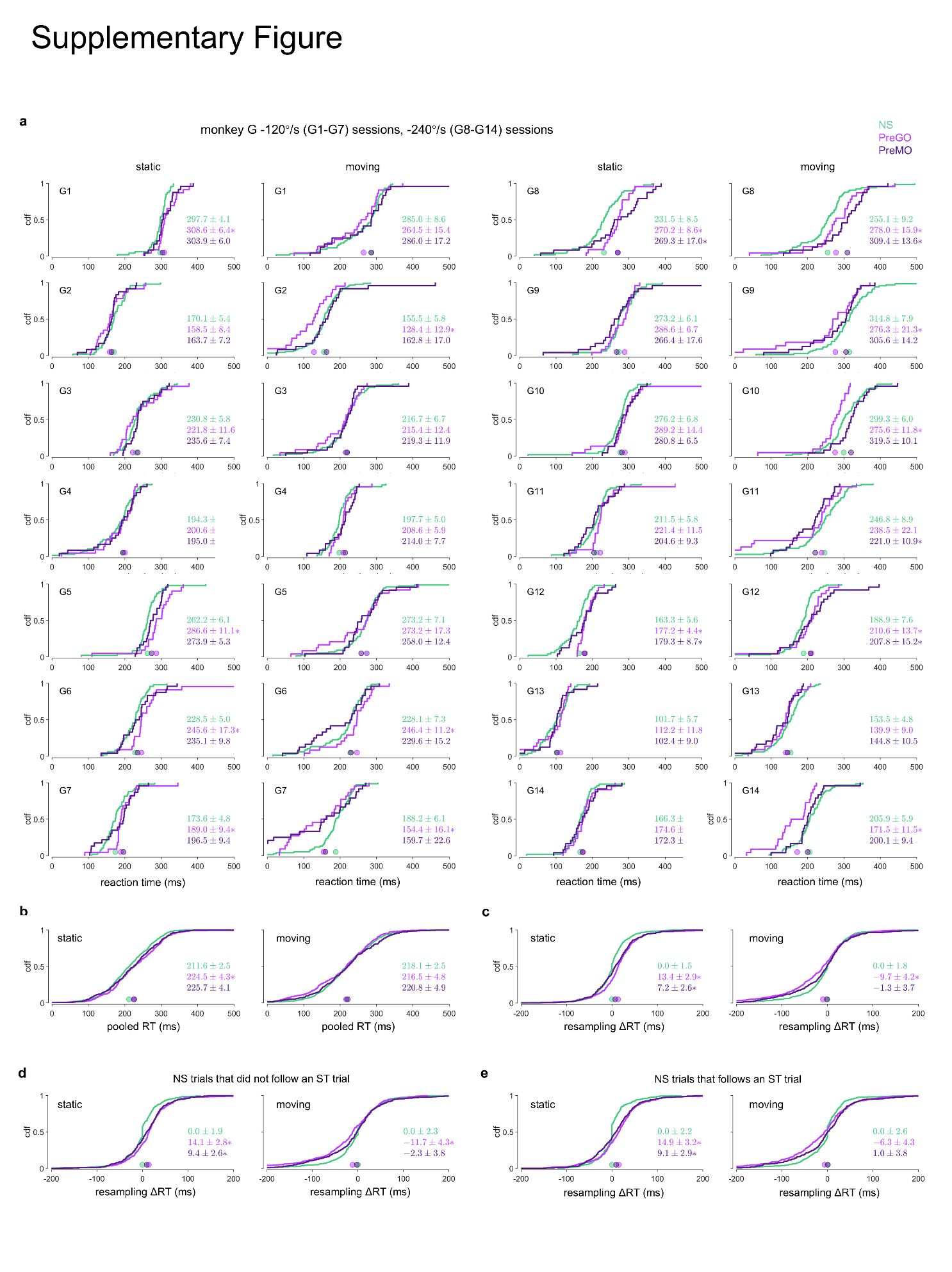

**Supplementary Figure 3. Controlled analyses for post-ICMS reaction time (RT) changes in monkey G.**

**a.** Cumulative distribution function (CDF) of RT distributions for individual sessions of monkey G. Median RT across all trials for each condition in one session is displayed, with significance of difference between NS and ST trials assessed using Wilcoxon rank sum test (*p < 0.05).

**b.** CDF of RT distributions pooled across all 14 sessions, with annotations identical to those in panel **a**.

**c.** CDF of resampled changes in reaction times (ΔRT = $\text{RT}_{\text{ST}}$ – $\text{RT}_{\text{matched NS}}$) pooled across all sessions, with annotations identical to those in panel **a**.

**d.** CDF of resampled ΔRT distributions for NS trials that did not follow an ST trial. Data are pooled across all sessions, with annotations identical to those in panel **a**.

**e.** CDF of resampled ΔRT distributions for NS trials that followed an ST trial. Data are pooled across all sessions, with annotations identical to those in panel **a**.

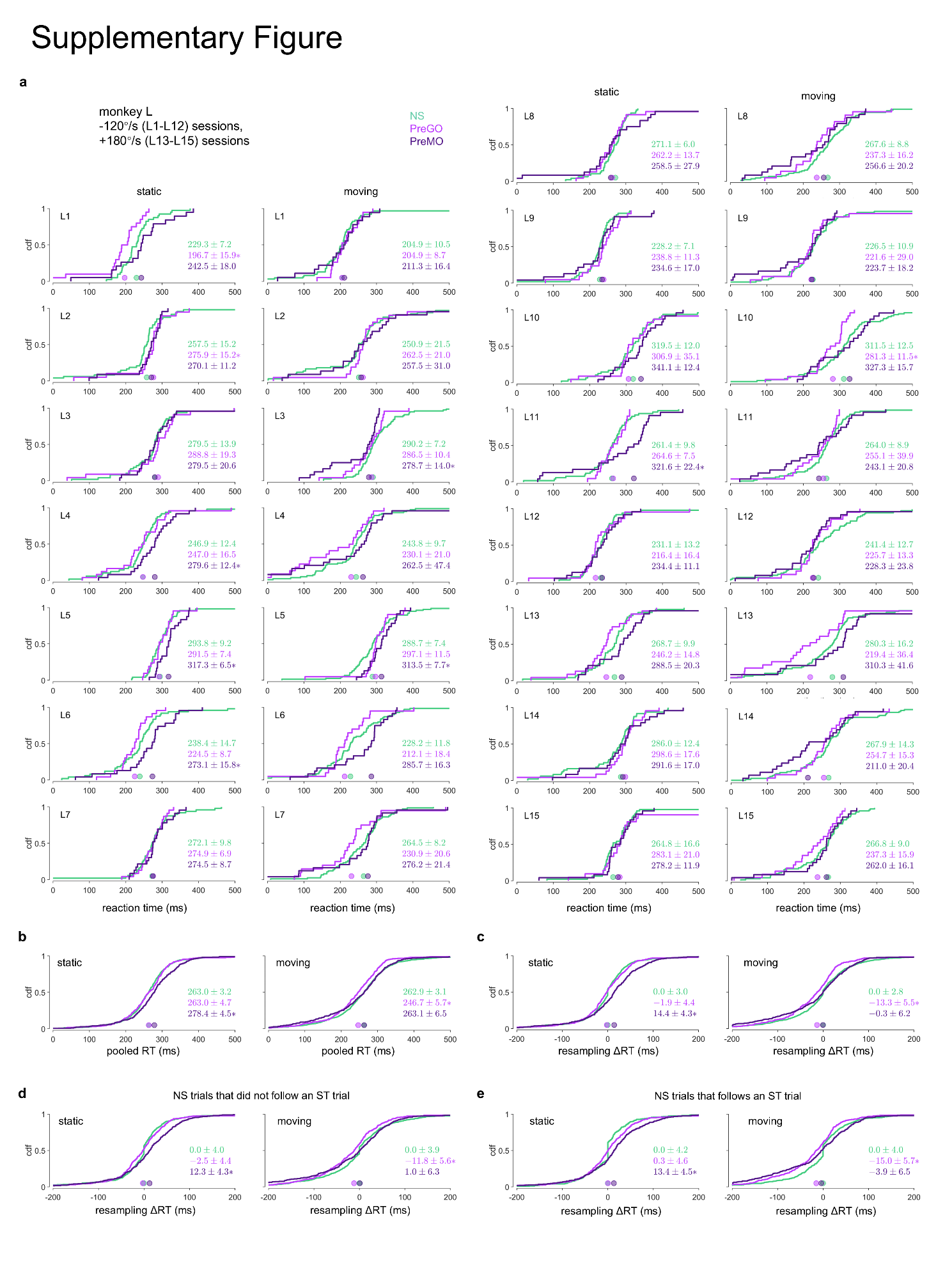

**Supplementary Figure 3. Controlled analyses for post-ICMS reaction time (RT) changes in monkey L.**

**a.** Cumulative distribution function (CDF) of RT distributions for individual sessions of monkey G. Median RT across all trials for each condition in one session is displayed, with significance of difference between NS and ST trials assessed using Wilcoxon rank sum test (*p < 0.05).

**b.** CDF of RT distributions pooled across all 15 sessions, with annotations identical to those in panel **a**.

**c.** CDF of resampled changes in reaction times (ΔRT = $\text{RT}_{\text{ST}}$ – $\text{RT}_{\text{matched NS}}$) pooled across all sessions, with annotations identical to those in panel **a**.

**d.** CDF of resampled ΔRT distributions for NS trials that did not follow an ST trial. Data are pooled across all sessions, with annotations identical to those in panel **a**.

**e.** CDF of resampled ΔRT distributions for NS trials that followed an ST trial. Data are pooled across all sessions, with annotations identical to those in panel **a**.

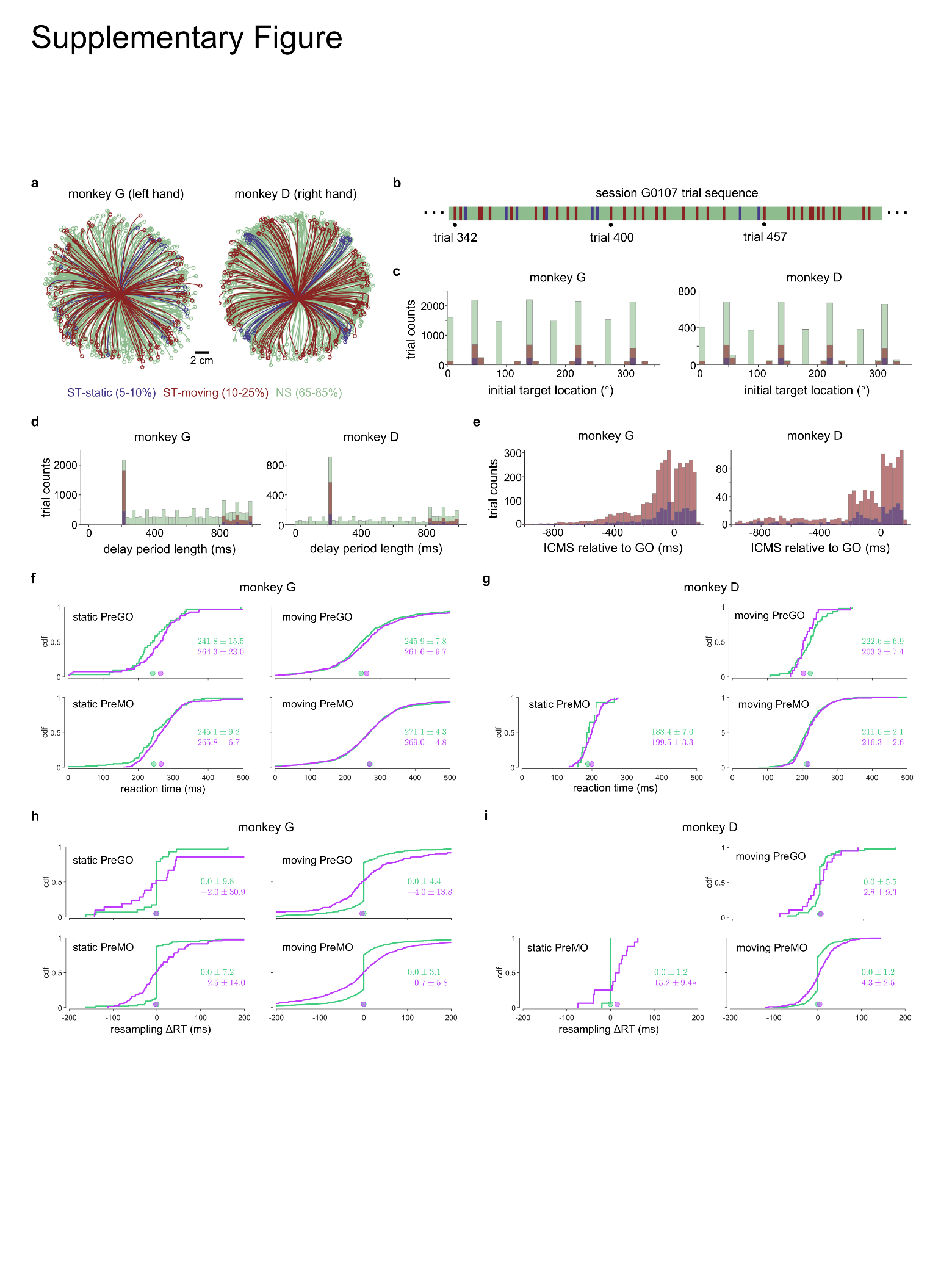

**Supplementary Figure 5. Post-ICMS reaction times (RTs) changes in sessions with random conditions.**

**a.** Reach trajectories for a representative session including random conditions (initial target locations, delay period lengths, and ICMS timings) for monkey G and D, comprising 927 and 675 trials, respectively.

**b.** Trial sequence for a session including random condition. Color coding corresponds to that in panel **a**.

**c. d. e.** Histogram of trial numbers for different initial target locations (**c**), l delay period lengths (**d**), and ICMS timings (**e**). Data are pooled across all sessions. Color coding corresponds to that in panel **a**. Trials: n = 15999 (NS), 931 (static), and 3698 (moving) across 32 sessions for monkey G; n = 4670 (NS), 288 (static), and 1143 (moving) across 9 sessions for monkey D.

**f. g.** Cumulative distribution function (CDF) of RT distributions pooled across all random sessions for monkey G (**f**) and monkey D (**g**). The median RT for each condition in one session is displayed, with significance of differences between NS and ST trials assessed using Wilcoxon rank-sum test (*p < 0.05). The datasets are the same as those presented in panel **c**.

**h. i.** CDF of resampled changes in reaction times (ΔRT = $\text{RT}_{\text{ST}}$ – $\text{RT}_{\text{matched NS}}$) pooled across all random-condition sessions for monkey G (**h**) and monkey D (**i**). The datasets are consistent with those in panel **c**, with annotations alignning with those in panel **f**.

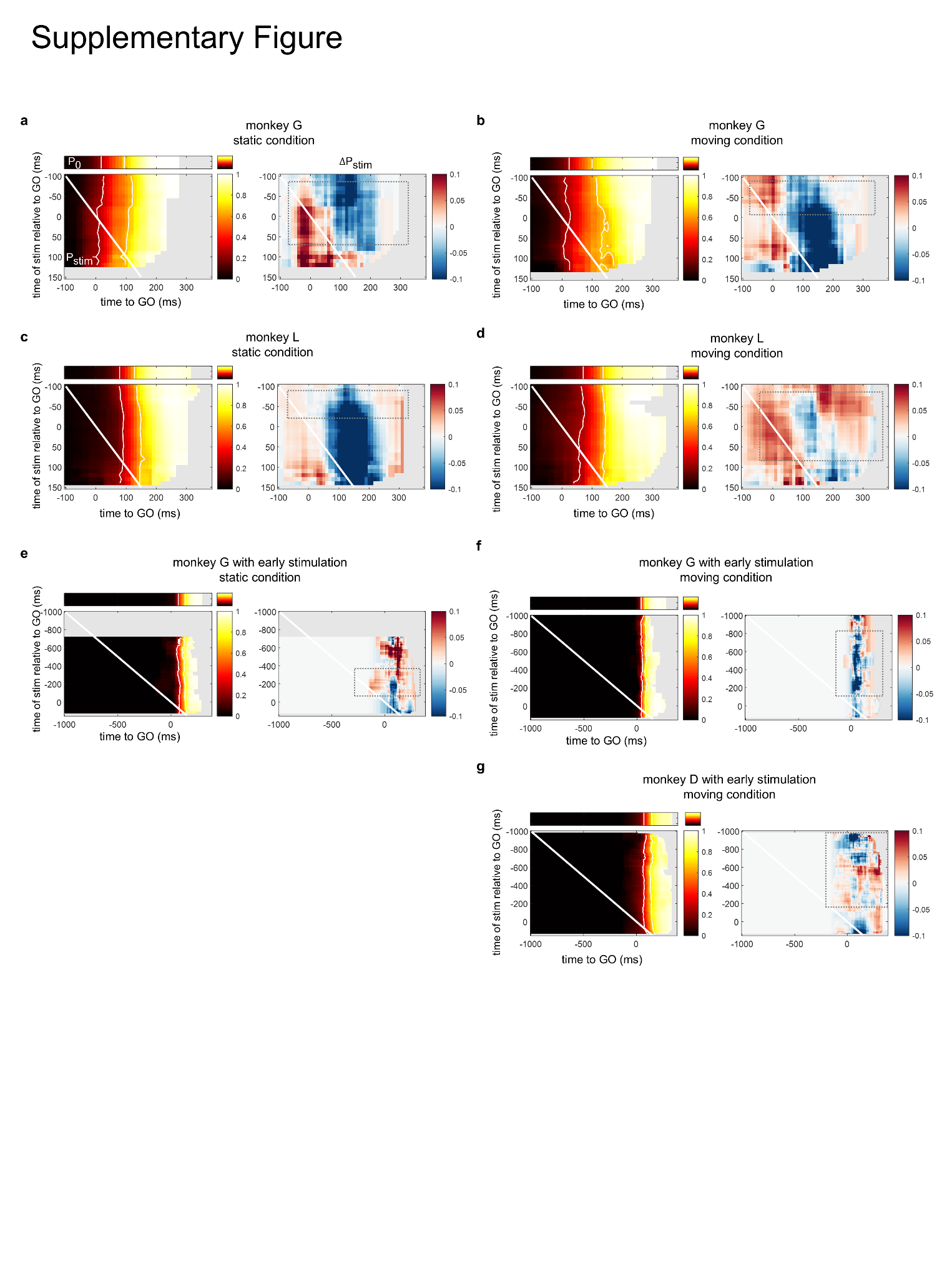

**Supplementary Figure 6. Stimulation-aligned changes in the odds of initiating versus not initiating a reach.**

**a. b.** Heatmap of $P_{0}$(top left) and $P_{stim}$(bottom left), and ${\Delta P}_{stim}$(right) for static (**a**) and moving (**b**) conditions of monkey G. For $P_{stim}(t,t_{s})$, each row represents the probability of initiation as a function of time relative to GO, for a specific time of stimulation. White lines indicate the onset of microstimulation for each row. White contour lines indicate *P* = 0.25 and *P* = 0.65, respectively. Gray dotted box highlights regions inconsistent with the biphasic ICMS impact observed in the SMA in prior study (Zimnik et al., 2019). Trials: n = 679 (static), 994 (moving) for non-stimulation condition, and n = 469 (static), 475 (moving) for stimulation condition.

**c. d.** Heatmap of $P_{0}$and $P_{stim}$, and ${\Delta P}_{stim}$for static (**c**) moving (**d**) conditions of monkey L. Trials: n = 562 (static), 848 (moving) for non-stimulation condition, and n = 531 (static), 524 (moving) trials for stimulation condition.

**e. f.** Heatmap of $P_{0}$and $P_{stim}$, and ${\Delta P}_{stim}$for static (**e**) moving (**f**) conditions of monkey G during sessions with random target initial locations and early stimulation. Trials: n = 328 (static), 3065 (moving) for non-stimulation condition, and n = 461 (static), 1879 (moving) for stimulation condition.

**g.** Heatmap of $P_{0}$and $P_{stim}$, and ${\Delta P}_{stim}$for the moving condition of monkey D during sessions with random target initial locations and early stimulation. The static condition of monkey D is not shown due to insufficient trial numbers in the preliminary task design. Trials: n = 1097 (moving) for non-stimulation condition, and n = 577 (moving) for stimulation condition.

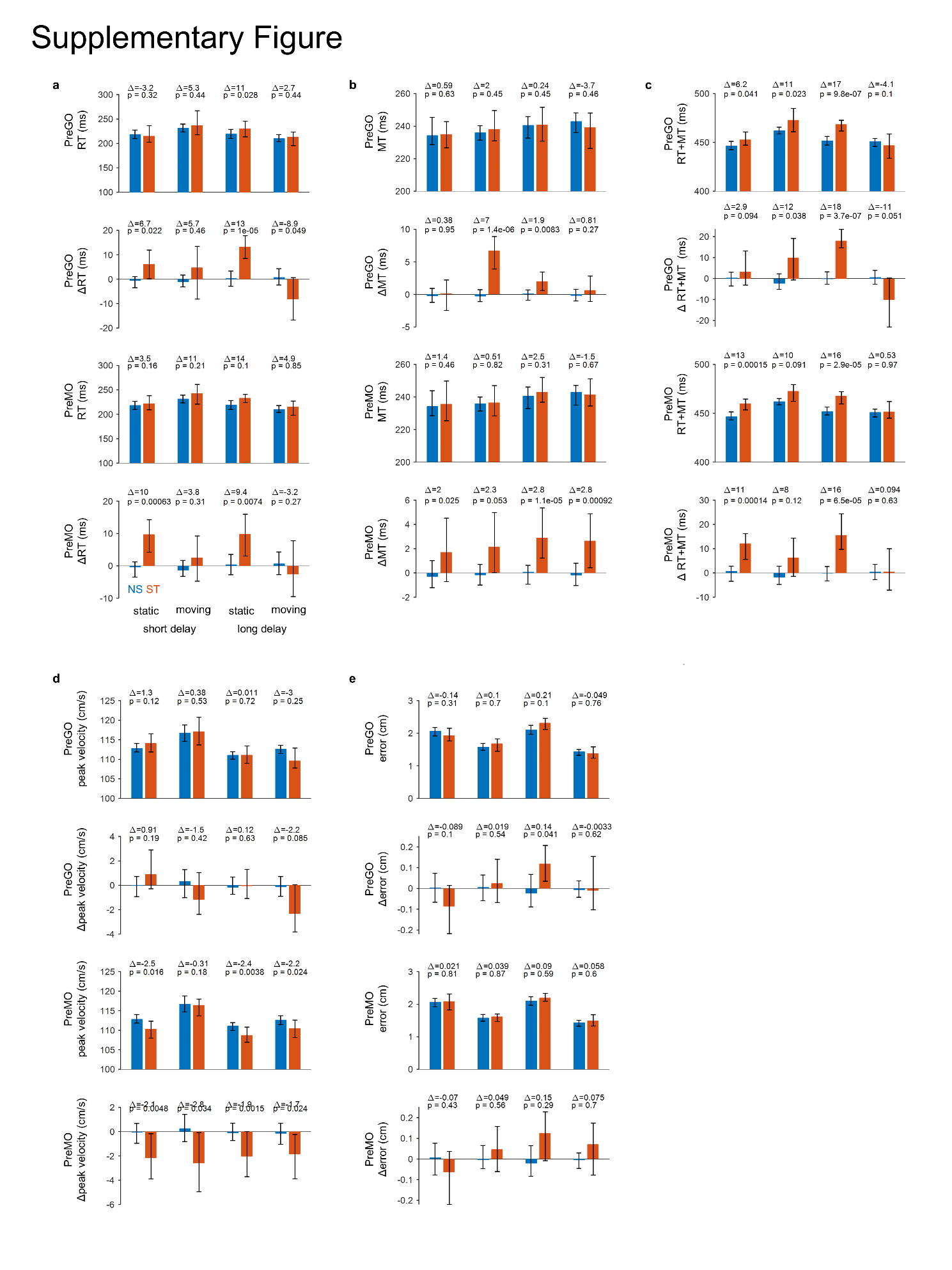

**Supplementary Figure 7. Reach kinematics after ICMS of monkey G.**

**a.** Reaction time (RT). Data are represented as median$\pm$95%CI for both raw data and de-NS data (stimulated minus median of condition-matched non-stimulated). Differences between NS and ST trials (Δ = ST-NS) are indicated above each condition, along with p-values from the Wilcoxon rank-sum test.

**b.** Movement time (MT). Annotations are consistent with panel **a**.

**c.** RT+MT. Annotations are consistent with panel **a**.

**d.** Peak hand velocity. Annotations are consistent with panel **a**.

**e.** Touchpoint errors. Annotations are consistent with panel **a**.

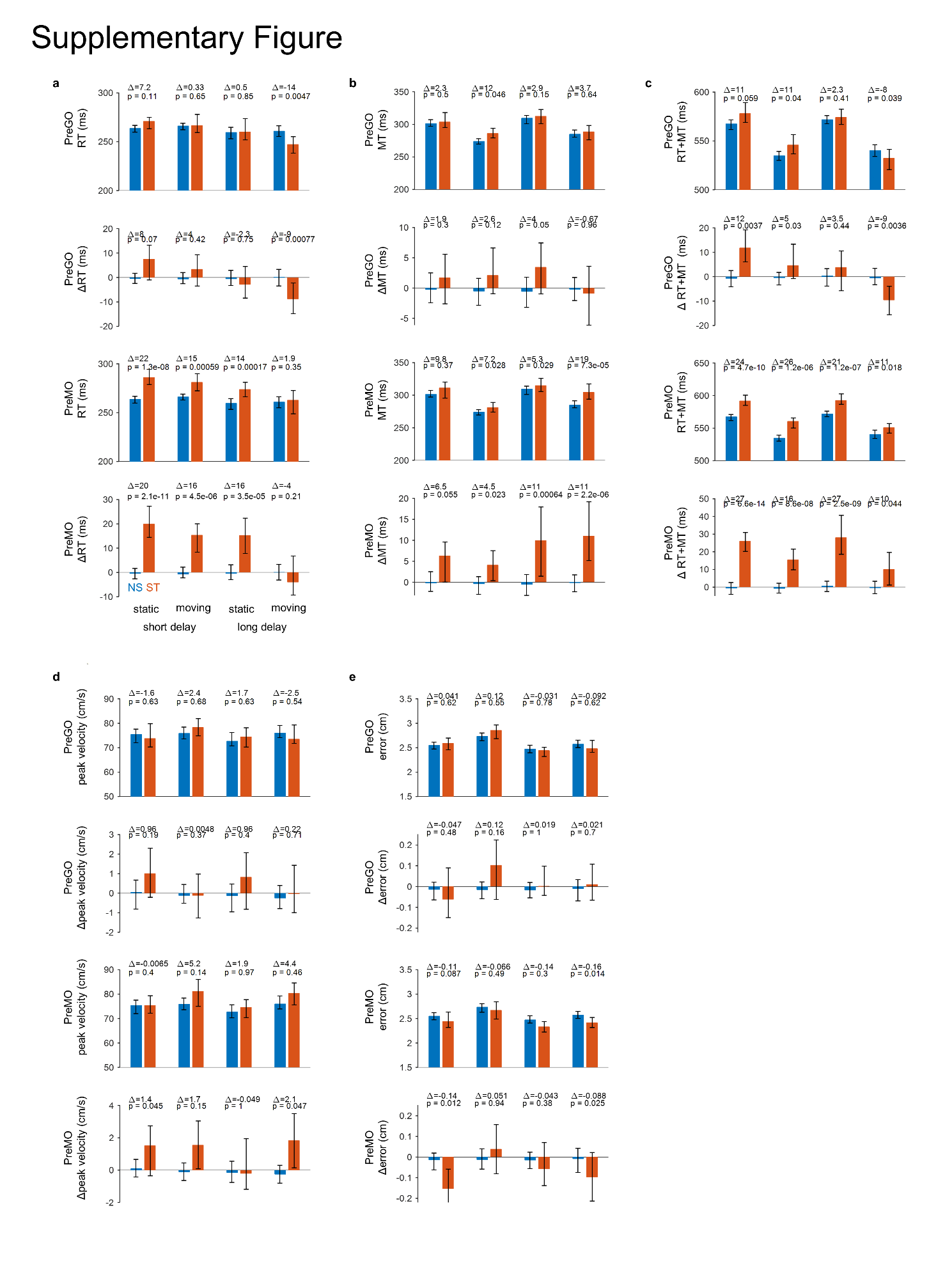

**Supplementary Figure 8. Reach kinematics after ICMS of monkey L.**

**a.** Reaction time (RT). Data are represented as median$\pm$95%CI for both raw data and de-NS data (stimulated minus median of condition-matched non-stimulated). Differences between NS and ST trials (Δ = ST-NS) are indicated above each condition, along with p-values from the Wilcoxon rank-sum test.

**b.** Movement time (MT). Annotations are consistent with panel **a**.

**c.** RT+MT. Annotations are consistent with panel **a**.

**d.** Peak hand velocity. Annotations are consistent with panel **a**.

**e.** Touchpoint errors. Annotations are consistent with panel **a**.

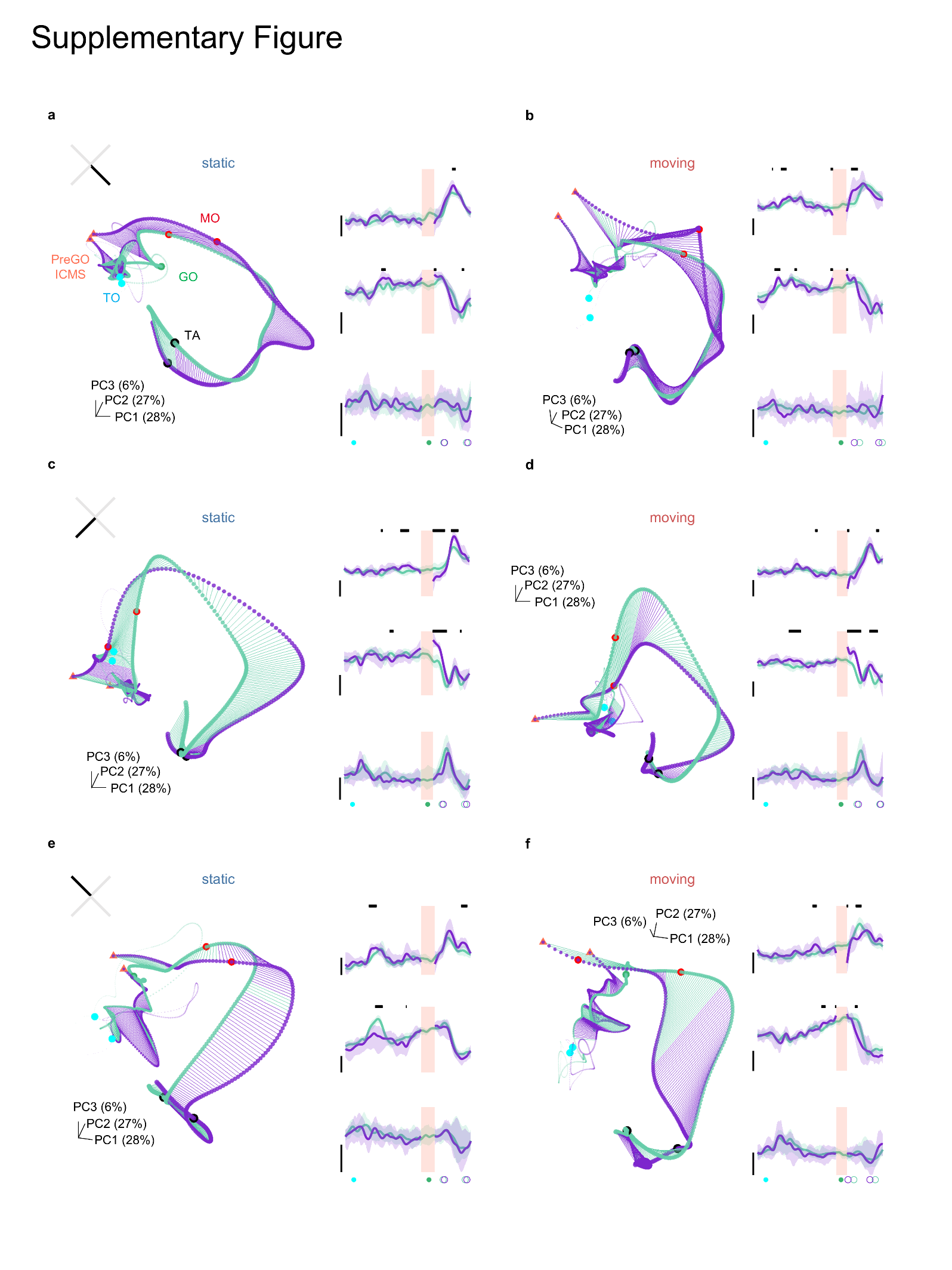

**Supplementary Figure 9. Neural trajectories in alternative reach directions corresponding to Fig. 3d, e**

**a. b.** Neural trajectories (right) and the first three PCs ($\pm$95%CI) (left) for static (**a**) and moving (**b**) conditions, with reaches directed toward quadrant 2. Annotations are consistent with those in **Fig. 3d, e**.

**c. d.** Neural trajectories and the first three PCs for static (**c**) and moving (**d**) conditions, with reaches directed toward quadrant 3.

**e. f.** Neural trajectories and the first three PCs of static (**e**) and moving (**f**) conditions, with reaches directed toward quadrant 4.

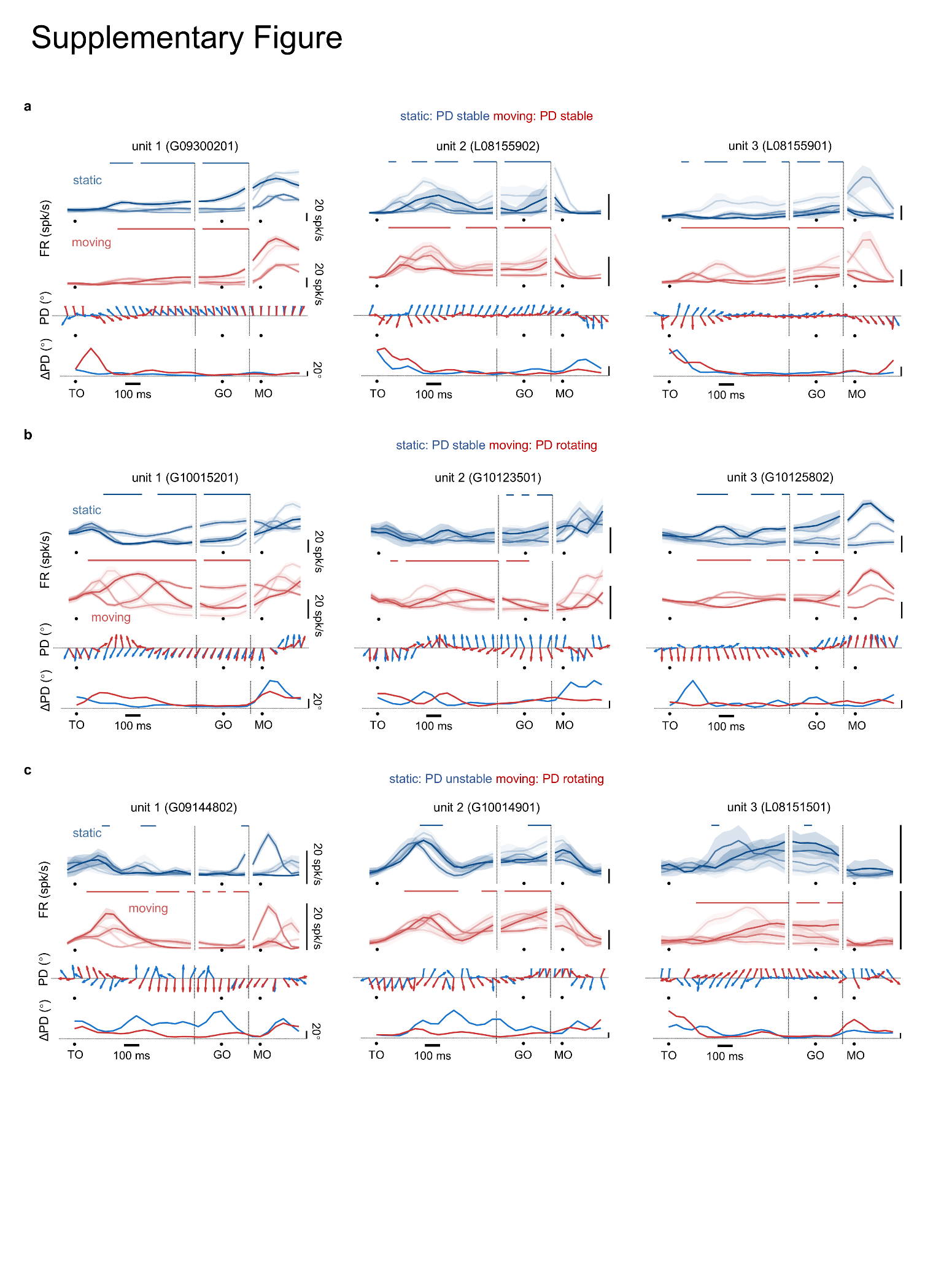

**Supplementary Figure 10. individual neural activity examples corresponding to Fig. 4e.**

**a.** Concatenated PETH (upper), preferred direction (PD, middle) and PD changes (ΔPD, bottom) for three example neurons exhibiting stable PD in both the static and moving conditions.

**b.** PETH, PD and ΔPD for three example neurons with stable PD in the static condition and rotating PD in the moving condition.

**c.** PETH, PD and ΔPD for three example neurons with unstable PD during the late delay in the static condition and rotating PD in the moving condition.

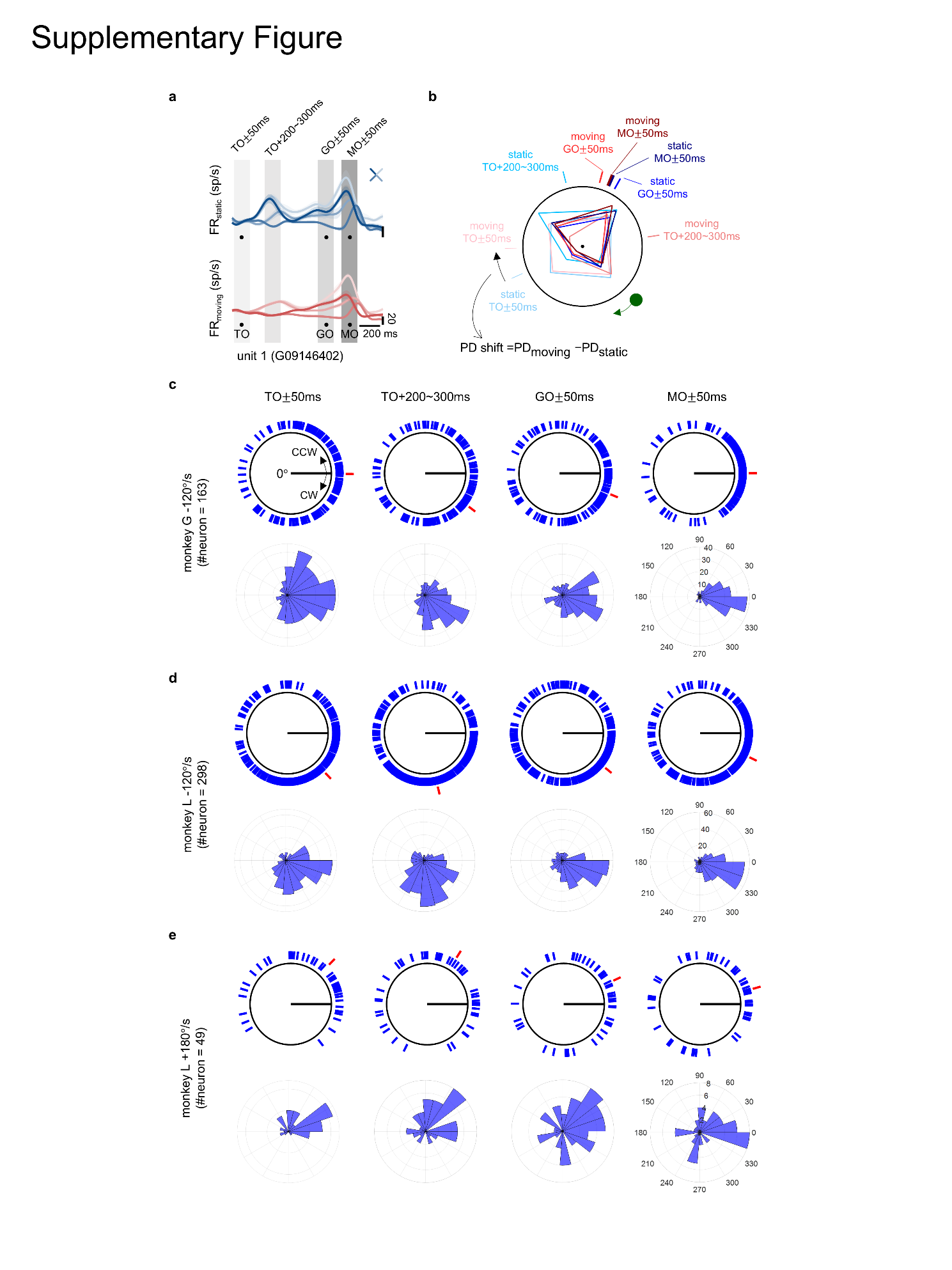

**Supplementary Figure 11. Preferred direction (PD) analyses**

**a.** The time epochs selected for PD analyses are shaded on PETH of an example neuron.

**b.** Polar plots illustrating the tuning function of the neuron shown in panel **a**. Colored lines outside the polar plots indicate the preferred direction axis for the corresponding condition at different times.

**c.** Population directional-bias plots for monkey G sessions with moving target velocity of -120°/s. For each neuron, PDs in the moving condition are plotted relative to PDs in the corresponding static condition, aligned to 0° in a polar plot (upper) and a polar histogram (bottom). Red lines indicate the median of PDs.

**d.** Population directional-bias plots for monkey L sessions with a moving target velocity of -120°/s.

**e.** Population directional-bias plots for monkey L sessions with a moving target velocity of +180°/s.

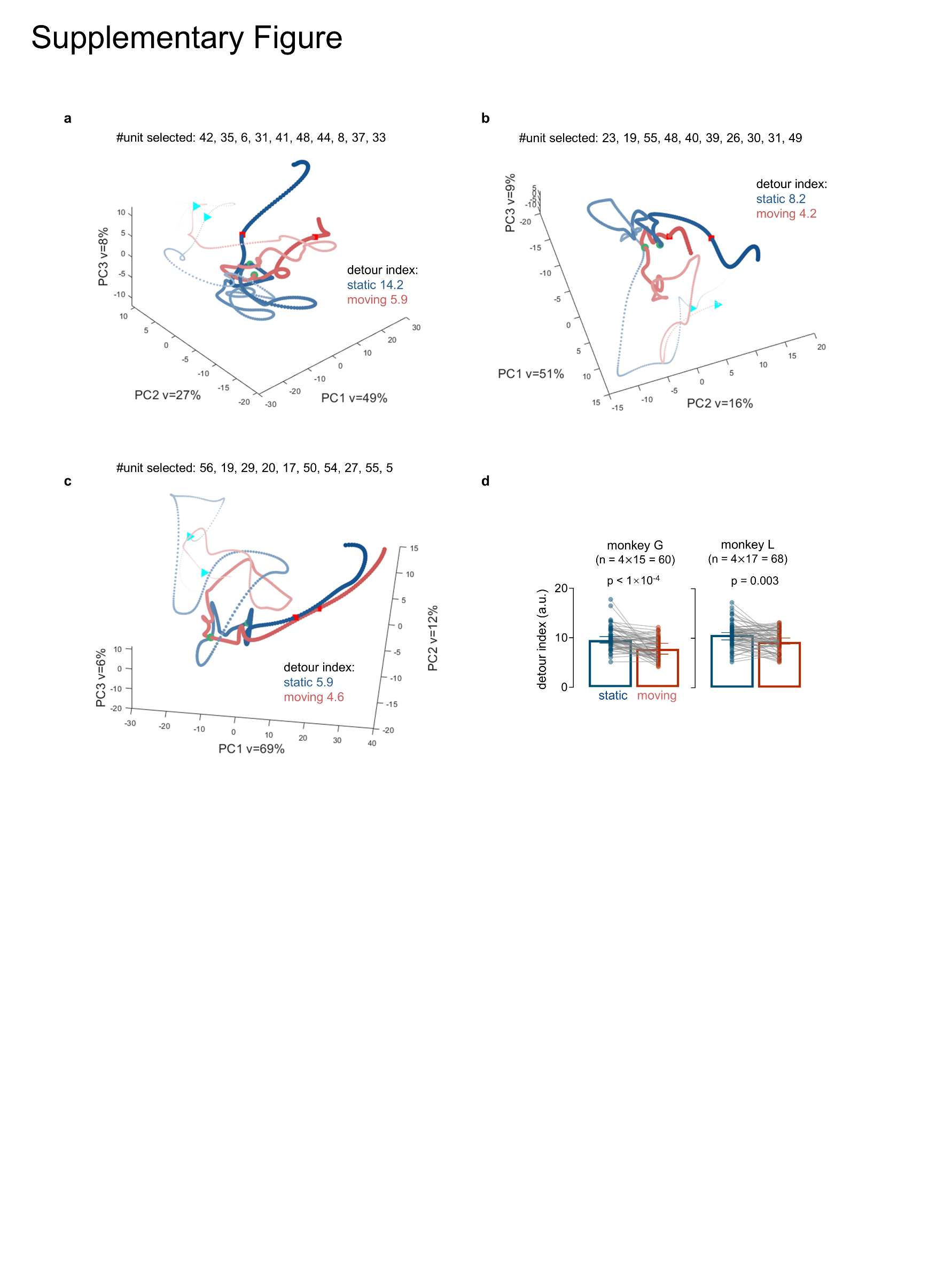

**Supplementary Figure 12. The impact of the number of neurons on the tortuosity of neural trajectories**

**a. b. c.** Three example neural trajectories computed from bootstrap samples. Ten neurons were randomly selected from a total of 56 neurons in the same dataset of monkey G in **Fig. 5a**. Despite some variance, the detour index remains higher in the static condition.

**b.** The detour index is computed through resampling of both neurons and trials. Differences between two conditions are assessed using the Wilcoxon matched-pairs signed rank test.

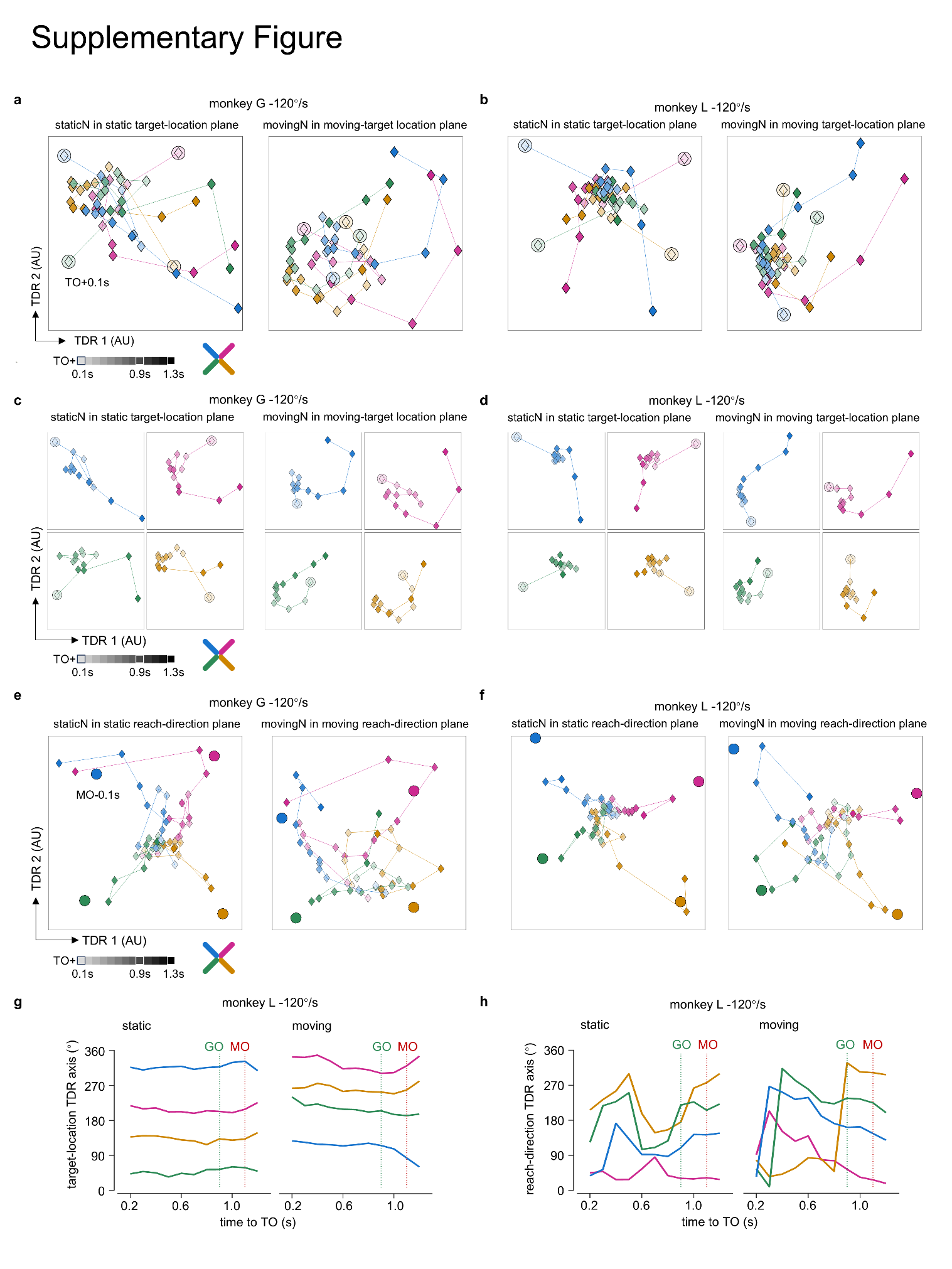

**Supplementary Figure 13. Neural activity in target-location and reach-direction TDR plane of monkey G and L**

**a. b.** Neural states in the moving condition over time within the target-location TDR subspace, constructed from neural activity in static (left) or moving (right) condition at TO+0.1 s (large filled circles). Data are averaged across seven sessions of monkey G with 181 neurons (**a**) and 14 sessions of monkey L with 416 neurons (**b**).

**c. d.** Neural states in the moving condition over time in target-location TDR subspace of monkey G (**c**) and monkey L (**d**), with four reach directions plotted separately. These data are the same as in panels **a** and **b**.

**e. f.** Neural states in the moving condition over time in reach-direction TDR subspace of monkey G (**e**) and monkey L (**f**), constructed from neural activity of the moving condition at MO-0.1 s (large filled circles). These data are the same as in panels **a** and **b**.

**g.** Time course of the relative change in position of neural states with respect to TO+0.1 s in static (left) and moving (right) target-location TDR subspace of monkey L. The data are the same as in panel **a**, **b**.

**h.** Time course of absolute change in neural state position in static (left) and moving (right) target-location TDR subspace of monkey L. Neural states in both conditions converged to similar relative positions by MO. The data are the same as in panel **f**.

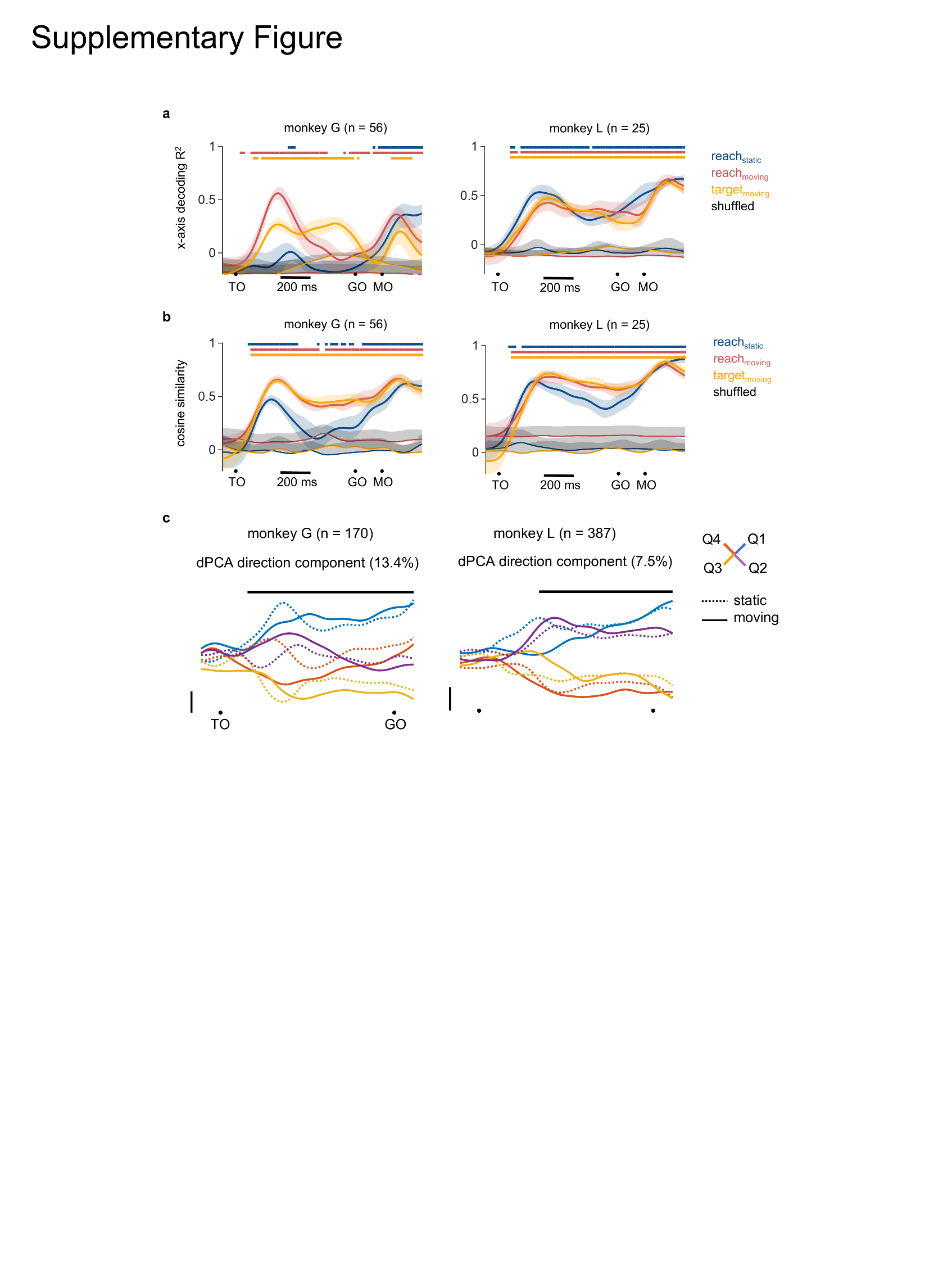

**Supplementary Figure 14. Supplementary neural decoding results and demixed PCA analyses.**

**a.** Goodness of fit *R^2^* ($\pm$95%CI) for linear decoding of reach directions (x-axis component) in static (reach_static_) and moving (reach_moving_) conditions, as well as instantaneous target location in the moving condition (target_moving_), derived from neural population activities in an example dataset shown in **Fig 5g**. Colored lines above denote significant differences from the shuffled level, determined by Wilcoxon rank-sum test (p < 0.05).

**b.** Cosine similarity of reach directions or target locations in degree for data in panel **a**.

**c.** Largest demixed principal components corresponding to reach direction. Thick black lines indicate time intervals during which the respective task parameters can be reliably extracted from single-trial activity. Explained variances are shown as percentages.

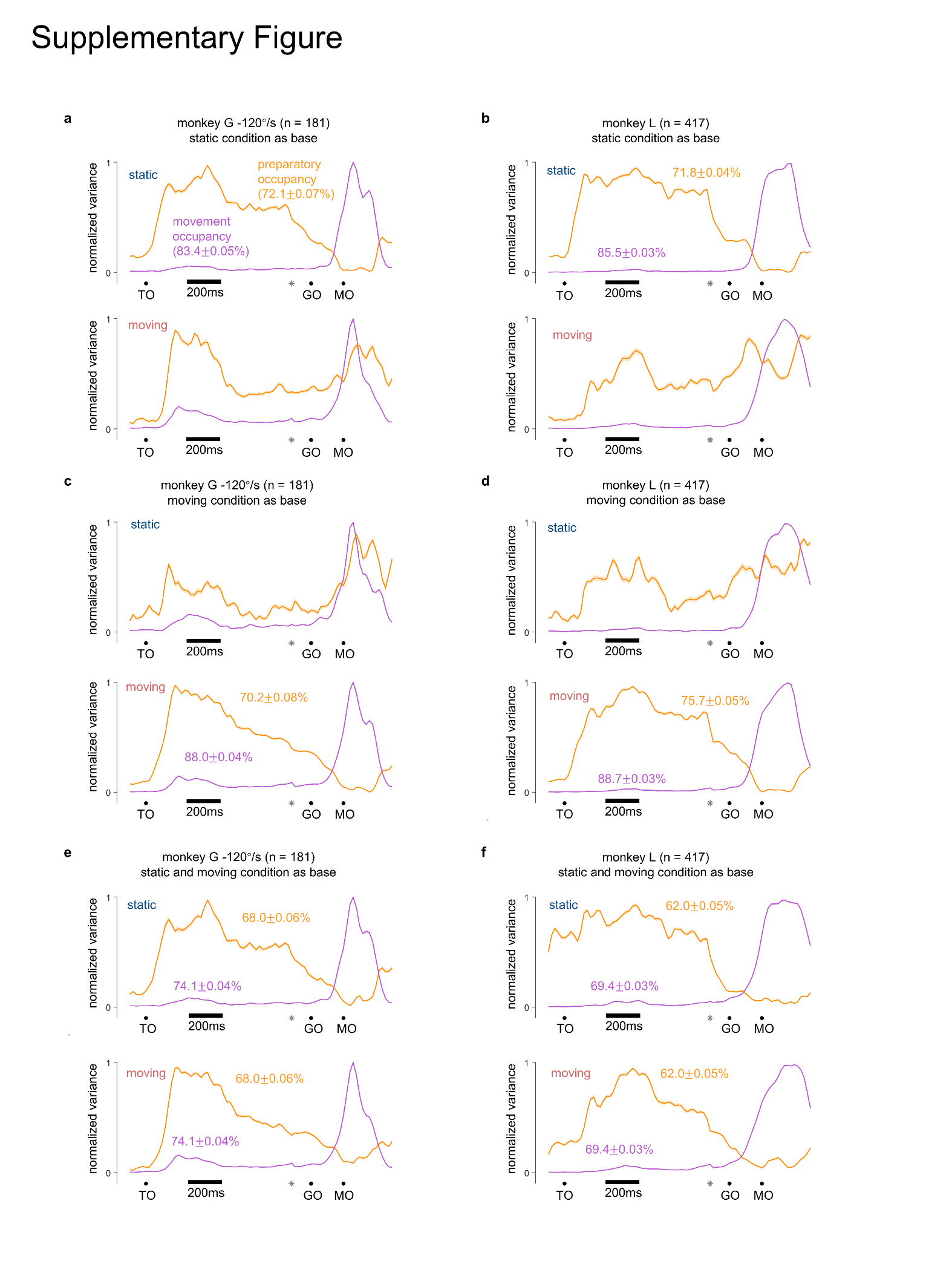

**Supplementary Figure 15. Preparatory and movement–subspace occupancy.**

**a. b.** Preparatory and movement subspace occupancy (mean$\pm$95%CI) of the static (upper) and moving (bottom) conditions, using the static condition as subspace base for monkey G (**a**) and monkey L (**b**). Gray symbols indicate the times when TO-aligned data and MO-aligned data were concatenated.

**c. d.** Preparatory and movement subspace occupancy (mean$\pm$95%CI) with the moving condition used as subspace base for monkey G (**c**) and monkey L (**d**).

**e. f.** Preparatory and movement subspace occupancy (mean$\pm$95%CI) with both the static and moving condition used as subspace base for monkey G (**c**) and monkey L (**d**).

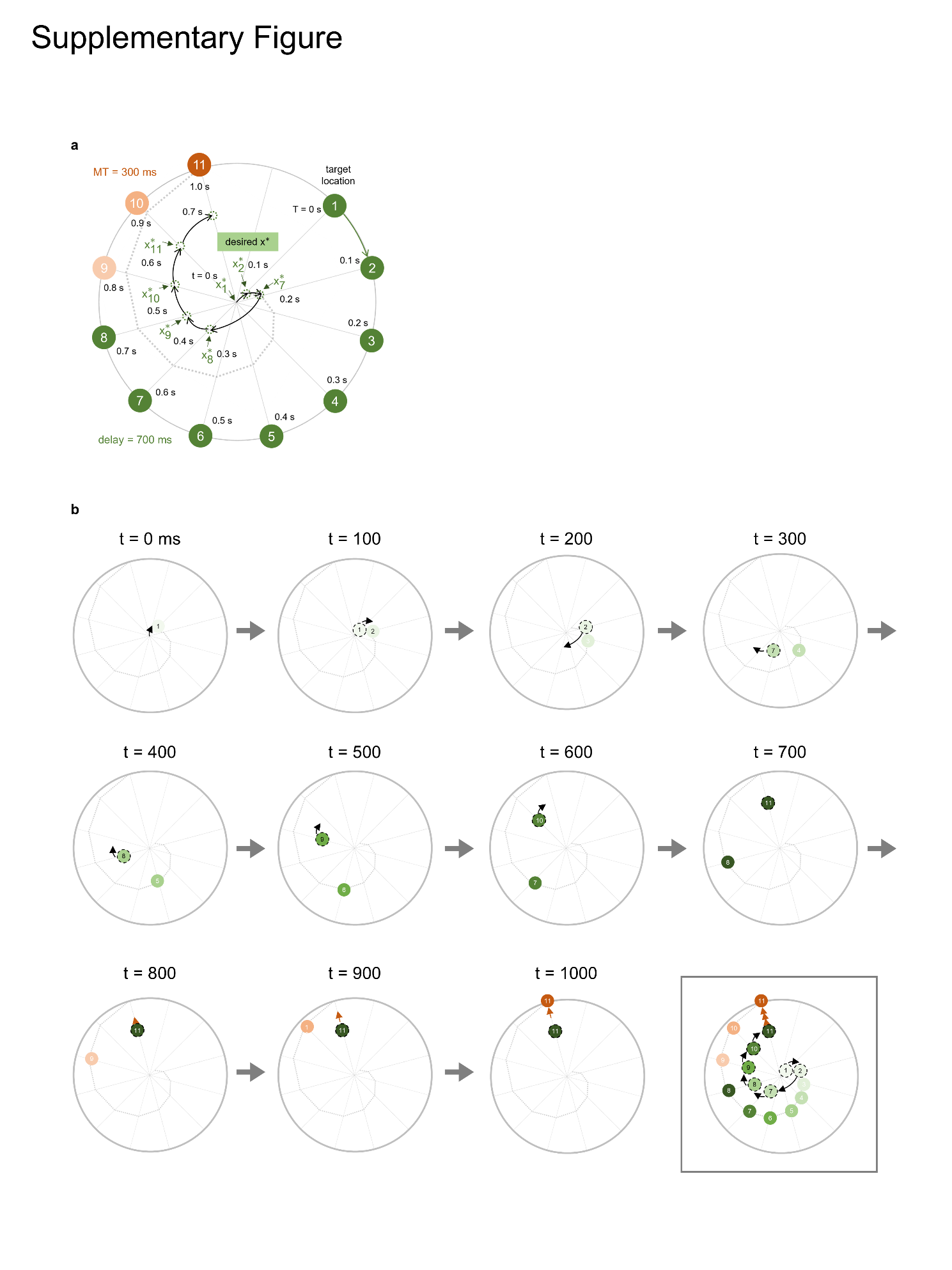

**Supplementary Figure 16. Optimal states of the moving condition in model.**

**a.** Schematic illustrating the optimal states of the moving condition in model. Twelve optimal states (dotted-line circle) correspond to 12 reach directions (filled circle). The target moves at a velocity of 300°/s in a clockwise direction. After target onset (T = 0 s), the desired state $x_{k}^{*}$ is assigned for the network to achieve, indicated by solid green arrows that show when and what the state is allocated. The motor intention precedes the actual target position by 400 ms and is updated every 100 ms until movement onset (T = 0.7 s). After that, preparatory control input u is withdrawn to generate movement with a movement time (MT) of 300 ms.

**b.** A step-by-step demonstration of panel **a**. Each sub-panel depicts the target location (filled circles on the gray trajectory) and the desired states (filled circles with dotted edges) required for the network to evolve every 100 ms. The final panel illustrates the entire process.

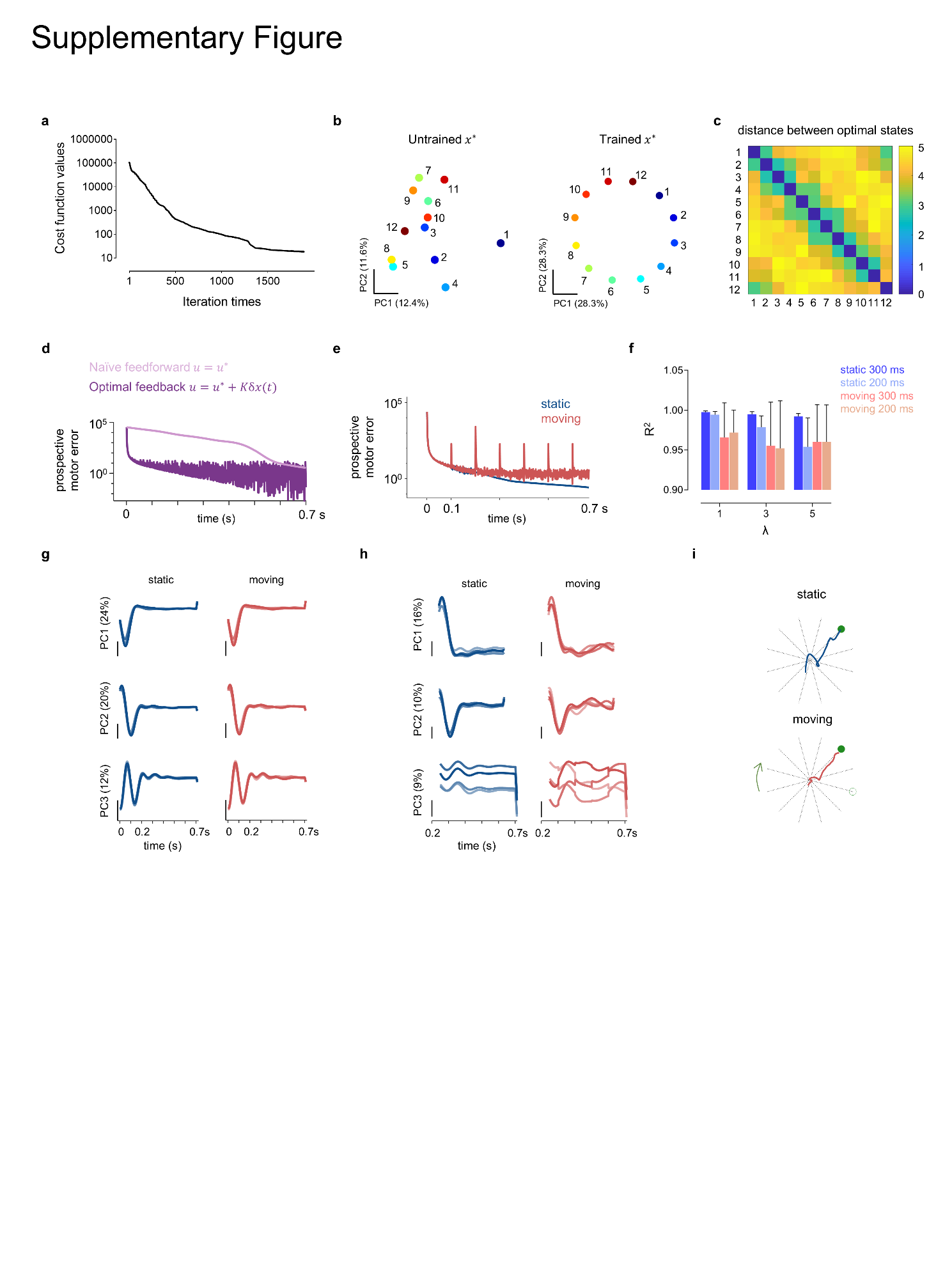

**Supplementary Figure 17. Features of the input-driven model.**

**a.** Cost function value during the optimization of readout matrix $C$ and initial conditions $x^{*}$ at each iteration.

**b.** The first two principal components of neural states before (right) and after (left) optimization. Neural activity with 200 dimensions were processed by PCA. The sequence of states aligns with the order of reach directions.

**c.** Euclidean distances between neural states of all dimensions (n = 200) as illustrated in panel **b**.

**d.** Prospective motor errors in the static condition when input u is either naïve feedforward or optimal feedback.

**e.** Prospective motor error in the moving condition and the static condition with mixed control strategies. Stars indicate the errors resulting from switches in target states every 100 ms in the moving condition.

**f.** Goodness of fit (*R^2^*) of model with varying λ values and epochs length with optimal feedback input. Results are repeated 10 times with different initial states.

**g.** The first three principal components of neural states (TO to GO+10 ms, #units = 200) for four reach directions indicated in **Fig. 6c**.

**h.** Neural states in panel **g** during the period from TO+250 ms to GO+10 ms.

**i.** Subsequent reach trajectories for one representative simulation shown in **Fig. 6e**.

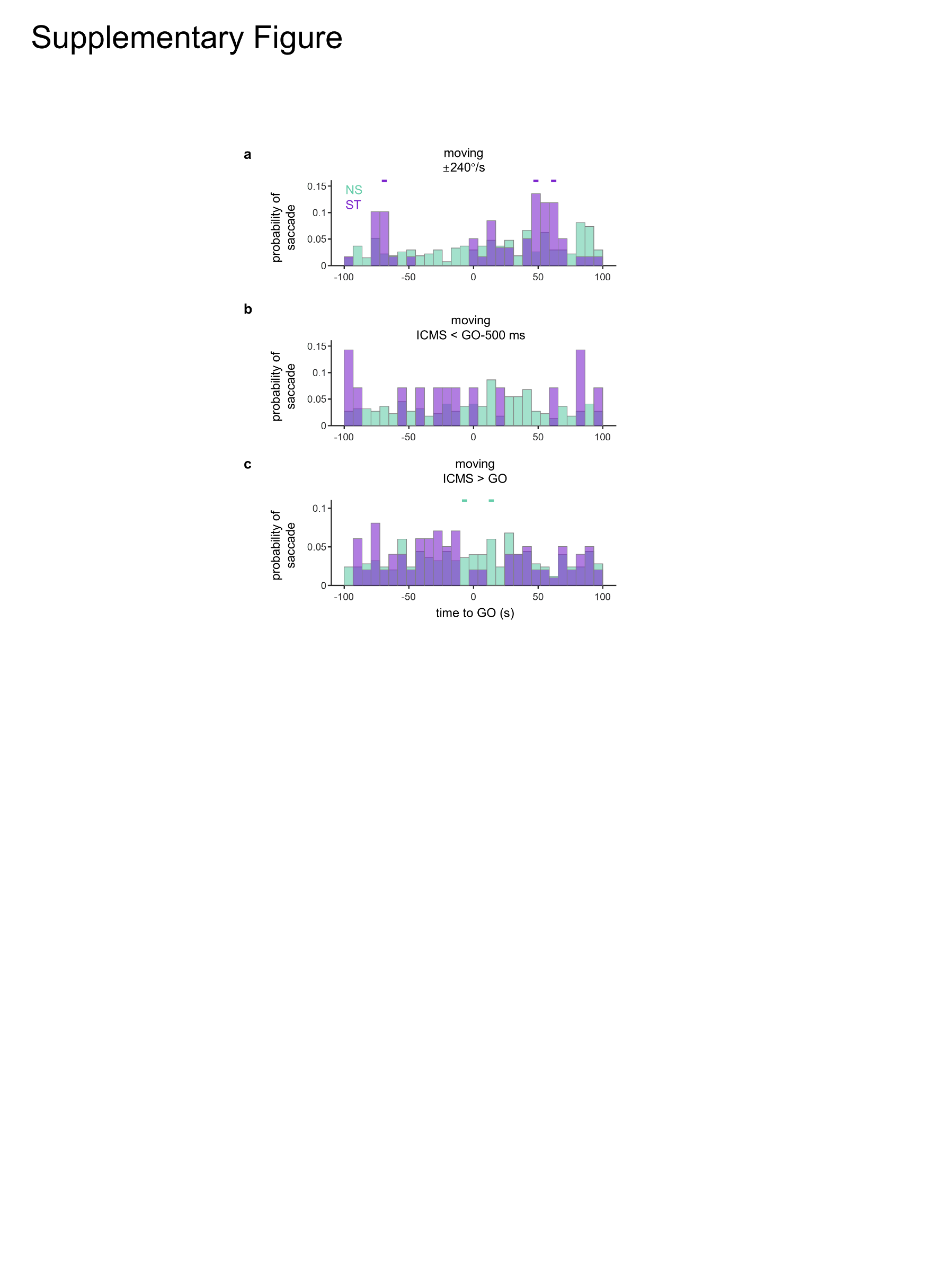

**Supplementary Figure 18. Control results for the probability of saccade.**

**a.** Comparison of saccade probability between NS and ST trials in the moving condition with a target velocity of ±240°/s, where ICMS was applied from GO-100ms to GO. The same annotations apply to panels **b** and **c**.

**b.** Comparison of saccade probability between NS and ST trials in the moving condition with a target velocity of ±120°/s, where ICMS was applied earlier than GO-500 ms.

**c.** Comparison of saccade probability between NS and ST trials in the moving condition with a target velocity of ±120°/s, where ICMS was applied later than GO.
